# CITE-seq reveals inhibition of NF-κB pathway in B cells from vitamin D-treated multiple sclerosis patients

**DOI:** 10.1101/2023.09.25.559400

**Authors:** Manon Galoppin, Manon Rival, Anaïs Louis, Saniya Kari, Sasha Soldati, Britta Engelhardt, Anne Astier, Philippe Marin, Eric Thouvenot

**Affiliations:** IGF, University of Montpellier, CNRS, INSERM, Montpellier, France; Department of Neurology, Nîmes University Hospital, University of Montpellier, Nîmes, France; MGX-Montpellier GenomiX, University of Montpellier, CNRS, INSERM, Montpellier, France; Toulouse Institute for Infectious and Inflammatory Diseases (Infinity), University of Toulouse, CNRS, INSERM, France; Theodor Kocher Institute (TKI), University of Bern, Switzerland

**Author notes:** Corresponding author: Prof. Éric Thouvenot Service de Neurologie, Hôpital Carémeau 4 place du Professeur Robert Debré 30029 Nîmes cedex 9, France.

## Abstract

Vitamin D deficiency is a recognized risk factor for multiple sclerosis (MS) and has been associated with disease activity and progression. Vitamin D treatment has emerged as potentially protective, despite conflicting results from randomized controlled trials. Here, we used single-cell RNA-sequencing (scRNA-seq) combined with barcoded antibodies targeting surface markers (CITE-seq) to uncover candidate genes and pathways regulated in PBMC subpopulations from MS patients receiving high-dose vitamin D (n=5) or placebo (n=5). Best candidates were combined with genes involved in immune function and vitamin D metabolism for validation in a new cohort (n=8 in each group) by high-throughput quantitative polymerase chain reaction (HT-qPCR) in FACS-sorted naive CD4, Th1, Th17, Treg, naive CD8, memory and naive B cells, and MAIT cells. CITE-seq revealed no significant changes in the proportions of these subpopulations in response to vitamin D treatment. Out of the 92 candidate genes identified by CITE-seq, we validated differential expression of five genes (UXT, SNRPN, SUB1, GNLY and KLF6) using HT-qPCR. Furthermore, CITE-seq uncovered vitamin D-induced regulation of several pathways in naive and memory B cells, including MAPK, TLR and interleukin pathways, that may contribute to counteract Epstein-Barr virus (EBV)-induced resistance to apoptosis, notably through inhibition of the NF-κB pathway.

**Graphical Abstract:** 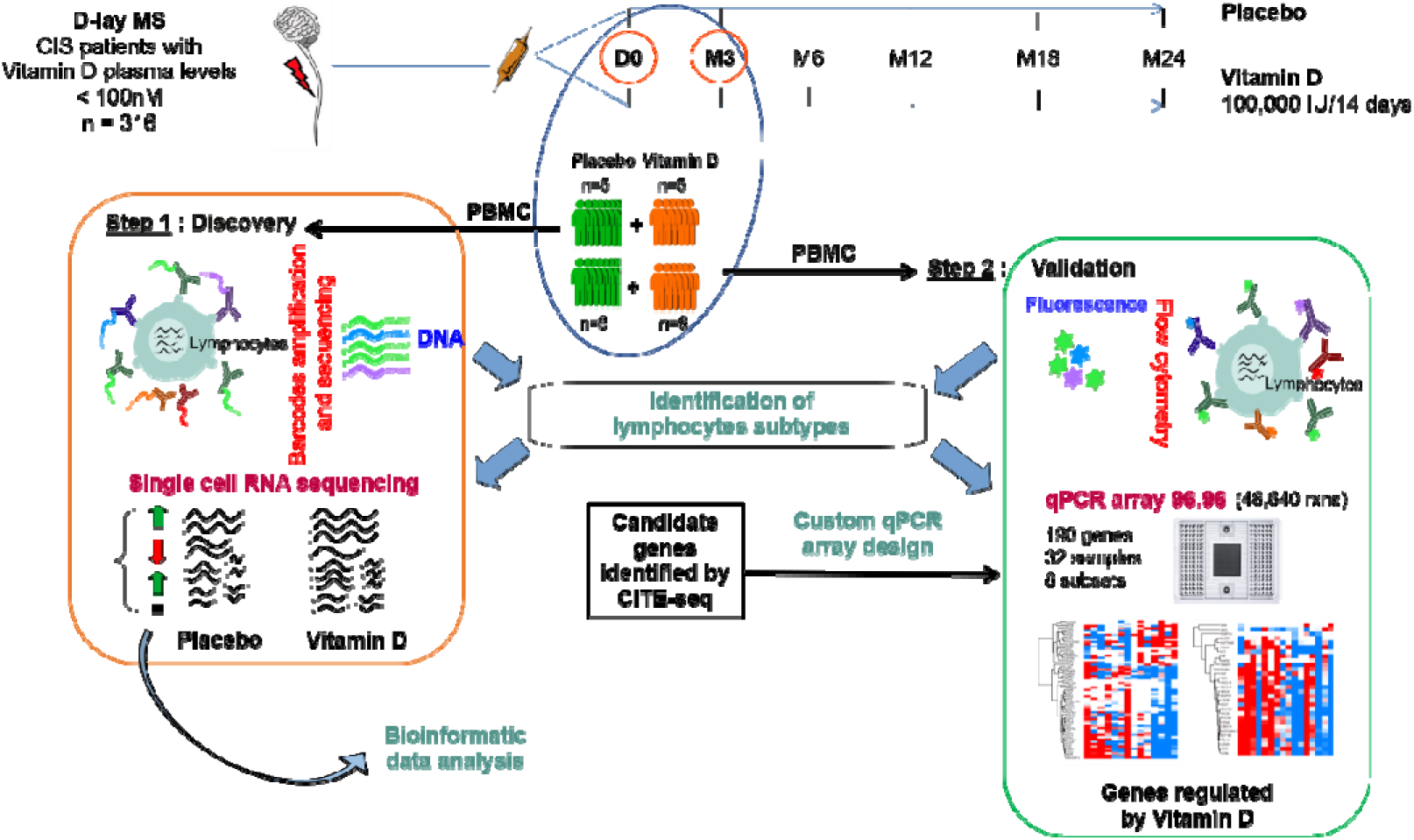

## Introduction

Multiple sclerosis (MS) is a chronic inflammatory, neurodegenerative and autoimmune disease that primarily affects young women. Epidemiological studies have revealed genetic risk factors such as major histocompatibility complex (MHC) loci, especially HLA DRB1*15:01. Additionally, genome-wide association studies (GWAS) have identified over 200 susceptibility variants unrelated to MHC (1). These variants are part of multiple signaling pathways involved in both innate and adaptive immune responses, distributed across immune cells and glial cells, highlighting the potential influence of altered responses in peripheral immune cells and brain-resident cells in the development of MS.

Various environmental factors including smoking, obesity, infection with the Epstein-Barr Virus (EBV), sun exposure, and vitamin D deficiency (2), also contribute to the risk for developing MS. Circulating levels of Vitamin D are influenced by specific genetic variants, skin color and sun exposure, especially during early life. Numerous studies have established a clinical correlation between circulating vitamin D levels and MS disease activity, as well as disability progression (3). Vitamin D binds to the vitamin D receptor (VDR)-retinoid x receptor (RXR) heteromer and the resulting complex translocates to the nucleus and binds to vitamin D response elements (VDREs) in the genomic DNA to modulate the expression of more than 200 genes (4). Most of them are related to vitamin D metabolism and immune function, highlighting the potential role of vitamin D deficiency in MS and other inflammatory diseases. Although a few immunological effects of vitamin D have been reported *in vivo,* studies have demonstrated pleiotropic effects of vitamin D in animal models of MS (EAE) or *in vitro* and suggested the therapeutic potential of vitamin D in MS patients (5). However, therapeutic trials evaluating the clinical and radiological benefits of vitamin D treatment in MS patients provided conflicting results, potentially because the effect of vitamin D was evaluated in conjunction with MS immunomodulators (6–8).

Transcriptomic studies allow unbiased characterization of immune system regulation and can help understanding MS pathophysiology and identifying new potential therapeutic targets for disease-modifying drugs (DMDs) (9). In-depth analysis of the impact of vitamin D in MS patients might reveal gene under its control in different immune cell subtypes *in vivo* and its impact on the physiopathology of MS. A recent transcriptomics study evaluated the effects of vitamin D on gene expression in only three major immune cell types (CD4^+^ T-cells, CD19^+^ B-cells, CD14^+^ monocytes) from MS patients and identified differences in gene expression (10).

In the present study, we compared gene expression in peripheral lymphocytes from early MS patients, naive of DMDs and treated with high-dose vitamin D (100,000 IU every 2 weeks for three months) or placebo as part of a randomized clinical trial (NCT01817166). We used single-cell RNA-seq (scRNA-seq) coupled with barcoded antibodies targeting surface markers (CITE-seq) to discover candidate genes and pathways regulated in various PBMC subpopulations. Best candidates were combined with genes involved in immune function and vitamin D metabolism for validation in a new cohort by high-throughput qPCR (HT-qPCR) in naive CD4, Th1, Th17, Treg, naive CD8, memory and naive B cells, and MAIT cells.

## Results

### Characteristics of the cohorts

Demographic and baseline characteristics of MS patients included in the discovery (n=5 patients per group) and validation (n=8) cohorts and treated with placebo or vitamin D are depicted in Table 1. At inclusion (D0), differences in sex, age and for vitamin D plasma levels were not significant between placebo and vitamin D groups in the discovery (*p*=1, *p*=0.88 and *p*=0.84, respectively) and validation cohorts (*p*=0.20, *p*=0.46 and *p*=0.22, respectively). In addition, the delay since the last corticosteroid infusion was similar in both groups (*p*=0.60). After three months of treatment (M3), plasma levels of vitamin D were significantly different between placebo and vitamin D group (*p*=0.0079 and p=0.0002 in the discovery and the validation cohorts, respectively).

**Table 1.** Demographic characteristics of MS patients included in discovery and validation cohorts.

|  | Discovery |  |  | Validation |  |  |
| --- | --- | --- | --- | --- | --- | --- |
|  | Placebo group (n = 5) | Vitamin D group (n = 5) | p-value between groups | Placebo group (n = 8) | Vitamin D group (n = 8) | p-value between groups |
| Age, y, mean (SD) | 35.8 (10.3) | 34.6 (7.70) | 0.881 | 40.6 (8.62) | 37.4 (8.83) | 0.458 |
| Women, % | 80.0 | 80.0 | > 0.9999 | 87.5 | 62.5 | 0.200 |
| Conversion rate (%) | 100 | 100 | > 0.9999 | 62.5 | 37.5 | 0.619 |
| Days from last steroid infusion (SD) | 35.7 (8.08) | 42.7 (20.5) | 0.600 |  |  |  |
| Serum 25(OH)D ng/mL, mean (SD) D0 | 56.8 (29.2) | 58.4 (14.0) | 0.841 | 60.4 (22.6) | 46.7 (14.3) | 0.220 |
| Serum 25(OH)D ng/mL, mean (SD) M3 | 52.6 (26.6) | 155 (21.1) | 0.0079 | 53.8 (18.0) | 143.3 (17.5) | 0.0002 |
| p-value serum 25(OH)D D0 vs. M3 | > 0.9999 | 0.0079 |  | 0.631 | 0.0002 |  |

### Analysis of lymphocyte subpopulations using CITE-seq

To decipher the effects of vitamin D on the immune balance, we used CITE-seq to describe the evolution of the proportion of each lymphocyte subset in PBMCs from MS patients before and after the 3-month placebo or vitamin D treatment (four patient groups). The expression of specific genes and surface markers was used to annotate major PBMC populations (Supplemental Figure 1), including CD4^+^ T-cells, CD8^+^ T-cells, B-cells (CD19^+^), natural killer (NK) cells (CD56^+^), natural killer T-cells (NKT; CD3^+^CD56^+^), MAIT cells (CD161^+^), monocytes, platelets, and dendritic cells (DCs). CD4^+^ subsets were further annotated with a combination of surface markers including CD45RA^+^ CCR7^+^ (naive), CXCR3^+^CCR6^-^TBX21^+^(Th1), CXCR3^-^CCR6^-^CCR4^+^GATA3^+^ (Th2), CXCR3^-^CCR6^+^RORC^+^ (Th17), CD45RA^-^CXCR5^+^ICOS (Tfh) and CD45RA^-^CD25^+^FOXP3^+^ (Treg). CD8^+^ cell subsets were annotated with CD45RA^+^ CCR7^+^ (naive), CD45RA^-^ CCR7^+^ (central memory, CM), CD45RA^-^CCR7^-^ (effector memory, EM), CD45RA^+^CCR7^-^ (effector, EMRA^+^), and γδTCR*^+^* (γδ T-cells). B cell subsets were annotated using CD27^-^IgD^+^ (naive), CD27^+^IgD^+^ (transitional) and CD27^+^IgD^-^ (memory). UMAPs showed similar profiles in the four groups studied (Figure 2A).

**Figure 1.**
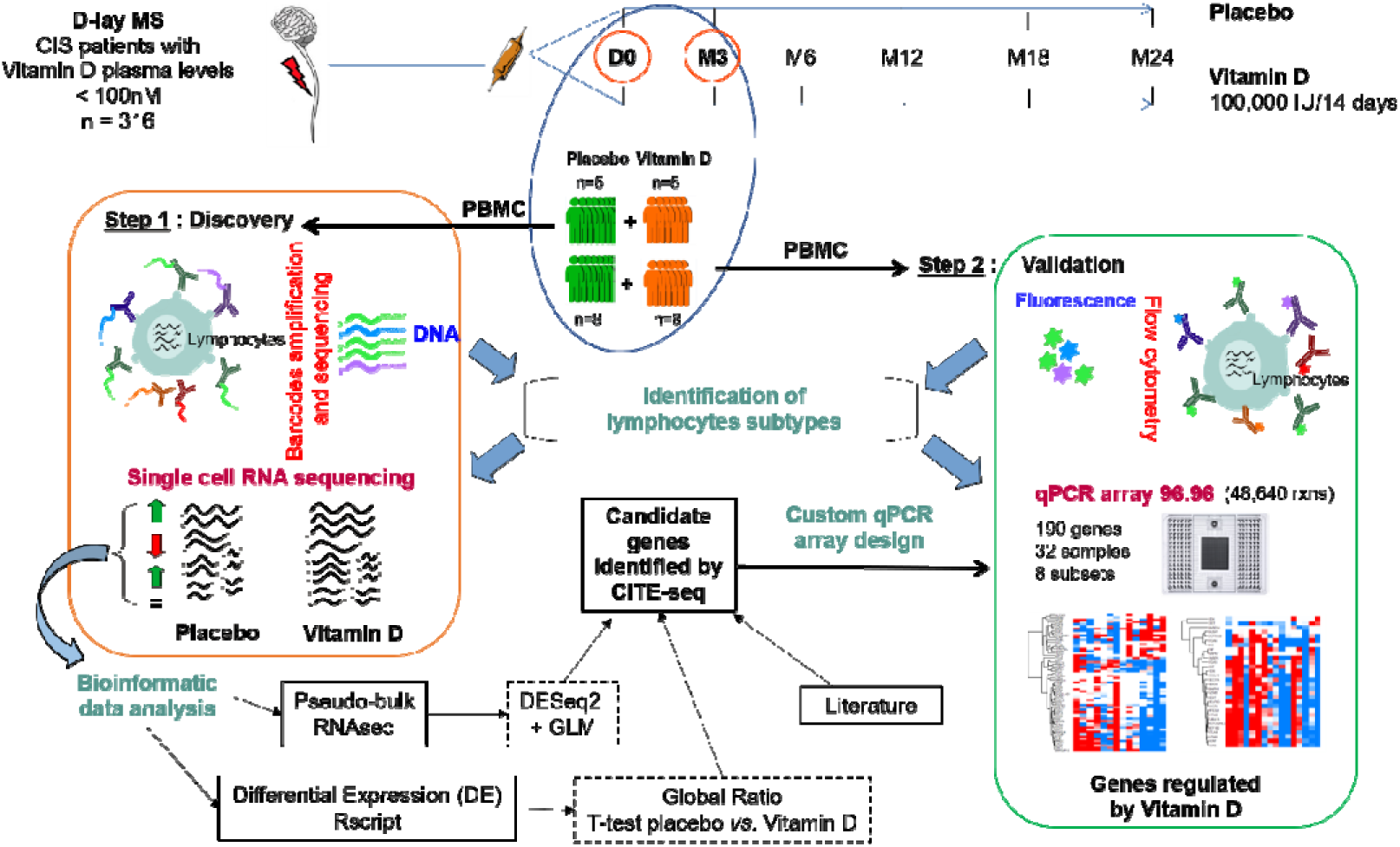
Overview of the study design and analysis method.

**Figure 2.**
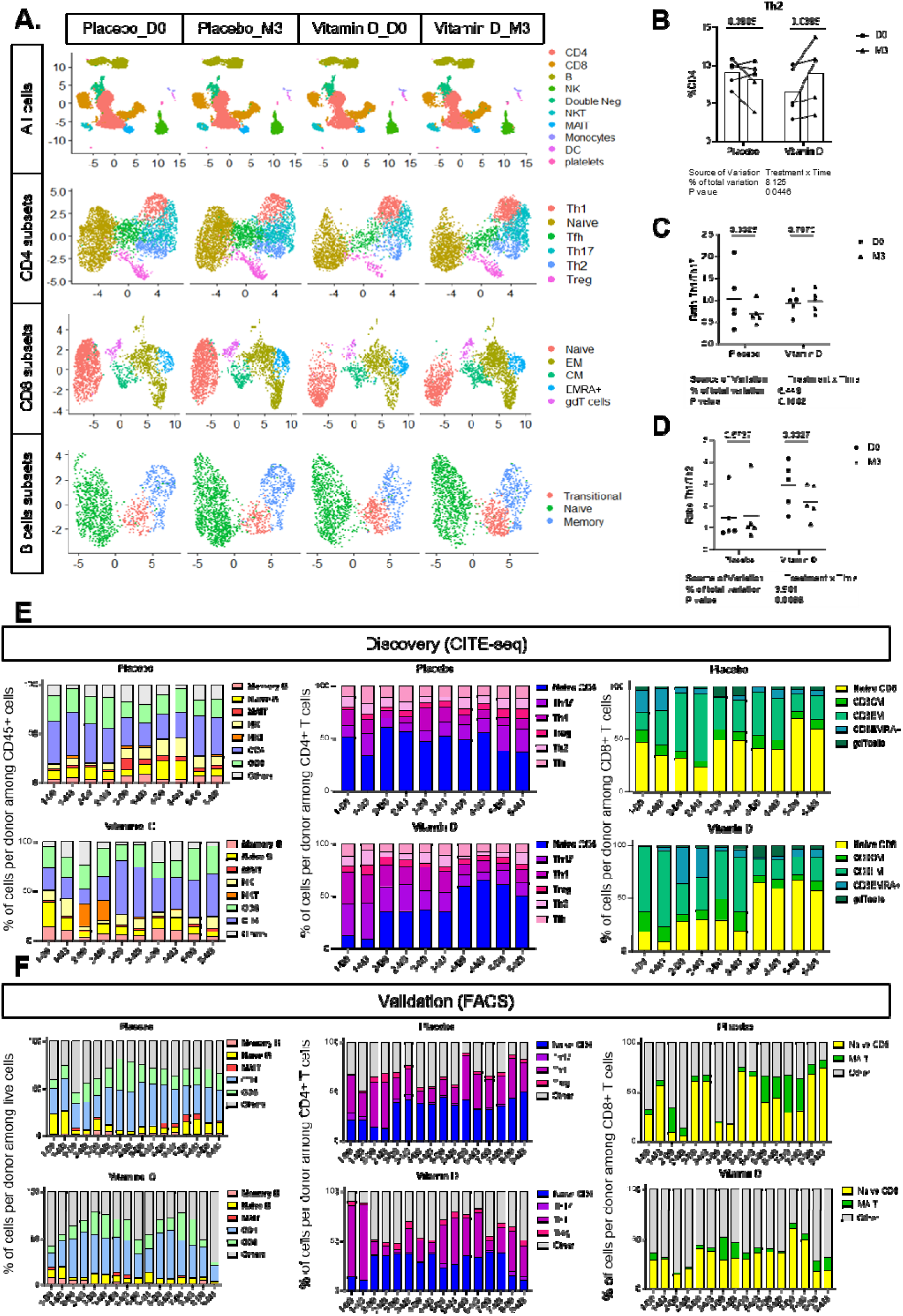
Char acterization of circulating lymphocytes subsets in PBM Cs from MS patie nts using CIT E-seq. (A) UMAP and clustering of different lymphocytes subsets of PBMCs. NK, natural killer; NKT, natural killer T-cells; MAIT, mucosal-associated invariant T-cells; EM, effector memory; CM, central memory, EMRA+, terminally differentiated effector memory cells re-expressing CD45RA. (B) Change in the proportion of Th2 cells before/after treatment in the placebo and vitamin D groups. (C and D) The Th1/Th17 and Th1/Th2 cell ratio were expressed as the ratio of the percentage of Th1 and Th17 or Th2 subsets in each patient. (E) Proportion of each lymphocyte subset before and after Vitamin D or placebo treatment in the discovery step. (F) Proportion of each lymphocyte subset before and after Vitamin D or placebo treatment in the validation step. Significance was determined by 2way-ANOVA (B, C, D) with original FDR BH correction.

For each cluster, we extracted a list of cells, counted and normalized them as a percentage, and compared the proportion of each lymphocyte subset before and after Vitamin D or placebo treatment (Figure 2E). In all groups, we observed an inter-individual variability in the distribution of the various cell subsets. However, for each patient, no significant changes were seen in the proportions of the different lymphocyte subpopulations after the 3-month treatment (Supplemental Figure 2), except for Th2, which was increased after vitamin D treatment (2-way ANOVA, *p*=0.0446, Figure 2B).

To further explore the regulation of lymphocyte populations by vitamin D, we analyzed changes in main subset ratios. The Th1/Th17 ratio showed no significant difference between the groups. A non-significant decrease (p=0.092) was measured after the treatment in the placebo group but not in the vitamin D group (*p*=0.79, Figure 2C). On the other hand, the Th1/Th2 ratio was significantly decreased in the vitamin D group (*p*=0.0027), but not in the placebo group (*p*=0.57, Figure 2D), consistent with the increase of Th2 subset in the vitamin D group (Figure 2B).

### Modulation of gene expression

To identify genes affected by vitamin D treatment in MS patients, we looked for differentially expressed genes (DEGs) in vitamin D and placebo-treated patients using two different methods. We first analyzed the global treatment effect ratio ((M3/D0 Vitamin D) / (M3/D0 placebo)) for each lymphocyte subset. This revealed 16 DEGs in memory B-cells (9 up-regulated genes and 7 down-regulated genes; *p* < 0.05), 19 DEGs in naive B-cells, (8 up, 11 down) (Figure 3A). *JUN* was the only common DEG between these two B-cell types (Supplemental Table 3). Among CD4^+^ T-cells, 21 DEGs were identified in naive CD4^+^ (2 up, 19 down), 77 in Th17 (72 up, 5 down), 13 in Th1 (7 up, 6 down), and 27 in Treg (14 up, 13 down). Two of them were shared by 2 subsets: *SNHG14* in Th1 and Th17 cells and *EEF1B2* in Th17 and Treg. In naive CD8^+^ T-cells, 20 DEGs (17 up, 3 down) were identified. Finally, 50 DEGs (47 up, 3 down) were identified in MAIT cells.

**Figure 3.**
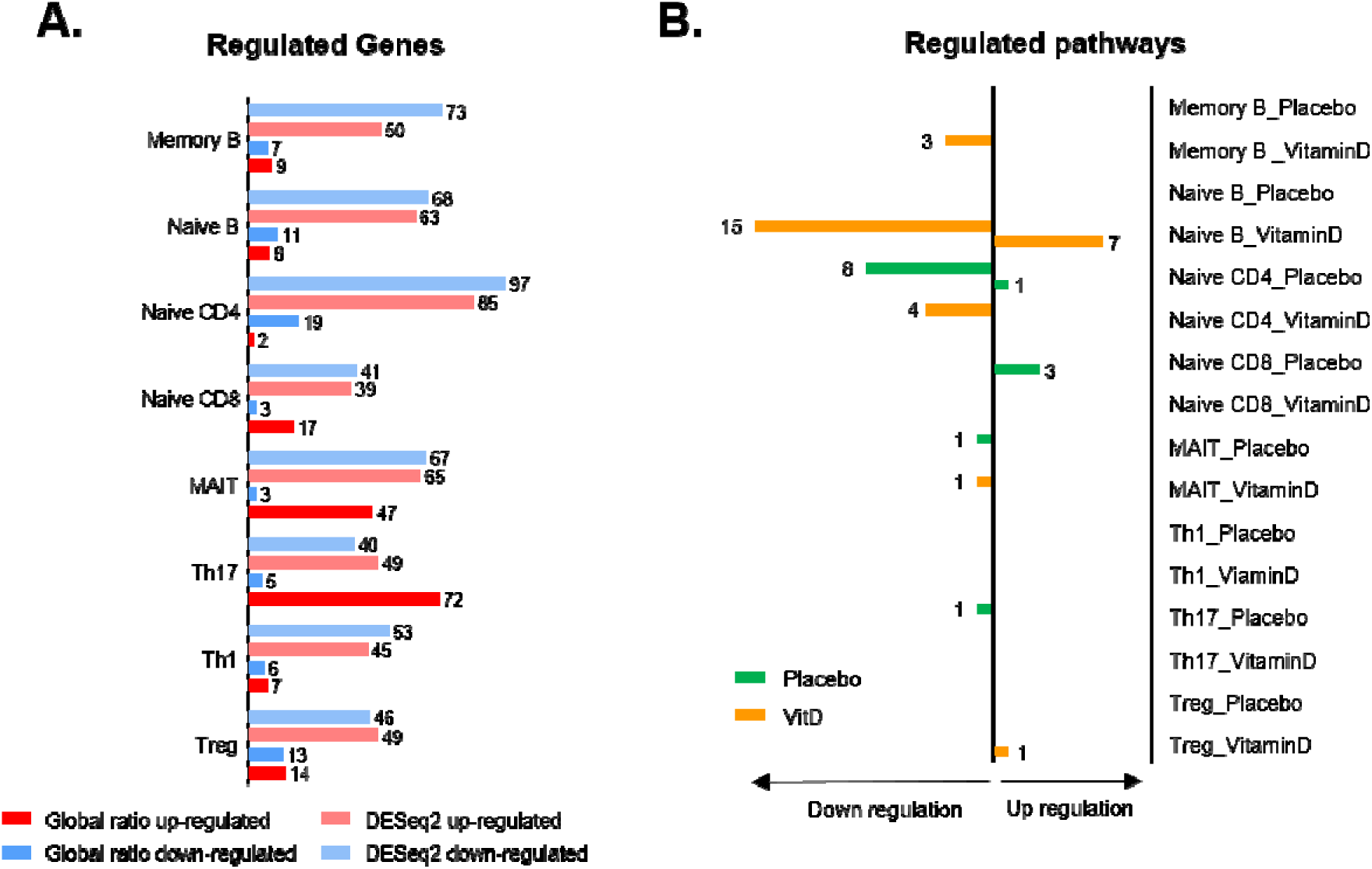
Regulation of gene expression and pathways by 3-months placebo or vitamin D treatment. (A) Number of differentially expressed genes (DEGs, p<0.05) in lymphocytes subsets according to the method used for analysis (global ratio or DESeq2 analysis). (B) Number of significantly overrepresented and underrepresented pathways (p<0.05) from the Reactome Pathway Database in lymphocytes subsets assessed by Single Cell Pathway Analysis (SCPA) Package from RStudio.

Among all DEGs identified with this first approach, some were found in several lymphocyte populations: *JUN* (memory B-cells, naive B-cells, naive CD4^+^), *C9orf78* (memory B-cells, MAIT), *HNRNPDL* (memory B-cells, naive CD4^+^, naive CD8^+^), *PSMB3* (memory B-cells, Th17, Treg), *UXT* (memory B-cells, Th17), *B2M* (naive B-cells, Treg), *BOD1L1* (naive B-cells, MAIT), *GCC2* (naive B-cells, naive CD4^+^), *TUBA1A* (naive B-cells, naive CD8^+^), *IKZF1* (naive CD4^+^, naive CD8^+^, MAIT), *LAMTOR4* (naive CD4^+^, naive CD8^+^), *NDUFA1* (naive CD4^+^, MAIT), *EID1* (naive CD8^+^, Th17), *RICTOR* (naive CD8^+^, Th1), *YPEL3* (naive CD8^+^, Treg), *ENO1*, *HLA-E*, *MTDH*, *PDIA3*, (MAIT, Th17), *KIF2A*, *PABPC1* (MAIT, Th1), *SNRPN* (MAIT, Treg) (Supplemental Table 3).

We then performed DESeq2 analysis, which identified 123 DEGs (p-value < 0.05) in memory B-cells (50 up-regulated, 73 down-regulated), 131 in naive B-cells (63 up, 68 down), 182 in naive CD4^+^ (85 up, 97 down), 80 in naive CD8^+^ (39 up, 41 down), 132 in MAIT cells (65 up, 67 down), 89 in Th17 (49 up, 40 down), 98 in Th1 (45 up, 53 down), and 95 in Treg (49 up, 46 down). Among all DEGs identified with this second approach, some were found in several lymphocyte populations (Supplemental Table 3 and Supplemental Figure 3).

Five DEGs were identified in both analyses in memory B-cells (*S100A6, S100A4, IRF8, ZFP36, PSMB3*), 9 in naive B-cells (*IGCL2, B2M, TUBA1A, KLF2, PSME1, JUN, CAST, DUSP1, SET*), 2 in naive CD4^+^ (*CDC42SE2, JUN*), 5 in naive CD8^+^ (*TUBA1A, RNF213, SP100, PABPC4, CLEC2D*), 4 in MAIT cells (*NDUFA1, NDUFA11, SAT1, SUB1*) and 3 in Th17 (*CYTIP, IFITM2, NUCKS1*). No common genes were found in Th1 and Treg subsets (Supplemental Table 3).

Among all DEGs, we selected those exhibiting the strongest modulation (>1.2-fold change) upon vitamin D treatment. These include 11 DEGs in memory B-cells, 12 in naive B-cells, 11 in naive CD4^+^, 10 in naive CD8^+^, 15 in MAIT and Th17, 13 in Th1 and 17 in Treg were selected for the qPCR validation step (Supplemental Table 3).

### Analysis of pathways regulated by vitamin D

We next investigated which pathways are over-represented after 3 months of treatment by Vitamin D (or placebo) using Single Cell Pathway Analysis (SCPA) Package from CITE-seq data and the Reactome Pathway Database.

After three months of placebo treatment, subtle changes were observed: one pathway was over-represented and eight were under-represented in naive CD4^+^, three were over-represented in naive CD8^+^, one was under-represented in MAIT and one was under-represented in Th17 (Figure 3B and Supplemental Table 4). No significant pathway modulation was seen in memory B-cells, naive B-cells, Th1 and Treg. The three-months vitamin D treatment induced more robust changes: three pathways were significantly under-represented in memory B-cells, seven were over-represented and 15 were under-represented in naive B-cells, four were under-represented in naive CD4^+^ cells, one was under-represented in MAIT and one was over-represented in Treg (Figure 3B and Supplemental Table 4). We observed no significant pathway modulation in naive CD8^+^, Th1 and Th17. Naive B-cell subsets had the larger number of modulated pathways after vitamin D treatment, including several mitogen-activated protein kinase (MAPK) pathways (MAPK6 MAPK4 SIGNALING, RAF INDEPENDENT MAPK 1,3 ACTIVATION, MAPK TARGETS NUCLEAR EVENTS MEDIATED BY MAP KINASES, MAPK FAMILY SIGNALING CASCADES, NEGATIVE REGULATION OF MAPK PATHWAY) (e.g., *JUN, FOS, DUSP1, PI3KCA, PPP2CB,* Supplemental Table 4). In addition, naive B-cells showed modulation of TOLL LIKE RECEPTOR 9 TLR9 CASCADE, TOLL LIKE RECEPTOR TLR1 TLR2 CASCADE, MYD88 INDEPENDENT TLR4 CASCADE, and INTERLEUKIN 1 SIGNALING (e.g., *JUN, FOS, PPP2CB, PSMD2, USP14*), four pathways activated by microbial components including LPS or CpG and involved in IL-1 signaling in lymphocytes. The TRAF6 MEDIATED NF KB ACTIVATION and INTERLEUKIN 1 FAMILY SIGNALING pathways were regulated in memory B-cells (e.g., *TRAF6, PSMB4, PSMD7*).

Moreover, DEFECTIVE INTRINSIC PATHWAY FOR APOPTOSIS was under-represented in naive B-cells, indicating an increased apoptosis in these cells, while SIGNALING BY ROBO RECEPTORS was under-represented in naive CD4 T-cells, indicating a reduction of migration capacity of these cells.

Finally, TOLL LIKE RECEPTOR 9 TLR9 CASCADE and MAPK TARGETS NUCLEAR EVENTS MEDIATED BY MAP KINASES pathways were modulated in both naive B-cells and naive CD4^+^ T-cells, while TOLL LIKE RECEPTOR TLR1 TLR2 CASCADE was modulated in both naive B-cells and MAIT cells (Supplemental Table 4).

### Validation of candidate genes by high throughput qPCR

In order to enlarge the number of genes selected for validation by HT-qPCR, we added 92 genes encoding for cytokines/chemokines and their receptors, as well as proteins involved in adhesion, proliferation, activation, migration, and vitamin D metabolism (Supplemental Table 2)to the DEGs identified in the CITE-seq-based discovery step, leading to a list of 178 genes that were analyzed by HT-qPCR in a new cohort of MS patients (Table 1).

PBMC samples from this cohort were sorted by FACS to select 8 subpopulations of interest (memory B-cells, naive B-cells, naive CD4, naive CD8, Th1, Th17, Treg, MAIT cells). As for the CITE-seq analysis, we observed an important inter-individual variability in the proportions of these lymphocyte subpopulations at baseline (Figure 2F). No significant changes were seen in the proportion of each lymphocyte subpopulation after three months of placebo or vitamin D treatment (Figure 2F).

Comparing placebo and vitamin D patients after three months of treatment identified 4 DEGs in memory B-cells and naive B-cells, 5 DEGs in naive CD4^+^, 3 DEGs in MAIT cells, 8 DEGs in Treg, 23 DEGs in Th17 and 38 DEGs in Th1 (Figure 4). Some of them were previously identified by the global ratio method in the same population (UXT in memory B-cells and SNRPN in MAIT cells), or the DESeq2 method (ITGA5 in MAIT cells and GNLY in Th17) or both methods (SUB1 in MAIT cells, NEDD4L and MIER2 in Th17, and KLF6 in Th1). Concordant variations were found in the discovery and validation steps for five of these genes (UXT decrease in memory B-cells, KLF6 decrease in Th1, GNLY decrease in Th17, SNRPN and SUB1 increase in MAIT cells). No candidate gene modulated by vitamin D in naive CD8^+^ was validated by HT-qPCR.

**Figure 4.**
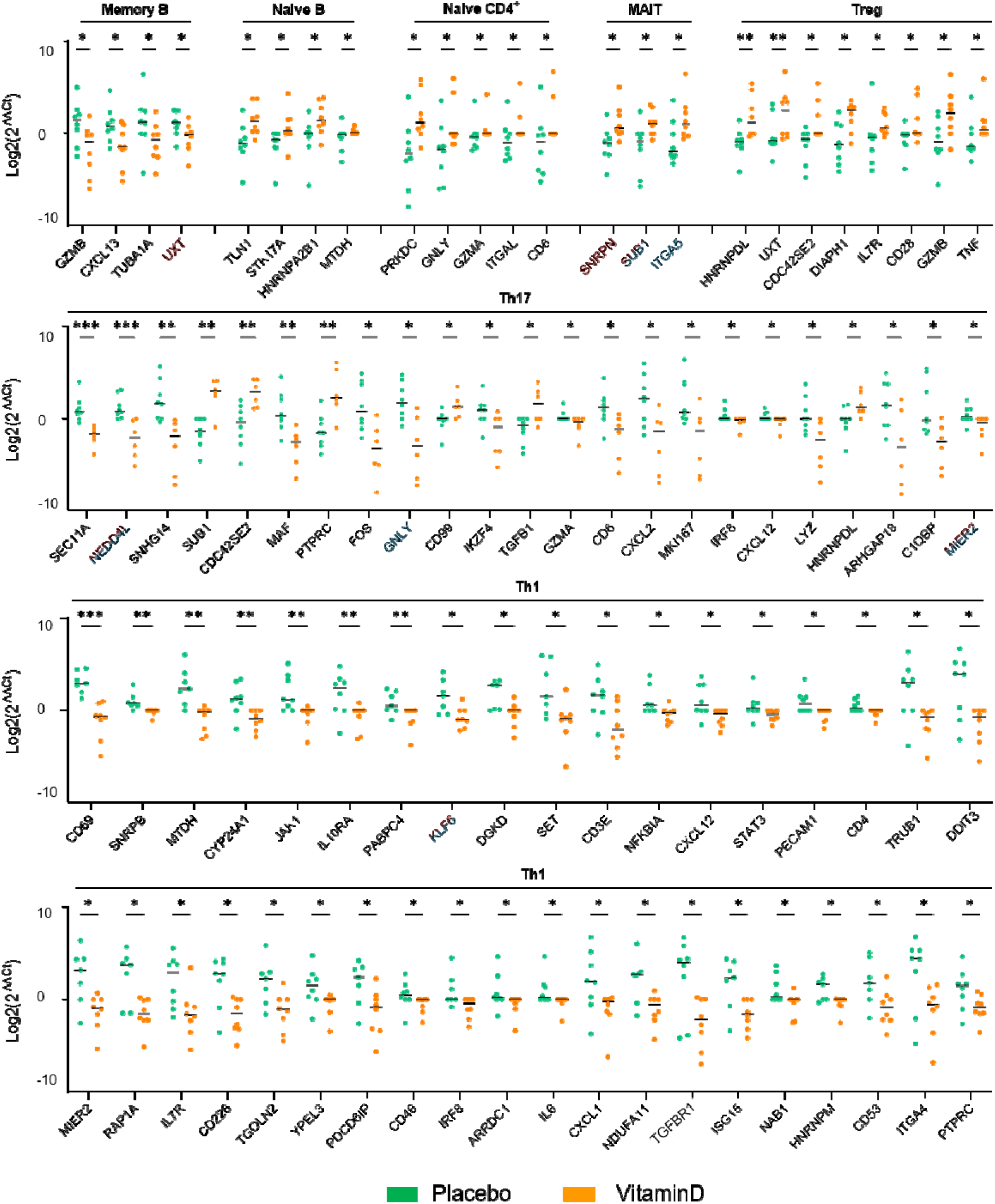
Validation of candidate genes by high-throughput qPCR in lymphocyte subset. Genes identified by the global ratio method are highlighted in red. Genes identified with the DESeq2 method are highlighted in blue. Expression of each gene is plotted as Log2(*2*^−ΔΔCt^). Significance was determined by multiple Mann-Whitney tests, *p < 0.05, **p < 0.01, ***p<0.001.

Intriguingly, some regulated genes involved in identified pathways and selected for qPCR validation showed non-significant ratios by HT-qPCR in naive B-cells (*DUSP1,* p=0.10 and *FOS*, p=0.13) and memory B-cells (*JUN*, p=0.28). When a p-value threshold of 0.20 was considered, an opposite regulation of gene expression by vitamin D was found in naive and memory B-cells: whereas most genes were downregulated in memory B-cells in the vitamin D-treated group, they were up-regulated in naive B-cells (Figure 5).

**Figure 5.**
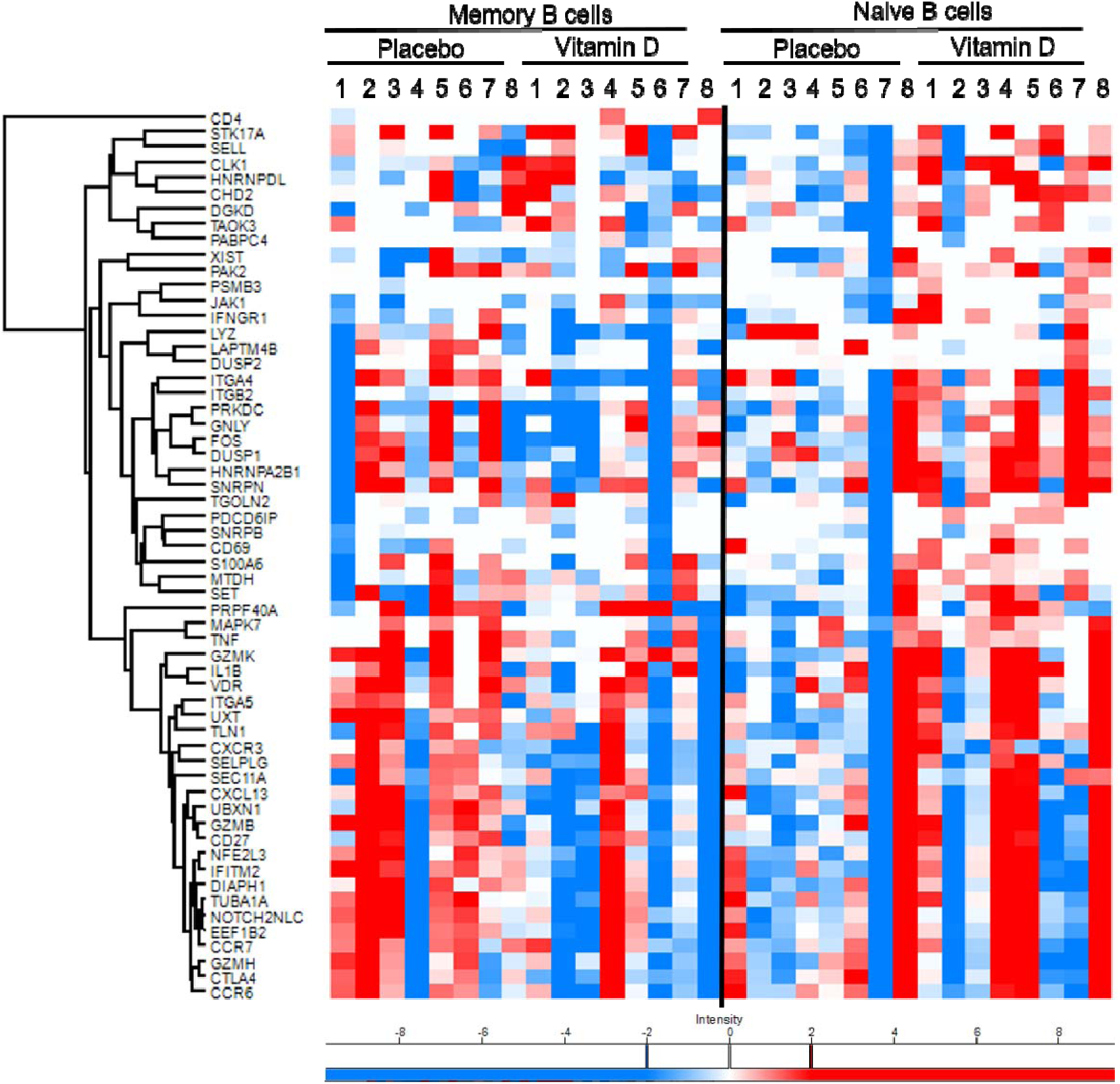
Heat-map of the DEGs (p<0.20) between memory and naive B-cells. For each group (vitamin D and placebo), eight patients (1 to 8) of the validation cohort are represented. Heat-map was performed using Perseus® (Max-Planck-Institute of Biochemistry, Martinsried, Germany, version 2.0.7.0). A gradient coloring scheme was applied to display upregulated (above 2-fold expression (in log2 display above 1, red color) *vs.* downregulated (below 2-fold expression (in log2 display below −1, blue color) genes. Heatmap was hierarchically clustered by Pearson’s correlation distance using row average linkage.

## Discussion

We performed the first CITE-seq analysis of PBMCs collected from MS patients before and after three months of treatment with high-dose vitamin D or placebo, and identified candidate genes and pathways modulated by vitamin D *in vivo*. Compared to previous transcriptomic studies (10), our strategy comparing PBMCs samples from the same individuals at D0 and M3 reduces interindividual variability inherent to clinical samples. Comparing vitamin D and placebo groups highlights gene expression changes actually modulated by vitamin D and not resulting from the recent MS attack and its management with high dose steroid therapy. In addition, single-cell transcriptome analysis allowed a more exhaustive characterization of the different lymphocyte subpopulations affected by vitamin D in MS than previous studies focusing on major lymphocyte populations (CD4^+^ T-cells, B-cells and monocytes). Using 35 different barcoded antibodies targeting specific surface markers, we did not only identify the main lymphocyte populations (CD4^+^ T-cells, CD8^+^ T-cells, B-cells, NK-cells, NKT-cells, MAIT cells), but also several CD4^+^ (naive, Th1, Th2, Th17, Tfh, Tregs), CD8^+^ (naive, CM, EM, EMRA^+^, γδ T-cells) and B-cell (naive and memory) subpopulations. In line with previous studies, vitamin D treatment did not significantly affect the proportion of each cell subset, except for the Th2 population (5). This suggests that vitamin D does not modulate immune cells through important changes in the proportion of pro-or anti-inflammatory cell subsets, but rather through a regulation of the intrinsic properties of these cells (proliferation, adhesion, migration).

Our CITE-seq unsupervised analysis provided a more accurate analysis of gene expression changes in different lymphocyte populations, than classical scRNA-seq performed without barcoded antibodies (11), and is the first to validate candidate genes by HT-qPCR in a different cohort. Using this two-step strategy, we identified and validated five genes: *UXT* in memory B-cells, GNLY in Th17, KLF6 in Th1, and SNRPN and SUB1 in MAIT cells. *UXT* encodes a Ubiquitously Expressed Prefoldin Like Chaperone. Its mRNA undergoes alternative splicing giving rise to multiple isoforms. One of them, UXT-V1, binds to TNF receptor-associated factor 2 and prevents TNF receptor-associated death domain protein from recruiting Fas-associated protein (FADD), and thus protects cells against TNF-induced apoptosis (12). Moreover, Epstein-Barr virus BGLF4 kinase downregulates NF-κB transactivation through engagement of UXT and reduces memory B-cell apoptosis (13). The decreased expression of UXT measured in memory B-cells upon vitamin D treatment suggests that chronic vitamin D treatment might restore B-cell susceptibility to apoptosis and consequently reduces the pro-inflammatory properties of memory B-cells in MS (14,15). Interestingly, we found an opposite regulation of *UXT* in Tregs, suggesting that vitamin D might protect Tregs from apoptosis induction through the stimulation of death receptors and thus prevent Treg deficiency in MS.

*GNLY* encodes granulysin, a protein released in cytotoxic granules by activated lymphocytes that displays cytolytic activity against microbial and tumor cells (16). At micromolar concentrations, granulysin is a chemoattractant for monocytes, CD4^+^ and CD8^+^ memory cells and actively recruits immune cells to inflammation sites (17). The decreased *GNLY* expression observed in Th17 cells from vitamin D-treated patients is consistent with previous findings indicating that vitamin D decreases the percentage of total granulysin-positive cells, including CD4^+^, CD8^+^ and NK-cells in pulmonary tuberculosis patients (18).

*KLF6* encodes Krueppel-like factor 6, a DNA binding protein involved in activation of transcription and post-translational regulatory pathways. KLF family members have been implicated in the differentiation, activation, quiescence, or homing of various T-cell subsets. A previous transcriptomic analysis showed an overexpression of *KLF6* gene in activated T-cells from psoriasis patients (19). The decreased expression of *KLF6* in Th1 after vitamin D treatment found in our study supports the notion that vitamin D can modulate Th1 activation in MS (20).

The *SNRPN*-encoded small nuclear ribonucleoprotein polypeptide N (Sm-D) plays a role in gene expression regulation and in pre-mRNA processing. It is the target of auto-antibodies in systemic lupus erythematosus. Expression of genes encoding Sm proteins was positively correlated with the infiltration of Th2 cells, but negatively correlated with the infiltration of mast cells, Th1 cells and NK cells in a model of lung cancer (21). However, the significance of *SNRPN* expression increase in MAIT cells after vitamin D treatment in the context of MS remains unclear.

*SUB1* encodes positive cofactor 4 (PC4), a transcription factor involved in Tregs activation through a pathway distinct from *FOXP3* (22). Luo *et al.* have shown that *SUB1* overexpression increases *IL10*, *HMGB2*, and *MKI67* mRNA levels in Tregs (22). In diffuse large B-cell lymphoma, PC4 is overexpressed and acts as an upstream regulator of c-Myc. PC4 is closely related to lymphoma clinical staging and prognosis. PC4 knockdown and the resulting inhibition of c-Myc expression can induce autophagic cell death (23). However, the significance of *SUB1* expression increase in MAIT and Th17 cells observed in our study remains to be explored.

The limited number of vitamin D regulated genes identified in our study compared with previous studies using *in vitro* or *ex vivo* preparations challenged or not with vitamin D might have several reasons. First, gene expression analysis was performed after chronic vitamin D treatment (3 months), which might have precluded the identification of early and transiently regulated genes whereas previous *in vitro* or *ex vivo* studies rely on acute vitamin D treatment (24). Second, vitamin D levels in our patients increased from around 50nM to 150 nM, which differs from main *ex vivo* protocols where cells are cultured in culture media deprived from vitamin D and stimulated by adding various and sometimes higher concentrations of vitamin D (generally 100 to 400nM) (25). These different vitamin D concentration ranges might explain why many genes identified (but not validated) in those studies, such as genes encoding cytokines and chemokines, were not found in our CITE-seq analysis. To further explore the influence of vitamin D on the expression of these genes in MS patients, we decided to add to the DEGs identified by CITE-seq a set of genes potentially involved in the pathophysiology of MS, for validation by HT-qPCR. Hence, we observed a decreased expression of *CXCL13* transcript in memory B-cells after vitamin D treatment. CXCL13 is a chemokine that attracts B lymphocytes and its expression is increased in the CSF of RRMS patients and reflects the presence of B-cells (26). Our results suggest that vitamin D could reduce memory B-cell chemo-attraction in the CNS, thereby reducing local inflammation.

Likewise, we found a decreased expression of *IL-6, JAK1* and *STAT3 i*n Th1 cells from vitamin d-treated patients. This contrasts with data from Chauss et al. in healthy donors, showing a paradoxical increase in expression of *IL-6* and *STAT3* transcripts in Th1 cells, that peaks at 48 h post-vitamin D stimulation (20). Nevertheless, our data are consistent with the activation of the IL-6/JAK1/STAT3 pathway described in Th17 from EAE mice (5). CD46 and CD69 were also decreased in our study, consistent with previous data showing a negative correlation between CD46 expression on T-cells and the levels of circulating vitamin D in a cohort of MS patients supplemented by vitamin D (27) and a decreased CD69 expression in active T-cells from Crohn’s disease patients (28). Finally, the decreased expression in Th1 of two genes involved in vascular cell adhesion, namely *ITGA4* that encodes for CD49d (alpha-4 subunit of VLA4, target of natalizumab) and *CD226* might suggest an inhibitory effect of vitamin D on Th1 ability to cross BBB (27). Notably, CD226 is expressed by both Th1 and Th17 cells and anti-CD226 treatment ameliorates the course of EAE (29,30).

Intriguingly, we observed contrasting regulations of gene expression by vitamin D in naive and memory B-cells, two populations with opposite functions (31). Pathway analysis also showed distinct regulations by vitamin D of several MAPK, TLR and interleukin pathways in naive and memory B-cells. In naive B-cells, we found that vitamin D treatment down-regulates interleukin-1 signaling and TLR 1/2 and TLR9 pathways known to stimulate NF-κB signaling (32). NF-κB, MAP kinase, IRF7, and PI3 kinase pathways are known to be stimulated via latent membrane protein 1 (LMP1), a mechanism favoring B-cell immortalization during chronic EBV infection (33). Thus, vitamin D might restore sensitivity to apoptosis of naive B-cells, consistent with the under-representation of the DEFECTIVE INTRINSIC PATHWAY FOR APOPTOSIS. The under-representation of the TRAF6 pathway after vitamin D treatment also suggests a reduction of NF-κB activation in memory B-cells. Collectively, these data suggest that chronic vitamin D treatment contributes to reduction of chronic EBV-induced B-cell activation in MS patients. TLR9 and MAPK pathways are also negatively regulated by vitamin D in naive CD4^+^ T-cells, suggesting similar pro-apoptotic effects of vitamin D in naive B-cells and naive CD4^+^ T-cells. These pro-apoptotic effects might also explain the synergistic effects of vitamin D and glucocorticoid therapy in EAE (34).

## Conclusion

The analysis of gene expression in PBMC sub-populations from MS patients naive from any disease modifying drugs and chronically treated with vitamin D deciphered many subpopulations not investigated by previous transcriptomics studies performed on larger cell populations sorted by FACS (10). It revealed changes in the expression of genes in different lymphocyte subsets that reflect *in vivo* modulation of the immune system by vitamin D in MS patients. It also identified pathways regulated by vitamin D in naive and memory B-cells that potentially prevent EBV-induced resistance to apoptosis through inhibition of the NF-κB pathway, thus highlighting the unexplored regulation by vitamin D of B-cells, a key therapeutic target in MS.

## Material and methods

### Sample of MS patients included in the discovery cohort

The cohort of MS patients is part of the D-lay MS study (NCT01817166), a multi-center, phase III, randomized, double-blind, placebo-controlled hospital-based clinical research protocol in early MS patients naive from DMDs. The primary objective of D-lay MS was to assess the efficacy of cholecalciferol (Vitamin D3) monotherapy in reducing the rate of conversion to active MS after a clinical isolated syndrome (CIS) (according to 2005 McDonald criteria (35)), and its safety over 2 years. Three hundred and sixteen patients, aged between 18 and 55, who had presented a typical CIS less than 90 days ago with the presence of lesions suggestive of MS on brain and spinal cord MRI (36) and low plasma vitamin D levels (< 100 nmol/L) were included and randomized to 100,000 IU cholecalciferol or placebo every 2 weeks. Cerebral and spinal cord MRI, blood sampling for vitamin D, calcium, creatinine, and urinalysis were performed at the initial visit (D0) and after 3 months (M3). Patients underwent follow-up visits at 3, 6, 12, 18 and 24 months and follow-up MRIs at 3, 12 and 24 months to assess safety and disease activity. Five MS patients treated with vitamin D (mean age 42 years-old) and 5-matched MS patients treated with placebo (39 years-old) were selected (Table n°1). Plasmatic vitamin D levels were assayed using the electrochemiluminescence on Cobas 8000 (Roche™) on the e801 module.

### Blood sampling and PBMC preparation

Blood samples of participants of the D-lay MS study was collected in EDTA-containing tubes at D0 and M3 of treatment (placebo or vitamin D). PBMCs were isolated on Ficoll® gradient and cryopreserved in liquid nitrogen at a concentration of 5 × 10^6^ cells per ml in PBS solution containing using 4% human albumin and 10% DMSO.

PBMCs were rapidly thawed in a water bath at 37°C, immediately resuspended in cell culture medium (RPMI 1640, 10% FCS, L-glutamine, penicillin (50U/mL), streptomycine (50 µg/mL), sodium pyruvate 1 mM, Hepes 10 mM, and non-essential amino acids (both Life Technologies), centrifuged (350 X g for 7min at 20°C) and washed once with cell staining Buffer (BioLegend). The PBMC pellet was resuspended and cells were stained with Human TruStain FcX Fc Blocking Reagent (BioLegend) for 10 min at 4°C and then with fixable viability dye DAPI (Merck), Vybrant Ruby (Life Technologies) and 35 TotalSeq-B antibodies (Supplemental Table 1) for 30 min on ice. Live cells were sort purified as a singlet on a BD FACSMelody™.

Sorted cells were washed in 0,4% BSA in PBS. Approximately 5,000 cells per samples were loaded on a 10x chip and run onto the 10x Chromium controller using Chromium Next GEM Single Cell 3’ Reagents Kits v3.1 with Feature Barcoding technology for Cell-Surface Protein (10x Genomics), according to the manufacturer’s protocol.

Gene expression and cell-surface protein expression libraries were multiplexed using individual Chromium i7 sample indices. Gene expressions libraries were sequenced on 3 lanes of a NovaSeq6000 S4 flow cell using 28-92nt paired-ends reads and 8nt for the i7 index. Cell-surface protein expression libraries were sequenced on a NovaSeq6000 SP flow cell using 28-92nt paired-ends reads and 8nt for the i7 index. Sequencing quality was assessed using SAV (Sequence Analysis Viewer) software from Illumina, raw data quality was assessed using FastQC (v0.11.9, Babraham Institute) and contaminant screening was performed using Fastq Screen (v0.15.1, Babraham Institute).

### Single-cell sequencing data processing

Cell Ranger software (10x Genomics, v.6.1.1) was used to demultiplex samples, process raw data, align reads to the GRCh38 human reference genome and summarize unique molecular identifier (UMI) counts. Filtered gene-barcode and cell-surface protein expression-barcode matrices containing only barcodes with UMI counts that passed the threshold for cell detection were aggregated into a single matrix and used for further analysis. Then, the aggregated filtered UMI count matrix was processed using the R package Seurat (version 4.3.0). Cells that expressed fewer than 200 genes and/or >15% mitochondrial reads, and genes expressed in fewer than three cells were removed from the count matrix. After quality control, only raw gene counts in high-quality cells were subjected to log-normalization, identification of high-variable genes by using the vst method, scaling, and regression against the number of UMIs and mitochondrial RNA content per cell. An unbiased calculation of the k-nearest neighbors, was applied to generate the neighborhood graph and embedding using UMAP. Annotation of Seurat clusters was manually curated using a combination of surface markers for each cluster and visual inspection of key markers using UMAP visualization. After initial cluster annotation, all clusters were split into CD4^+^, CD8^+^ and B-cells subsets and reanalyzed. After sub-setting, integration using reciprocal principal component analysis was performed to remove batch effects and the integrated assay was used for principal component analysis and unsupervised clustering. Seurat subclusters were annotated using a combination of canonical protein and mRNA markers.

Clustering of human PBMCs was performed using the following parameters: 18 PCs and resolution 1. Sub-clustering of human CD4^+^, CD8^+^ T-cells and B-cells was performed using the following parameters: 12 PCs and resolution 1.

To assess the quantitative effect of vitamin D, lymphocyte subsets were counted per patient and transformed into percentages relative to each patient. Two-way ANOVA with a false discovery correction according to the Benjamini-Hochberg approach was performed to assess the interaction of time and treatment. A Mann-Whitney test was performed on the M3/D0 percentage ratios.

### Differential gene expression between vitamin D and placebo groups

Differentially expressed genes were identified by two methods. The first one calculated the mean expression of each gene per patient and per subset. Means were aggregated by treatment group and time (D0 and M3). A ratio of M3/D0 means was then calculated followed by a Welch Test. A gene was considered significant with a p value < 0.05. The fold-change was then obtained by vitamin D/placebo ratio and considered as meaningful when > 1.2. The second method was to transform the scRNA-seq data to a pseudo-bulk RNA-seq data and perform a differential expression analysis using DESeq2 R package (v.1.38.3).

### Pathway Analysis

Pathway enrichment analysis was performed using Single Cell Pathway Analysis (SCPA) Package (v.1.5.3) (37) from CITE-seq data in RStudio® (Integrated Development for R. RStudio, PBC, Boston, MA) using Reactome Pathway Database (https://reactome.org) and M3-D0 for each treatment group. A pathway was considered significantly enriched with an adjusted p-value < 0.05.

### Validation cohort, Flow Cytometry and Cell Lysis

Eight MS patients treated with vitamin D (mean age 37.4 years-old) and 8-matched placebo (40.6 years-old) were selected (Table n°1). PBMCs were thawed, resuspended in (RPMI 1640, 10% FCS, L-glutamine, penicillin (50U/mL), streptomycine (50 µg/mL), sodium pyruvate 1 mM, Hepes 10 mM, and non-essential amino acids (both Life Technologies) and centrifuged (350 X g for 7min at 20°C). After resuspension in calcium- and magnesium-free PBS, PBMCs were stained with Zombie NIR–APC-Cy7 (viability dye, 1:3000) for 15min at 4°C. After incubation with FcBlock (PBS + 12% human serum AB (Merck)) for 10 min at 4°C, cells were divided into two panels and labelled as follows: *panel 1*: CD4 (BUV496, clone: SK3, Becton Dickinson), CD27 (BUV737, clone: M-T2H, Becton Dickinson), CD8a (Pacific Blue, clone: RPA-T8, BioLegend), CD45RA (BV510, clone: HI100, Becton Dickinson), CCR7 (BV787, clone: G043H7, BioLegend), IgD (FITC, clone: IA6-2, BioLegend), CD19 (PerCP-Cy5.5, clone: HIB19, BioLegend), CD3 (AF700, clone: OKT3, BioLegend) for memory B-cells, naive B-cells, naive CD4 and CD8 T-cells; and *panel 2*: CD4 (BUV496, clone: SK3, Becton Dickinson), CD8a (Pacific Blue, clone: RPA-T8, BioLegend), CD3 (AF700, clone: OKT3, BioLegend), CD127 (BV650, clone: 7019D5, BioLegend), CCR6 (BV711, clone: MI-15, Becton Dickinson), CD161 (FITC, clone: DX12, Becton Dickinson), CD25 (PE-CF594, clone: BC96, Becton Dickinson), CXCR3 (APC, clone: G025H7, BioLegend) for Th1, Th17, Treg and MAIT cells. For each cell-type and each panel, 100 cells were sorted in 5 µL RT Mix (Superscript® VILO™ cDNA Synthesis Kit, 20 U/ µL SUPERase-In™ RNase Inhibitor, 10% NP-40 Detergent Surfact-Amps Solution, ThermoFisher Scientific) using a BD FACSAria™ Fusion and stored at −80°C.

### High-throughput qPCR

#### PCR Primers

A set of 190 PCR primers (Supplemental Table 2) corresponding to the 178 genes of interest plus 12 housekeeping genes (*ACTB, GAPDH, HPRT1, IPO8, POLR2F, PPIA, RPL10, RPLP0, RPS18, TFRC, UBC, UBE2D2*) was prepared by Fluidigm for the Fluidigm DeltaGene Assay (Fluidigm Co., San Francisco, CA, USA, https://d3.fluidigm.com/account/login). Fluidigm analyses were carried out on several sub-populations at a time. Only the three most robust reference genes per population were used for analysis.

#### Reverse transcription

Reverse transcription was performed using the SuperScript® VILO™ cDNA Synthesis Kit (ThermoFisher Scientific, Waltham, MA, United States) according to the manufacturer’s protocol. The reverse transcription conditions were as follows: 25°C for 5min, 50°C for 30min, 55°C for 25min, 60°C for 5min and 70°C for 10min. Complementary DNAs (cDNAs) were stored at −20°C.

#### Pre-amplification

A pre-amplification step was included to increase the number of cDNA copies to a detectable level and to allow the concurrent amplification of the different gene expression targets, using the PreAmp Master Mix (Fluidigm Co., San Francisco, CA, USA). The pre-amplified products were treated with exonuclease I to remove the unincorporated primers and diluted 5-fold with the DNA suspension buffer before being assayed by PCR. The pre-amplified DNAs were stored at −20°C until testing.

#### Analysis of gene expression

Gene expression was analyzed by the Fluidigm qPCR procedure, using the Biomark HD™ microfluidic device. Primer-probe sets and samples were transferred to an integrated fluidic circuits (IFC) plate and loaded into an automated controller that prepares the nanoliter reactions. The IFC plate was run on the Biomark instrument, which uses a thermal cycler for real-time quantitative PCR following this cycle conditions: 70°C for 40 min, 60°C for 30 s, 95°C for 60 s, followed by 30 cycles of 96°C for 5 s and 60°C for 20 s, using the 96.96 dynamic array™ IFC (microfluidic chip). Distilled water, instead of cDNA, was used as negative control. Melting curves were generated for each gene by increasing the temperature from 60°C to 95°C while the fluorescence was measured. Non-detectable expression was considered a Ct of 24. ^Δ^Ct values were calculated relative to the geometric mean of three housekeeping genes in the 96-gene set for each subset. ^ΔΔ^Ct were calculated using D0 for reference for each gene in each subpopulation. Fold changes were calculated as 2^−ΔΔCt^.

### Statistical analysis

The Mann-Whitney test was used to evaluate statistical differences in the gene expression means between placebo and vitamin D group. P-values lower than 0.05 were considered statistically significant. P-values lower than 0.20 were considered as part of the discovery step of genes potentially modulated by vitamin D. Statistical analyses and box plot representation were performed using RStudio®. Heat-map was performed using the free-software Perseus® (Max-Planck-Institute of Biochemistry, Martinsried, Germany, version 2.0.7.0). A gradient coloring scheme was applied to display upregulated (above 2-fold expression (in log2 display above 1, red color)) *vs.* downregulated (below 2-fold expression (in log2 display below −1, blue color)) genes. The heatmap was hierarchically clustered by Pearson’s correlation distance using row average linkage.

### Study Approval

The study protocol was approved by a national review board, the CPP Sud-Méditerranée III ethics committee (2013.02.08 bis on March, 6th 2013) and health authorities (EudraCT 2013-000910-40, ANSM 130342A-31). Written informed consent was obtained from all the study participants before any study-related procedures were performed (ClinicalTrials.gov identifier NCT01817166). The study was conducted in accordance with the International Conference on Harmonization Guidelines for Good Clinical Practice and the Declaration of Helsinki.

## Data and code Availability

Anonymized raw sequencing data and counts matrix have been deposited in the Gene Expression Omnibus (GEO) with accession numbers GEO: GSE239626. The code used for the analysis can be shared on request.

## Author Contributions

M.G conducting experiments, acquiring data, analyzing data, writing and reviewing the manuscript; A.L acquiring, analyzing data and reviewing the manuscript; M.R analyzing data; S.K reviewing the manuscript; A.A reviewing the manuscript; S.S reviewing the manuscript; B.E reviewing the manuscript; P.M reviewing the manuscript; E.T designed research studies, writing and reviewing the manuscript.

## Acknowledgments

This research was supported by Agence Nationale de la Recherche N° ANR-19-CE14-0043, the Swiss National Science Foundation (SNSF grant n° 310030E_189312), and CHU Nîmes. We thank Brigitte Lafont, Hanane Agherbi for logistic assistance and the ressource biological center from Nîmes University Hospital and Nantes University Hospital for providing PBMC samples.

We thank Amélie Sarrazin, Myriam Boyer-Clavel and Stéphanie Viala from MRI platform for technical assistance in cell sorting experiments.

We thank Laurent Journot, Dany Severac, Simon George from MGX platform for CITE-seq experiments and technical assistance. MGX acknowledges financial support from France Génomique National infrastructure, funded as part of “Investissement d’avenir” program managed by Agence Nationale pour la Recherche (N° ANR-10-INBS-09).

We thank Christelle Reynes, Manuela Pastore and Lea Besnard from STATABIO platform for statistical R scripts and technical assistance.

Flow cytometry for sort experiments in the validation step were performed at INFINITY INSERM UMR1291 core facility connected to the ‘Toulouse Réseau Imagerie’ network. We thank Anne-Laure Iscache and Hugo Garnier for technical assistance.

We thank Sandra Espeout-Fois and Aurore Vernet from CIRAD for technical assistance in HT-qPCR experiments.

## Abbreviations

CIS: Clinically Isolated Syndrome
EAE: Experimental Autoimmune Encephalomyelitis
HT-qPCR: High-Throughput qPCR
MS: Multiple Sclerosis
SCPA: Single-Cell Pathway Analysis
UMAP: Uniform Manifold Approximation and Projection
UMI: Unique Molecular Identifier

## References

1. International Multiple Sclerosis Genetics Consortium, Patsopoulos NA, Baranzini SE, Santaniello A, Shoostari P, Cotsapas C, et al. Multiple sclerosis genomic map implicates peripheral immune cells and microglia in susceptibility. Science. 27 sept 2019;365(6460):eaav7188.

2. Ramagopalan SV, Dobson R, Meier UC, Giovannoni G. Multiple sclerosis: risk factors, prodromes, and potential causal pathways. The Lancet Neurology. juill 2010;9(7):727□39.

3. Thouvenot E, Orsini M, Daures JP, Camu W. Vitamin D is associated with degree of disability in patients with fully ambulatory relapsing-remitting multiple sclerosis. Eur J Neurol. mars 2015;22(3):564□9.

4. Ramagopalan SV, Heger A, Berlanga AJ, Maugeri NJ, Lincoln MR, Burrell A, et al. A ChIP-seq defined genome-wide map of vitamin D receptor binding: Associations with disease and evolution. Genome Research. 1 oct 2010;20(10):1352□60.

5. Galoppin M, Kari S, Soldati S, Pal A, Rival M, Engelhardt B, et al. Full spectrum of vitamin D immunomodulation in multiple sclerosis: mechanisms and therapeutic implications. Brain Communications. 4 juill 2022;4(4):fcac171.

6. Hupperts R, Smolders J, Vieth R, Holmøy T, Marhardt K, Schluep M, et al. Randomized trial of daily high-dose vitamin D _3_ in patients with RRMS receiving subcutaneous interferon β-1a. Neurology. 8 oct 2019;10.1212/WNL.0000000000008445.

7. Camu W, Lehert P, Pierrot-Deseilligny C, Hautecoeur P, Besserve A, Jean Deleglise AS, et al. Cholecalciferol in relapsing-remitting MS: A randomized clinical trial (CHOLINE). Neurol Neuroimmunol Neuroinflamm. sept 2019;6(5):e597.

8. Cassard SD, Fitzgerald KC, Qian P, Emrich SA, Azevedo CJ, Goodman AD, et al. High-dose vitamin D3 supplementation in relapsing-remitting multiple sclerosis: a randomised clinical trial. eClinicalMedicine. mai 2023;59:101957.

9. Shevtsov A, Raevskiy M, Stupnikov A, Medvedeva Y. In Silico Drug Repurposing in Multiple Sclerosis Using scRNA-Seq Data. IJMS. 4 janv 2023;24(2):985.

10. Kim D, Witt EE, Schubert S, Sotirchos E, Bhargava P, Mowry EM, et al. Peripheral T-Cells, B-Cells, and Monocytes from Multiple Sclerosis Patients Supplemented with High-Dose Vitamin D Show Distinct Changes in Gene Expression Profiles. Nutrients. 9 nov 2022;14(22):4737.

11. Stoeckius M, Hafemeister C, Stephenson W, Houck-Loomis B, Chattopadhyay PK, Swerdlow H, et al. Simultaneous epitope and transcriptome measurement in single cells. Nat Methods. sept 2017;14(9):865□8.

12. Huang Y, Chen L, Zhou Y, Liu H, Yang J, Liu Z, et al. UXT-V1 protects cells against TNF-induced apoptosis through modulating complex II formation. Heldin CH, éditeur. MBoC. 15 avr 2011;22(8):1389□97.

13. Chang LS, Wang JT, Doong SL, Lee CP, Chang CW, Tsai CH, et al. Epstein-Barr Virus BGLF4 Kinase Downregulates NF-κB Transactivation through Phosphorylation of Coactivator UXT. J Virol. 15 nov 2012;86(22):12176□86.

14. Duddy M, Niino M, Adatia F, Hebert S, Freedman M, Atkins H, et al. Distinct Effector Cytokine Profiles of Memory and Naive Human B Cell Subsets and Implication in Multiple Sclerosis. The Journal of Immunology. 15 mai 2007;178(10):6092□9.

15. Li R, Rezk A, Miyazaki Y, Hilgenberg E, Touil H, Shen P, et al. Proinflammatory GM-CSF–producing B cells in multiple sclerosis and B cell depletion therapy. Sci Transl Med [Internet]. 21 oct 2015 [cité 28 juill 2023];7(310). Disponible sur: https://www.science.org/doi/10.1126/scitranslmed.aab4176

16. Krensky AM, Clayberger C. Biology and clinical relevance of granulysin. Tissue Antigens. mars 2009;73(3):193□8.

17. Deng A, Chen S, Li Q, Lyu S chen, Clayberger C, Krensky AM. Granulysin, a Cytolytic Molecule, Is Also a Chemoattractant and Proinflammatory Activator. The Journal of Immunology. 1 mai 2005;174(9):5243□8.

18. Afsal K, Selvaraj P, Harishankar M. 1, 25-dihydroxyvitamin D3 downregulates cytotoxic effector response in pulmonary tuberculosis. International Immunopharmacology. sept 2018;62:251□60.

19. Palau N, Julià A, Ferrándiz C, Puig L, Fonseca E, Fernández E, et al. Genome-wide transcriptional analysis of T cell activation reveals differential gene expression associated with psoriasis. BMC Genomics. déc 2013;14(1):825.

20. Chauss D, Freiwald T, McGregor R, Yan B, Wang L, Nova-Lamperti E, et al. Autocrine vitamin D signaling switches off pro-inflammatory programs of TH1 cells. Nat Immunol [Internet]. 11 nov 2021 [cité 23 nov 2021]; Disponible sur: https://www.nature.com/articles/s41590-021-01080-3

21. Liu G, Li F, Chen M, Luo Y, Dai Y, Hou P. SNRPD1/E/F/G Serve as Potential Prognostic Biomarkers in Lung Adenocarcinoma. Front Genet. 3 mars 2022;13:813285.

22. Luo Y, Xu C, Wang B, Niu Q, Su X, Bai Y, et al. Single-cell transcriptomic analysis reveals disparate effector differentiation pathways in human Treg compartment. Nat Commun. 23 juin 2021;12(1):3913.

23. Caldwell RB, Braselmann H, Schoetz U, Heuer S, Scherthan H, Zitzelsberger H. Positive Cofactor 4 (PC4) is critical for DNA repair pathway re-routing in DT40 cells. Sci Rep. 4 juill 2016;6(1):28890.

24. Hanel A, Carlberg C. Time-Resolved Gene Expression Analysis Monitors the Regulation of Inflammatory Mediators and Attenuation of Adaptive Immune Response by Vitamin D. IJMS. 14 janv 2022;23(2):911.

25. Häusler D, Torke S, Peelen E, Bertsch T, Djukic M, Nau R, et al. High dose vitamin D exacerbates central nervous system autoimmunity by raising T-cell excitatory calcium. Brain. 1 sept 2019;142(9):2737□55.

26. Krumbholz M, Theil D, Cepok S, Hemmer B, Kivisäkk P, Ransohoff RM, et al. Chemokines in multiple sclerosis: CXCL12 and CXCL13 up-regulation is differentially linked to CNS immune cell recruitment. Brain. 1 janv 2006;129(1):200□11.

27. Killick J, Hay J, Morandi E, Vermeren S, Kari S, Angles T, et al. Vitamin D/CD46 Crosstalk in Human T Cells in Multiple Sclerosis. Front Immunol. 24 nov 2020;11:598727.

28. Bendix M, Greisen S, Dige A, Hvas CL, Bak N, Jørgensen SP, et al. Vitamin D increases programmed death receptor-1 expression in Crohn’s disease. Oncotarget. 11 avr 2017;8(15):24177□86.

29. Dardalhon V, Schubart AS, Reddy J, Meyers JH, Monney L, Sabatos CA, et al. CD226 Is Specifically Expressed on the Surface of Th1 Cells and Regulates Their Expansion and Effector Functions. The Journal of Immunology. 1 août 2005;175(3):1558□65.

30. Zhang R, Zeng H, Zhang Y, Chen K, Zhang C, Song C, et al. CD226 ligation protects against EAE by promoting IL-10 expression *via* regulation of CD4+ T cell differentiation. Oncotarget. 12 avr 2016;7(15):19251□64.

31. Leandro MJ. B-cell subpopulations in humans and their differential susceptibility to depletion with anti-CD20 monoclonal antibodies. Arthritis Res Ther. 2013;15(Suppl 1):S3.

32. Gaglia MM. Anti-viral and pro-inflammatory functions of Toll-like receptors during gamma-herpesvirus infections. Virol J. déc 2021;18(1):218.

33. Ersing I, Bernhardt K, Gewurz B. NF-κB and IRF7 Pathway Activation by Epstein-Barr Virus Latent Membrane Protein 1. Viruses. 21 juin 2013;5(6):1587□606.

34. Hoepner R, Bagnoud M, Pistor M, Salmen A, Briner M, Synn H, et al. Vitamin D increases glucocorticoid efficacy via inhibition of mTORC1 in experimental models of multiple sclerosis. Acta Neuropathol. sept 2019;138(3):443□56.

35. Polman CH, Reingold SC, Edan G, Filippi M, Hartung HP, Kappos L, et al. Diagnostic criteria for multiple sclerosis: 2005 revisions to the “McDonald Criteria”. Ann Neurol. déc 2005;58(6):840□6.

36. Swanton JK, Rovira A, Tintore M, Altmann DR, Barkhof F, Filippi M, et al. MRI criteria for multiple sclerosis in patients presenting with clinically isolated syndromes: a multicentre retrospective study. The Lancet Neurology. août 2007;6(8):677□86.

37. Bibby JA, Agarwal D, Freiwald T, Kunz N, Merle NS, West EE, et al. Systematic single-cell pathway analysis to characterize early T cell activation. Cell Reports. nov 2022;41(8):111697.

