## Supplementary Material for "CITE-seq reveals inhibition of NF-κB pathway in B cells from vitamin D-treated multiple sclerosis patients"

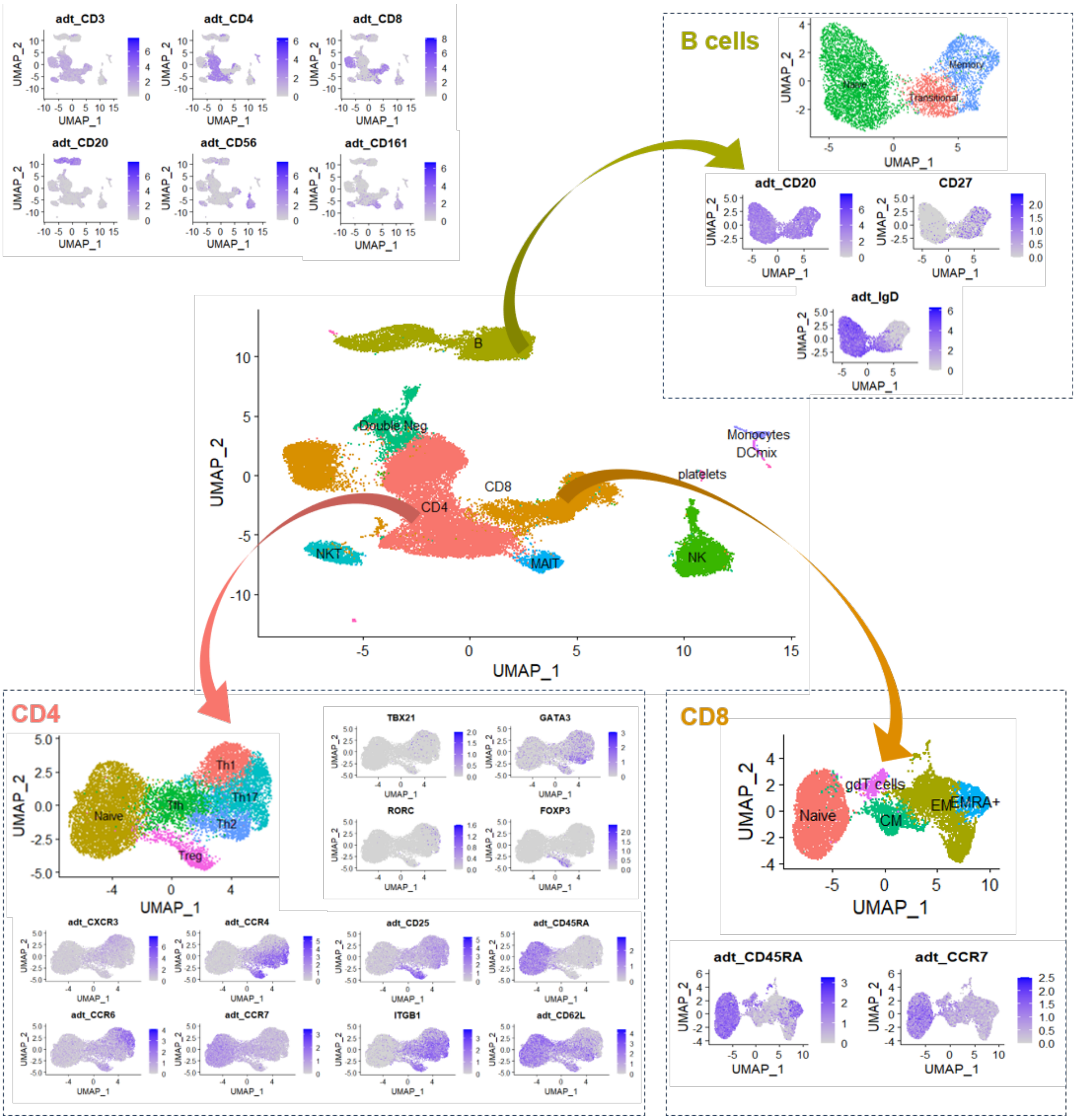


**Figure Sup 1**. **Annotation of Seurat clusters using a combination of surface markers and gene expression and visual inspection of key markers using UMAP visualization.**
Expression of characteristic surface markers and gene expression was used to annotate major PBMC populations including CD4$^+$ T-cells (CD4$^+$), CD8$^+$ T-cells (CD8$^+$), B-cells (CD19$^+$), natural killer (NK) cells (CD56$^+$), natural killer T-cells (NKT; CD3$^+$CD56$^+$), MAIT cells (CD161$^+$), monocytes, platelets, and dendritic cells (DCs). CD4$^+$ subsets were further annotated with a combination of surface markers and gene expression including CD45RA$^+$ CCR7$^+$ (naive), CXCR3$^+$CCR6$^-$TBX21$^+$ (Th1), CXCR3$^-$CCR6$^-$CCR4$^+$GATA3$^+$ (Th2), CXCR3$^-$CCR6$^+$RORC$^+$ (Th17), CD45RA$^-$CXCR5$^+$ICOS (Tfh), CD45RA$^-$CD25$^+$FOXP3$^+$ (Treg). CD8$^+$ T cells subsets were annotated with CD45RA$^+$ CCR7$^+$ (naive), CD45RA$^-$ CCR7$^+$ (central memory, CM), CD45RA$^-$CCR7$^-$ (effector memory, EM), CD45RA$^+$CCR7$^-$ (effector, EMRA$^+$), and γδTCR$^+$ (*γδ* T cells). B-cells subsets were annotated using CD27$^-$IgD$^+$ (naïve), CD27$^+$IgD$^+$ (transitional) and CD27$^+$IgD$^-$ (memory).

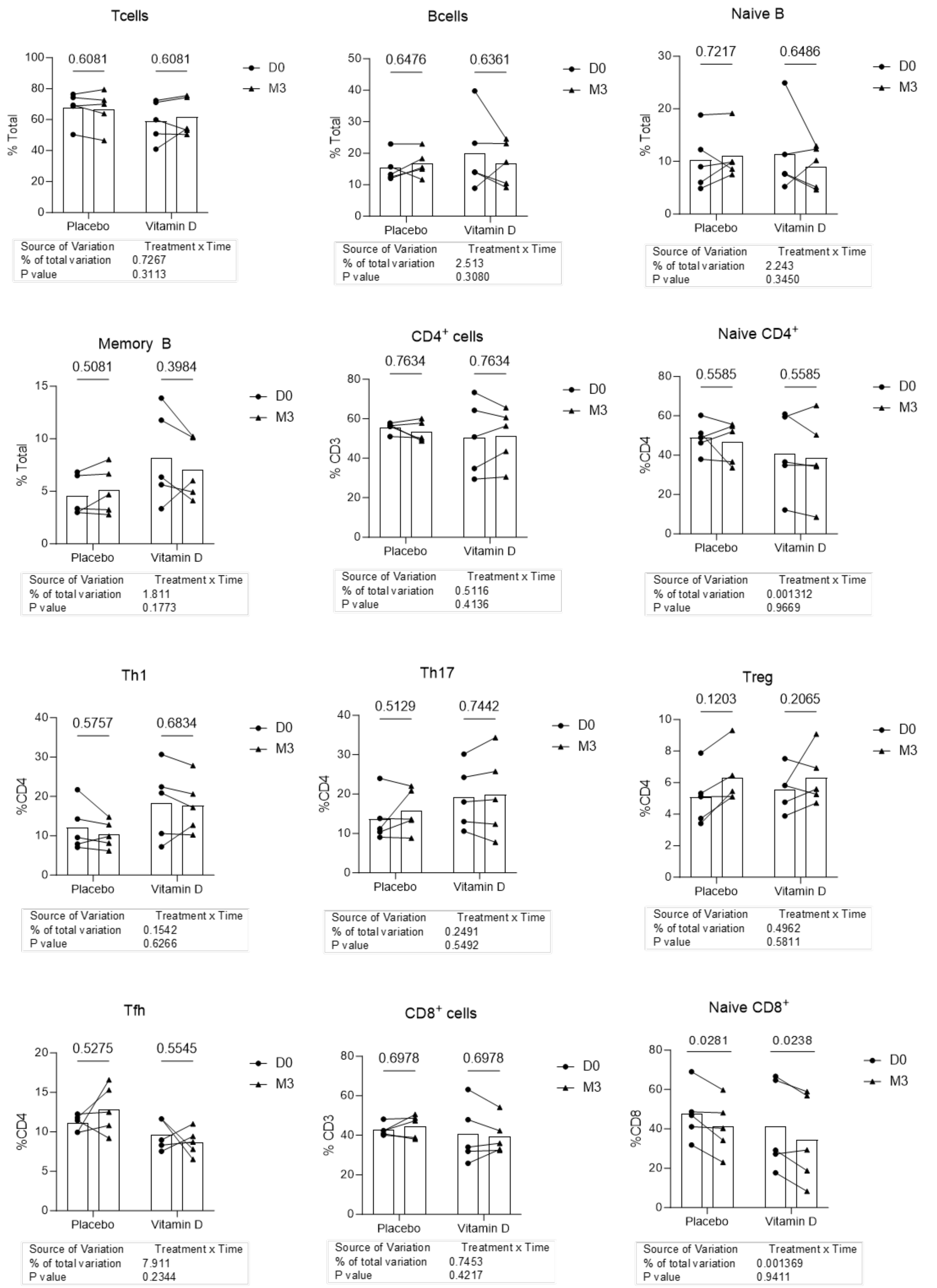

Tcells
0.6081
0.6081
D0
M3
% Total
Placebo
Vitamin D
Source of Variation Treatment x Time
% of total variation 0.7267
P value 0.3113
Bcells
0.6476
0.6361
% Total
Source of Variation Treatment x Time
% of total variation 2.513
P value 0.3080
Naive B
0.7217
0.6486
% Total
Source of Variation Treatment x Time
% of total variation 2.243
P value 0.3450
Memory B
0.5081
0.3984
% Total
Source of Variation Treatment x Time
% of total variation 1.811
P value 0.1773
CD4+ cells
0.7634
0.7634
% CD3
Source of Variation Treatment x Time
% of total variation 0.5116
P value 0.4136
Naive CD4+
0.5585
0.5585
%CD4
Source of Variation Treatment x Time
% of total variation 0.001312
P value 0.9669
Th1
0.5757
0.6834
%CD4
Source of Variation Treatment x Time
% of total variation 0.1542
P value 0.6266
Th17
0.5129
0.7442
%CD4
Source of Variation Treatment x Time
% of total variation 0.2491
P value 0.5492
Treg
0.1203
0.2065
%CD4
Source of Variation Treatment x Time
% of total variation 0.4962
P value 0.5811
Tfh
0.5275
0.5545
%CD4
Source of Variation Treatment x Time
% of total variation 7.911
P value 0.2344
CD8+ cells
0.6978
0.6978
% CD3
Source of Variation Treatment x Time
% of total variation 0.7453
P value 0.4217
Naive CD8+
0.0281
0.0238
%CD8
Source of Variation Treatment x Time
% of total variation 0.001369
P value 0.9411

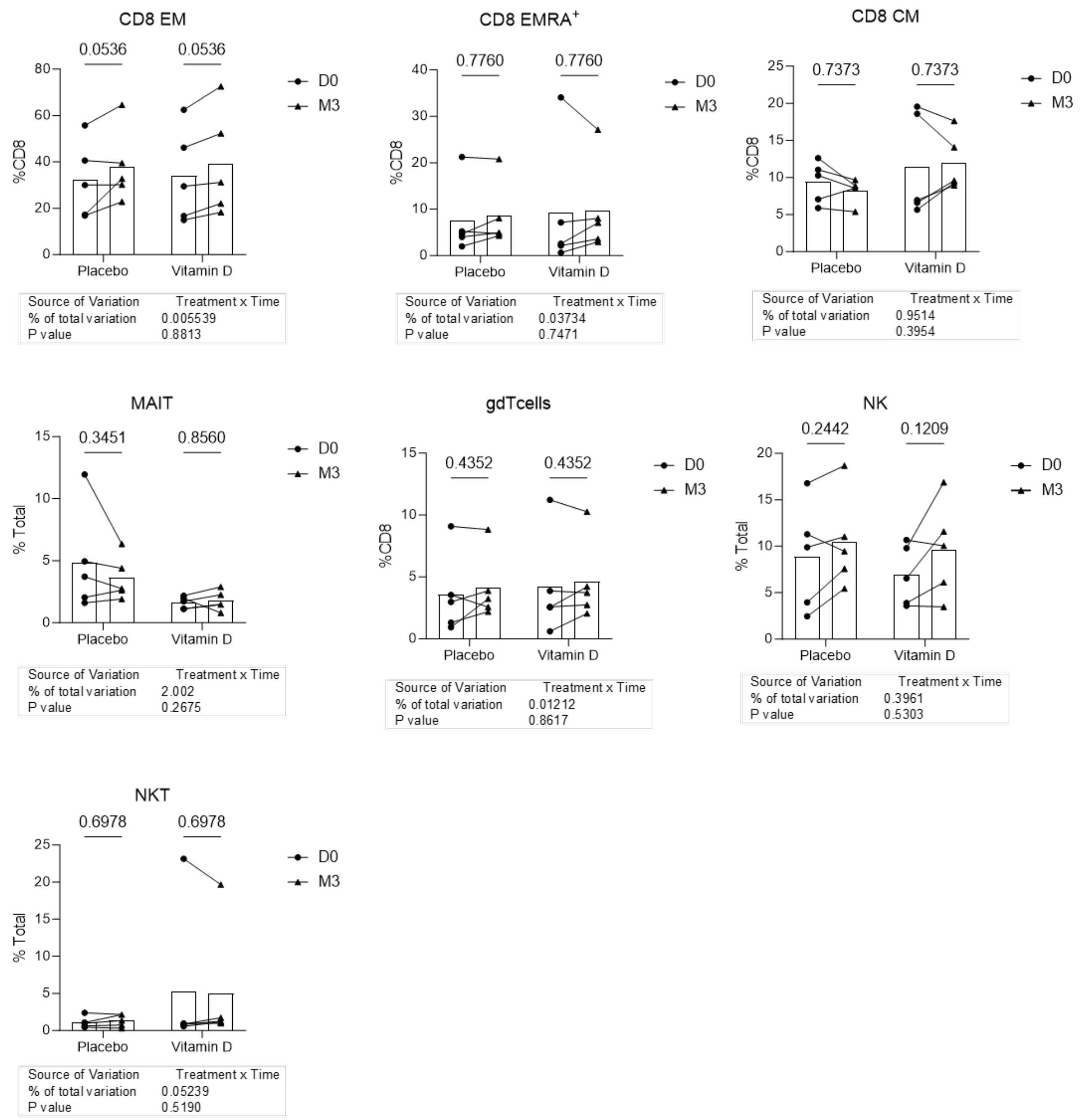


**Figure Sup 2. Changes in the proportions of lymphocytes subsets after treatment with placebo or vitamin D**.

Significance was determined by 2way-ANOVA with original FDR BH correction.

Naive B-cells

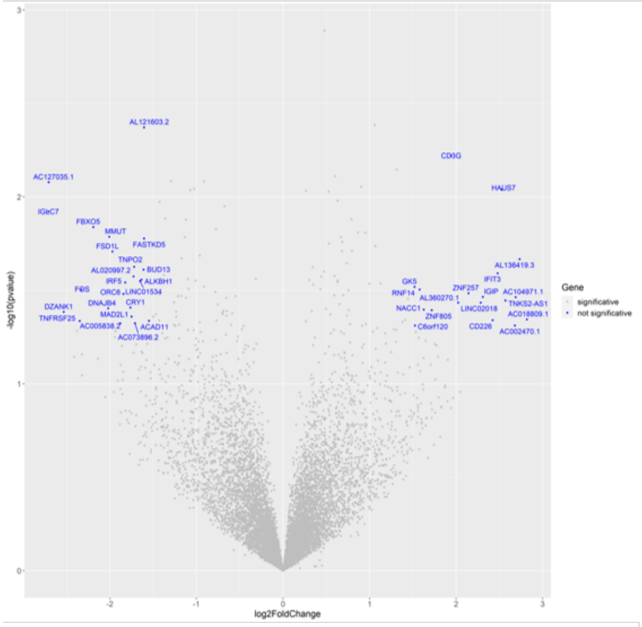


Memory B-cells

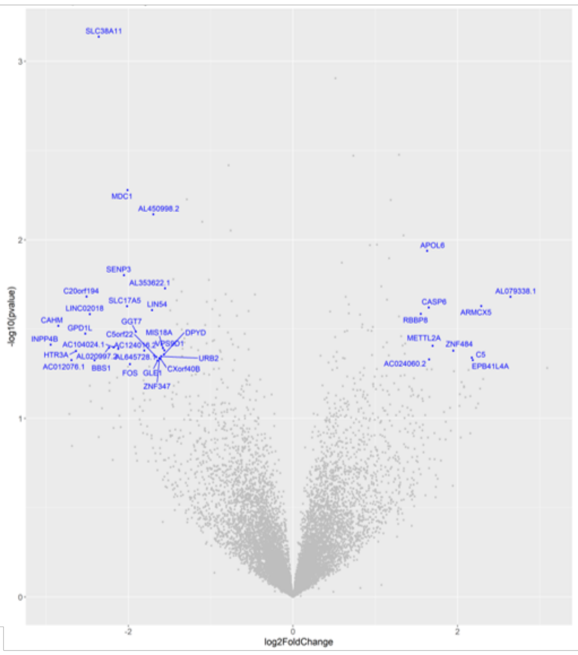


MAIT cells

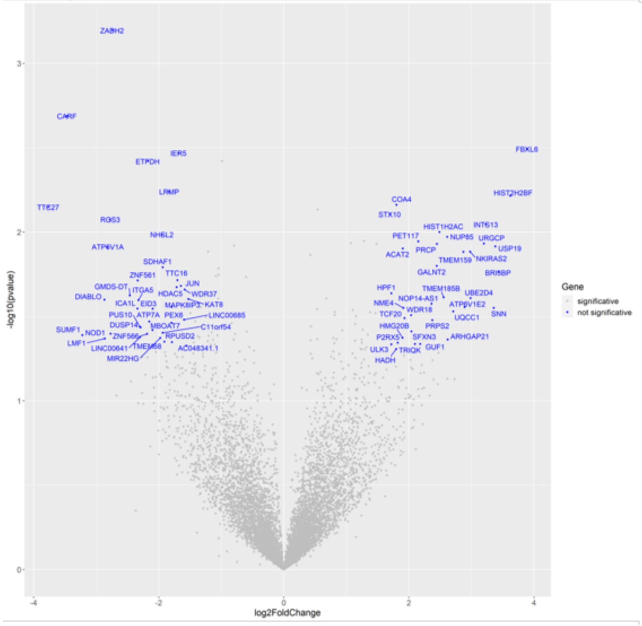


Naive CD8 T-cells

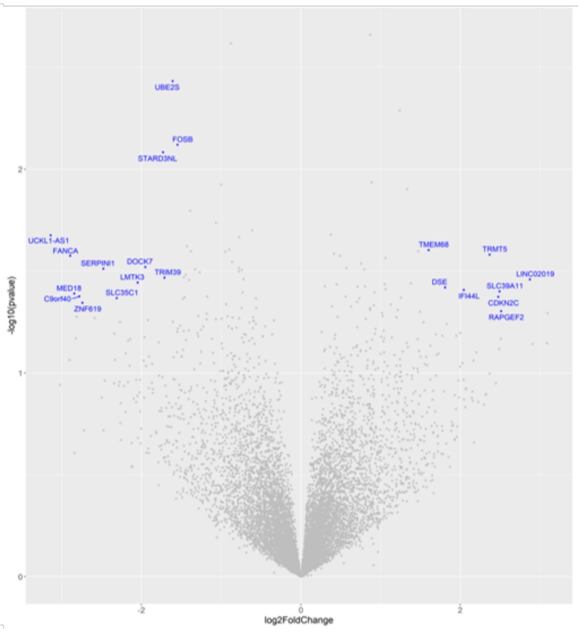

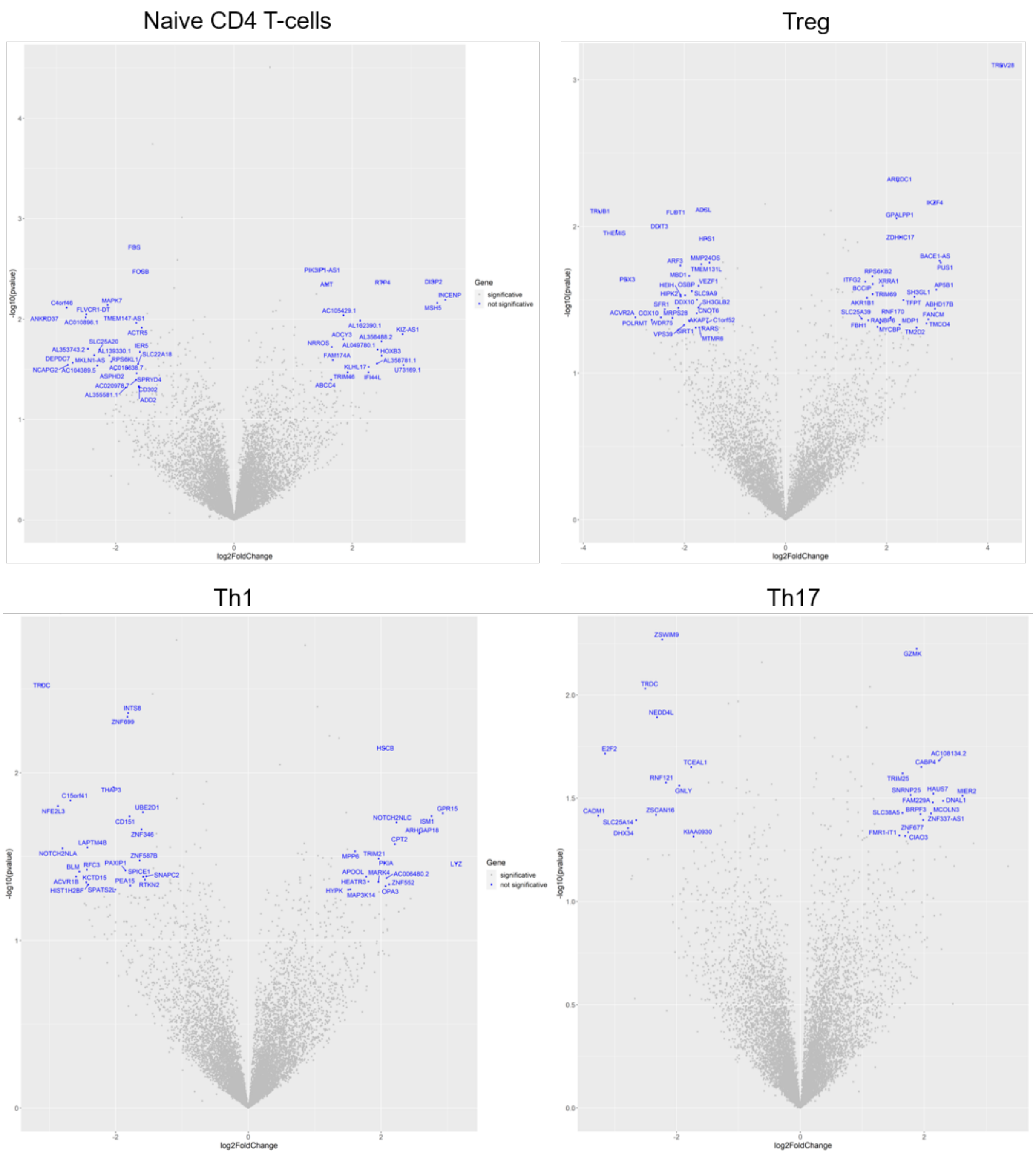


**Figure Sup 3. Volcano plot representation of regulated genes identified by DESeq2 analysis.**

| Antibody | Clone | CITE-seq Barcode |
|---|---|---|
| CCR4 | L291H4 | AGCTTACCTGCACGA |
| CCR5 | J418F1 | CCAAAGTAAGAGCCA |
| CCR6 | G034E3 | GATCCCTTTGTCACT |
| CCR7 | G043H7 | AGTTCAGTCAACCGA |
| CD11a | TS2/4 | TATATCCTTGTGAGC |
| CD127 | A019D5 | GTGTGTTGTCCTATG |
| CD138 | MI15 | ACTCTTTCGTTTACG |
| CD161 | HP-3G10 | GTACGCAGTCCTTCT |
| CD20 | 2H7 | TTCTGGGTCCCTAGA |
| CD226 | 11A8 | TCTCAGTGTTTGTGG |
| CD24 | ML5 | AGATTCCTTCGTGTT |
| CD25 | BC96 | TTTGTCCTGTACGCC |
| CD27 | O323 | GCACTCCTGCATGTA |
| CD3 | SK7 | TATCCCTTGGGATGG |
| CD38 | HIT2 | TGTACCCGCTTGTGA |
| CD4 | RPA-T4 | TGTTCCCGCTCAACT |
| CD45RA | HI100 | TCAATCCTTCCGCTT |
| CD49b | P1E6-C5 | GCTTTCTTCAGTATG |
| CD49d | 9F10 | CCATTCAACTTCCGG |
| CD56 | 5.1H11 | TCCTTTCCTGATAGG |
| CD62L | DREG-56 | GTCCCTGCAACTTGA |
| CD71 | CY1G4 | CCGTGTTCCTCATTA |
| CD8 | SK1 | GCGCAACTTGATGAT |
| CTLA-4 | BNI3 | ATGGTTCACGTAATC |
| CXCR3 | G025H7 | GCGATGGTAGATTAT |
| CXCR4 | 12G5 | TCAGGTCCTTTCAAC |
| CXCR5 | J252D4 | AATTCAACCGTCGCC |
| GITR | 108-17 | ACCTTTCGACACTCG |
| HLA-DR | L243 | AATAGCGAGCAAGTA |
| ICOS | C398.4A | CGCGCACCCATTAAA |
| IgD | IA6-2 | CAGTCTCCGTAGAGT |
| IgM | MHM-88 | TAGCGAGCCCGTATA |
| LAG3 | 11C3C65 | CATTTGTCTGCCGGT |
| TCRVa7.2 | 3C10 | TACGAGCAGTATTCA |
| TCRγ/δ | B1 | CTTCCGATTCATTCA |

**Table Sup 1. Total Seq-B antibodies used for CITE-seq (human, BioLegend).**

| Gene symbol | Accession number | Forward Primer | Reverse Primer |
|---|---|---|---|
| ACTB | NM_001101.3 | CCAACCGCGAGAAGATGAC | TAGCACAGCCTGGATAGCAA |
| RPLP0 | NM_053275.3 | GCGACCTGGAAGTCCAAC | CACATTGTCTGCTCCCACAA |
| UBE2D2 | NM_181838.1 | ACAGTGGTCTCCAGCACTAA | GGATCATCTGGATTGGGATCAC |
| HPRT1 | NM_000194.2 | GCTTTCCTTGGTCAGGCAGTA | ACTTCGTGGGGTCCTTTTCAC |
| UBC | NM_021009. | GGGGTGTCTAAGTTTCCCCTT | GGGAATGCAACAACTTTATTGAAAGG |
| RPS18 | NM_022551.2 | CGGAAAATAGCCTTTGCCATCA | TCCTCAACACCACATGAGCATA |
| PPIA | NM_001300981.1 | TCTGGTTCCTTCTGCGTGAA | CCAGGGAATACGTAACCAGACA |
| GAPDH | NM_002046. | GAGAACGGGAAGCTTGTCATCA | TGGACTCCACGACGTACTCA |
| POLR2F | NM_001301129. | AGCGAATCACCACACCATACA | ACCATCACAGGGGCACAC |
| RPL10 | NM_001303625. | TGCTCATGAGGCAGCAAAC | GAGGTCCAAGTCAAGCTCCA |
| IPO8 | NM_001190995.1 | TTCAGTGCAAAGGAAGGGGAA | ACCCCTCGAGTTAATCTCTCCA |
| TFRC | NM_001128148.1 | CCAGTTGGAGTGCTGGAGAC | AGCCTTTAAATGCAGGGACGAA |
| ALCAM | NM_002228.3 | AAGAACTCGGACCTCCTCAC | TGGATTATCAGGCGCTCCA |
| ARHGAP18 | NM_033515.N | TACCAGGAATGCGAATACCC | AAGAGGCCTTCTGTTTCCAA |
| ARRDC1 | NM_001317968.N | CCTGAACAGCATCCCAGACA | CCCGTCTTCACCAGCTTGTA |
| BCL2 | NM_000633.2 | ATGTGTGTGGAGAGCGTCAA | GTGCCGGTTCAGGTACTCA |
| C1QBP | NM_001212.N | GGGCAGAAGGTTGAAGAACA | GGCCTTCTTGCCATCATCA |
| CADM1 | NM_001301043.N | CAGACCCAGCGGTATCTAGAA | TTCCCGGGTTAAGCCTTGTA |
| CAST | NM_001190442.1 | TGTCCCTCCACTACAGAAACC | TAGGTGCTTTGGAGCTGGAA |
| CCL2 | NM_002982.3 | TAGCAGCCACCTTCATTCCC | CCTCTGCACTGAGATCTTCCTA |
| CCR4 | NM_005508. | TGCCTCACAGACCTTCCTCA | AGGCTCCTCAAGGCAGGT |
| CCR5 | NM_001100168.1 | TGAGACATCCGTTCCCCTACA | TGGCAGGGCTCCGATGTATA |
| CCR6 | NM_004367.5 | AGGCAGCGATGTCTGTGAA | AGCTCAAGCCCCAACATCA |
| CCR7 | NM_001301716.1 | TGAGGTCACGGACGATTACA | CGTCCTTCTTGGAGCACAAA |
| CD151 | NM_139029.N | ATCGCTGGTATCCTCGCCTA | GCTTGGTCATGGTGTCCTTCA |
| CD19 | NM_001178098.1 | GCAACCTGACCATGTCATTCC | TCCAGCCACCAGTCCTCA |
| CD226 | NM_006566.2 | CGTGATGAGATTGACTGTAGCC | CTCCAGCCACAAAGAGGGTA |
| CD27 | NM_001242.4 | CACTACTGGGCTCAGGGAAA | TGCTGGTCACAGTCCTTCA |
| CD28 | NM_001243077.1 | GTGGAGTCCTGGCTTGCTATA | GAGCCTGCTCCTCTTACTCC |
| CD302 | NM_014880.N | TCTACCTAATGGCAGCTCTTCC | GCCTCTTGGCCTGTTGAAAA |
| CD3E | NM_000733.3 | GCAAACCAGAAGATGCGAAC | CATCACATCCATCTCCATGCA |
| CD4 | NM_000616.4 | AAAGTTGCATCAGGAAGTGAACC | CCCACACCTCACAGGTCAAA |
| CD46 | NM_172352.2 | CCATGGTGTTGCTGCTGTAC | ACCAATGAGCTCCATAGCTTCA |
| CD53 | NM_001040033. | GGCTGCATGGGCTCTATCA | GTCACCTCAGCAAGGAGGATAA |
| CD6 | NM_006725.3 | CTGGCGGTTCAACAACTCC | GGAGTGGACAGATTGTGCAAA |
| CD69 | NM_001781.2 | TCACCCATGGAAGTGGTCAA | ACACACTTGTCAGACCCTGTA |
| CD8A | NM_001768.6 | CCATCATGTACTTCAGCCACTTCG | GCTGCGACGCGATGGT |
| CD99 | NM_001321367. | TGGCGCTGCTGCTCTT | AGGAAGGGCATCGGATAAATCG |
| CDC42SE2 | NM_020240.N | TCAACTGCTGTATTGCAGAAC | TGTGGGCTCTCCAATCATAC |
| CDK13 | NM_003718.4 | AGGCCACCTCCAGAACCTA | TGGCGTTAGCTCCAAGATCC |
| CHD2 | NM_001271.N | GACAGTTCGCTACACAGCAA | ACTGACTGCCTGAGTCTGAA |
| CLEC2D | NM_013269.4 | CAAGCTGCATGCCCAGAAA | TGATGTCCAGTTCTTGGTGTCA |
| CLK1 | NR_027856.N | AGGATTCTTGGACCTCTACCAA | CAGTCTAATCGATCGTGGTGAA |
| CTLA4 | NM_005214.4 | CATGGACACGGGACTCTACA | AATCTGGGTTCCGTTGCCTA |
| CXCL1 | NM_001511.2 | CTTGCCTCAATCCTGCATCC | AGCCACCAGTGAGCTTCC |

| | | | |
|---|---|---|---|
| CXCL12 | NM_199168.3 | GCTGGTCCTCGTGCTGAC | GAATCGGCATGGGCATCTGTA |
| CXCL13 | NM_006419.2 | GGACCCTCAAGCTGAATGGATA | ACACTGGAACTGGTAGAGTTGAA |
| CXCL2 | NM_002089.3 | CCAAAGTGTGAAGGTGAAGTCC | CGATGCGGGGTTGAGACA |
| CXCR3 | NM_001504.1 | AACTGTGGCCGAGAAAGCA | TTGAGGCAGCAGTGCATGTA |
| CYP24A1 | NM_001128915.1 | CGGGCAGAAGATTTGAGGAA | GTAAATGGTACACTCGGCGTAA |
| CYP27B1 | NM_000785. | CTCGTGTCCCAGACAAAGACA | TCCCTTGAAGTGGCATAGTGAC |
| CYTIP | NM_004288.4 | CCTGATCAGATCGTCCGGAAA | GCTTCAAGCTCCGTTCTTTTCA |
| DDIT3 | NM_001195056.1 | GAGCCAAAATCAGAGCTGGAAC | TGCTTTCAGGTGTGGTGATGTA |
| DGKD | NM_152879.N | CGTAGCTGAATCCAGTACCA | GCACACAAGATGAGCTTCC |
| DIAPH1 | NM_001079812.2 | TCCCTTCGTGTGTCTCTCAA | TAAGGAGGCCAAGCCTTCA |
| DUSP1 | NM_004417.3 | AGACATCAGCTCCTGGTTCA | CAGTGGACAAACACCCTTCC |
| DUSP2 | NM_004418.N | ACTCCAGGGCTCCTGTCTA | TGCAGGTCTGACGAGTGAC |
| E2F2 | NM_004091.N | TTCAAGCACCTGACTGAGGAC | AGTTGCCAACAGCACGGATA |
| EEF1B2 | NM_001037663.N | AGTGCGTCAGAAGCATTCAA | TCCTCCAGCATATCTGTTCCA |
| FGL2 | NM_006682.2 | AGTGTCCCAGCCAAGAACAA | CCTATTGCGTAGTAGTCAGAGCAA |
| FKBP1A | NM_001199786.N | AAGCGCGGCCAGACCT | TGGCACCATAGGCATAATCTGGAG |
| FOS | NM_005252. | CTACCACTCACCCGCAGAC | GCAGTGACCGTGGGAATGAA |
| FOSB | NM_006732.N | CCCTCTGCCGAGTCTCAATA | GAAGGAACCGGGCATTTCC |
| FOXP3 | NM_014009.3 | TGTGGGGTAGCCATGGAAA | GGGTCGCATGTTGTGGAA |
| GC | NM_001204306. | GGCGTTCAGGAAAGATCCAA | ACAGAGGAGCTTGTCCGTAA |
| GCC2 | NM_181453.3 | GAGTGACAGCACTACAGGAA | CGGACTTTGTAGCTCTCGAA |
| GNLY | NM_006433.3 | GACCTGTCTGACGATAGTCCAA | GGTCGCAGCATTGGAAACA |
| GPR15 | NM_005290.N | GTGCAAAGGGAGCTCCTACA | TAGCGGTCAACACTCATGCA |
| GZMA | NM_006144.3 | GAAGCCTCCGAGGTGGAA | GAAAACACCCTCGCACAACA |
| GZMB | NM_004131.4 | CCCCATCCAGCCTATAATCCTAA | CTGGGCCTTGTTGCTAGGTA |
| GZMH | NM_033423.4 | CCCCTACATGGCCTTTGTTCA | TGAGCAGCTGTCAGCACAAA |
| GZMK | NM_002104.2 | CACATTTCATCTGGGCTTCTTAAA | GTGACACTTCTTTCCCTCCA |
| HAVCR2 | NM_032782.3 | GGATCCAAATCCCAGGCATAA | CTTGGAAAGGCTGCAGTGAA |
| HDAC2 | NR_033441.1 | ACCAGAACACTCCAGAATATATGGAA | ACACCAGGTGCATGAGGTAA |
| HLA-DMB | NM_002118.4 | TTCTGGGGATCACTGACCAA | CAGGCTCCCTCGTGTTAAAA |
| HNRNPA2B1 | NM_031243. | TTGAGCCAAAACGTGCTGTA | GCCAACAAACAGCTTCTTCAC |
| HNRNPDL | NM_031372.2 | GGAATTCGCAGAGGGATCCAA | CCCAGCTCAAGCCTCCAATAA |
| HNRNPM | NM_005968.N | TGGCCAGCTGCTATTTGATA | CTCAGGAGGGAAGAAATCTCC |
| ICAM1 | NM_000201.2 | CCCCTACCAGCTCCAGACC | TGCGTGTCCACCTCTAGGAC |
| IFI44L | NM_006820.N | GTTTTATGGCCACCGTCAGTA | CCTCTTTATGTCGTCTAGGTTATCC |
| IFITM2 | NM_006435.2 | ACTGAGAACCATCCCGGTAAC | CCGCTGTTGACAGGAGAGAA |
| IFNG | NM_000619.2 | ACTGCCAGGACCCATATGTAA | GTTCCATTATCCGCTACATCTGAA |
| IFNgR1 | NM_000416.2 | AAGCCAGGGTTGGACAAAA | GATATCCAGTTTAGGTGGTCCAA |
| IKZF1 | NM_006060.5 | ATACTCCAGATGAGGGCGATG | TCTTGGAGCTTTGCTGTCCT |
| IKZF4 | NM_001351089.N | AACAGCCAGCACTCTTCTCC | GCTTGACTCCTCATCGCTGTA |
| IL10 | NM_000572.2 | CCGTGGAGCAGGTGAAGAA | GTCAAACTCACTCATGGCTTTGTA |
| IL10RA | NM_001558.3 | CGCCGAAAGAAGCTACCC | CGCTGGCTGATGAAGATGAA |
| IL17A | NM_002190.2 | ACTACAACCGATCCACCTCAC | ACTTTGCCTCCCAGATCACA |
| IL17RA | NM_014339.4 | CCAAACCACCAGTCCAAGAA | CTCATGCATGGCGTGGTTA |
| IL1B | NM_000576.2 | GACCTGAGCACCTTCTTTCC | CGTGCACATAAGCCTCGTTA |
| IL2 | NM_000586.3 | ACCCAGGGACTTAATCAGCAA | GCATATTCACACATGAATGTTGTTTCA |

| | | | |
|---|---|---|---|
| IL2RA | NM_000417.2 | TCCTGGGACAACCAATGTCA | GTCACTTGTTTCGTTGTGTTCC |
| IL6 | NM_000600.3 | AGAGCTGTGCAGATGAGTACAA | GTTGGGTCAGGGGTGGTTA |
| IL6R | NM_000565.2 | GTAGTGTCGGGAGCAAGTTCA | ATGTTGGCAGGCGGATCA |
| IL7R | NM_002185.2 | GGAGAAAGTGGCTATGCTCAA | CTGCGATCCATTCACTTCCA |
| IRF8 | NM_002163.2 | TGGACATTTCCGAGCCATACA | AGCAGTTGCCACGCCTA |
| ISG15 | NM_005101.3 | CTGAGAGGCAGCGAACTCA | GCTCAGGGACACCTGGAA |
| ISM1 | NM_080826.N | GCGGGAAGCGAGGAGTTTAA | TCGCTTTTGCAGCTCATCCA |
| ITGA2 | NM_002203.3 | TGCCCCGAGCACATCATTTA | CTGATGTCCACACGCAAATCC |
| ITGA4 | NM_000885. | GCCAACGCTTCAGTGATCAA | CAAGTCTTTCCACAAGGTTCTCC |
| ITGA5 | NM_002205.2 | ACCCGAATTCTGGAGTATGCA | TGAAGCCTCCTTGGCAGTAA |
| ITGA6 | NM_000210.2 | ATATCAGGCTGCCAAATGCA | ACTCCCGAATACTGAGCTACA |
| ITGAL | NM_001114380.1 | CAACAGCAGCAAGCATTTCC | CTTCTCATACACCACGTCAACC |
| ITGB1 | NM_002211.3 | GAATGTATACAAGCAGGGCCAAA | CGTGCAGAAGTAGGCATTCC |
| ITGB2 | NM_000211.3 | TCAACGAGATCACCGAGTCC | CTTATCAGGGTGCGTGTTCAC |
| ITGB7 | NM_000889.1 | CAGCCCATCTTTGCTGTCAC | CCCCAACTGCAGACTTAGGAA |
| JAK1 | NM_002227.2 | CCCACAAACACATCGTGTACC | CCCCTTCCACAAACTCTTCCA |
| JUN | NM_002228.3 | AAGAACTCGGACCTCCTCAC | TGGATTATCAGGCGCTCCA |
| KIF2A | NM_001243952.N | TCGACGGACTGTAGCTTCTA | TGCACGTGCTGAACCAA |
| KLF6 | NM_001300.5 | TGCCGTCTCTGGAGGAGTAC | GGGCTCGCTCTGGAGGTA |
| KLRB1 | NM_002258.2 | GGACATTCAACAGAGCAGGAA | CTCGGAGTTGCTGCCAATATA |
| KLRG1 | NM_005810.3 | GCCTAGCCAGAGACTCACA | AATCCAGCAAAAGGCCTCAC |
| LAG3 | NM_002286.5 | TGGAGCCTTTGGCTTTCAC | GAGGGTGAATCCCTTGCTCTA |
| LAPTM4B | NM_018407.N | TGGGAGGTGACTTTGAGTTCA | AGCCATAGCACATATCAGGATCA |
| LYST | NM_000081.2 | AATCCTGAAGATGGCGAAACC | GGACTGCTTGATGCACTGAA |
| LYZ | NM_000239.2 | GGAGCAGTTAATGCCTGTCA | CCCTCTTTGCACAAGCTACA |
| MAF | NM_001031804.2 | TCGACGACCGCTTCTCC | ATCACCTCCTCCTTGCTGAC |
| MALT1 | NM_006785.N | TGAATTCAGCCAGTGGTCAC | TTCAGAGACGCCATCAACAC |
| MAPK7 | NM_139033.N | TTGTGACCCAGCAGCTATCTA | CTTTGGTGTGCCTGAGAACA |
| MIER2 | NM_017550.N | GGACTCTCTTCCTGCCAACAA | TTGCTGAGGTCAGCTTGGAA |
| MKI67 | NM_001145966.1 | AGAGTAACGCGGAGTGTCA | CTTGACACACACATTGTCCTCA |
| MTDH | NM_001363137.N | ACTAGTGATCCAGCCGAAGTA | ATGGCTCGGTAGAAGTAGCA |
| NAB1 | NM_005966.3 | GGAAGGATGGGAAACATCTCAC | GCAGGGCATTATCCTTCACA |
| NDUFA1 | NM_004541. | GATAGGCGCATCTCTGGAGT | TCATCAATCAGGAAAATGCTTCCTT |
| NDUFA11 | NM_001193375.1 | AGCCTACAGCACCACCAGTA | CCCGGAGGATTGAGTGTGAC |
| NEDD4L | NM_001144967.N | GAATTCACTTGGATGGCCGAAC | TTCTGCAGTCTTGGGTCTTCC |
| NFE2L3 | NM_004289.N | CCTGACTGGGAGGCAGAAAA | GTCTTCCAGAGAGAAAGAAGTATCTGT |
| NFKBIA | NM_020529.2 | CACCTCCACTCCATCCTGAA | GGTAGCCATGGATAGAGGCTAA |
| NOTCH2NLC | NM_001364012.N | AGAGACTATGCGTGGCACAA | TGGGTTGTGTGTGCACTCTA |
| NUCKS1 | NM_022731.N | ATGCCCAAACCCAGACTAAA | CTTTTCTGGCGGGCTTTCC |
| OPTN | NM_001008211.N | AGCTGGCAGTTCTGCTGAAA | GACGACTCTGCATCTCCATCAA |
| PABPC4 | NM_001135653.N | GCTATGCCTACGTCAACTTCC | GCGGATTGGCTTTCCCTTAA |
| PAIP2 | NM_016480.3 | GCAGAGTACATGTGGATGGAA | CATTTCTTGGAAACAGCGTTCA |
| PAK2 | NM_002577.N | CTACGACTCCAACACAGTGAA | GCTGGTGTTCCAGAAGGAA |
| PCNA | NM_182649.1 | TCTGAGGGCTTCGACACCTA | CATTGCCGGCGCATTTTAGTA |
| PDCD1 | NM_005018.2 | GCAGCCTGGTGCTGCTA | GTGCGCCTGGCTCCTA |
| PDCD6IP | NM_001162429.N | TTGTTCTATTGAACCCAAATTCCC | CGAAAGCATCCTTCCAGGTA |

| | | | |
|---|---|---|---|
| PECAM1 | NM_000442. | ATGCCGTGGAAAGCAGATA | CATCTGGCCTTGCTGTCTA |
| PIK3IP1-AS1 | NR_110542.N | TGCATCCATCCTGCCATCTG | GCAGAGGCAGTTCCGTTCT |
| PRDX5 | NM_012094.N | GTTCGGCTCCTGGCTGAT | CAAAGATGGACACCAGCGAATC |
| PRKDC | NM_006904.N | GCTTGGATCTCTAGGAGGACAA | AGGCCACATAGCTCTTCATCA |
| PRPF40A | NM_017892.3 | AACAAGCTGCTCCTCCGATA | TCCTCAAATGCTGGCTCTTTTAC |
| PSMB3 | NM_002795.2 | ATGGTGGCCAACCTCTTGTA | CTTAAAGGTCTTCGGGTCCAAC |
| PSMB9 | NM_002800.N | TTGTGATGGGTTCTGATTCCC | GGGGACAGCTTGTCAAACA |
| PSME1 | NM_006263.2 | CGCTTGAAGCCTGAGATCA | ACCATCCTCAATCCGAGGTA |
| PTPRC | NM_002838.4 | GTGGCTTAAACTCTTGGCATTT | GGGAAGGTGTTGGGCTTT |
| RAP1A | NM_001010935.1 | CCAACAGTGTATGCTCGAAATCC | CAAAACCTTGGCCGTTCTTCA |
| RICTOR | NM_152756.3 | CTTCCGTGTCGGAGGTTCATA | ACACAGCCTCTGCTTCTTCA |
| RNF213 | NM_020954.2 | AGGAAGCAGATGTCCAGGAA | AGAGAGATGATGGCGTGGAA |
| RORC | NM_001001523.1 | CAAGACTCATCGCCAAAGCA | TTTCCACATGCTGGCTACAC |
| RSF1 | NM_016578.N | TGGCAAATCTGTTACTGCAGACA | TCCCATGCCCAGGTACTGTTA |
| RTKN2 | NM_145307.N | AAGAACGAACAGCATGTAAAGGAA | GCTGAAGTGATCAGAGTCTTTCC |
| RXRA | NM_002957.4 | AGGAAACATGGCTTCCTTCAC | TCGCAGCTGTACACTCCATA |
| S100A4 | NM_002961.2 | AGGGTGACAAGTTCAAGCTCAA | TTGTCCCTGTTGCTGTCCAA |
| S100A6 | NM_014624.3 | AAGCACACCCTGAGCAAGAA | AGCCTTGCAATTTCAGCATCC |
| SEC11A | NM_001271918. | TCATCGGCACTAATGATCTGGAA | CACCACTACAATCGGACTTTCAC |
| SELL | NM_000655.4 | CACCTGTGGACCATTTGGAA | GTGCTGATAGAGGCTCACAC |
| SELPLG | NM_001206609.1 | AGAAACGGAGCCTCCAGAAA | TGGTAGACTCAGGGGTTCCA |
| SET | NM_001122821.1 | GGTGATCCATCTTCGAAGTCC | TGACTCGAACGTTTCGTCAA |
| SNHG14 | NR_146177.N | AACATGTTTGGCATGGTCTCC | ACAGACACAGTCCTTCCTTCC |
| SNRPB | NM_003091. | AGGGGAGAATCTGGTCTCAA | AGTGGAACTCGAGCAATACC |
| SNRPN | NM_003097.3 | AGCCCCAACACAGTACCC | TACCAGGTGGAGGAGCCATA |
| SP100 | NM_001080391.1 | CTTCATCTCAGCACCGAGAA | CTTGGCCCTCCTTTTCATCA |
| SRI | NM_003130.2 | ACACTGACAGGAGTGGAACA | GCCTGGGGACTCAACCTAA |
| STAT3 | NM_003150.3 | GGAAATAATGGTGAAGGTGCTGAAC | CCGAGGTCAACTCCATGTCAAA |
| STAT4 | NM_003151.2 | CAGTGCTGGAGGTAAAGGAA | AGAGGCAGATCTGTGTTTCAA |
| STAT5A | NM_003152.3 | CCCAGGCTCCCTATAACATGTA | ATGGTCTCATCCAGGTCGAA |
| STK17A | NM_004760.2 | CTGAGTCGGCTGTTGATTTCA | AACCAGGGGTGCTTTAGACA |
| SUB1 | NM_006713.3 | CAGCAGCAGCAGAGATGATAA | ATCGCGAACACTAACGTACC |
| TAOK3 | NM_016281.3 | CCAAGAAGCAAGTGGCTATCA | GCCAAGATCTGTTGCTGGAA |
| TBX21 | NM_013351.1 | GGGCGTCCAACAATGTGAC | CCGTCGTTCACCTCAACGATA |
| TGFB1 | NM_000660.4 | CGTCTGCTGAGGCTCAAGTTA | TCGCCAGGAATTGTTGCTGTA |
| TGFbR1 | NM_004612.2 | GAAATTGCTCGACGATGTTCC | ACTGATGGGTCAGAAGGTACA |
| TGOLN2 | NM_001206840. | TTGCCTTGGTCCTCCTGAAC | TTCCTGCAGAAGGCCGTAC |
| THEMIS | NM_001164685.N | CCCTGATTCTCAAGCCTGTTTA | AGCTGAAGAAACCAGTTAGCA |
| TIGIT | NM_173799.3 | GTGGTGGTCGCGTTGACTA | TCCTGTCCAGCTGATTTTCTCC |
| TLN1 | NM_006289.3 | GCTGGGAAAGCTTTGGACTAC | TTCAGGGGTCTCTGTTTCTTCC |
| TNF | NM_000594.2 | CCCAGGGACCTCTCTCTAATCA | ATGGGCTACAGGCTTGTCAC |
| TNFRSF18 | NM_148901.1 | GCTGCTGCCGCGATTA | GAATTCAGGCTGGACACACA |
| TNFRSF9 | NM_001561.4 | GGGGCAGAAAGAAACTCCTGTA | TCTGGAAATCGGCAGCTACA |
| TRIM22 | NM_006074.N | AGCTGACAGATGTCCAGTAC | ATGTGCAGGTGCGTACA |
| TRUB1 | NM_139169.N | GTTGGAATTGGAAGCGGAAC | TAGCTTTCCCCAGTTCTCCA |
| TUBA1A | NM_001270399. | GCCAAACGTGCCTTTGTTCA | CACGGGCCTCTGAAAACTCA |

| | | | |
|---|---|---|---|
| UBXN1 | NM_015853.4 | AGATCGAGAGGGACAAAGCA | GCTGGGAGAAGAGGGAACA |
| UCKL1-AS1 | NR_027287.N | CCACAATGCCAAGGAAACCA | TCCTCAGTGTCGGTTTCACA |
| UXT | NM_153477.N | CTCACCAAGGACTCCATGAA | GCAGGCCTTGTAGTTCTCTA |
| VDR | NM_000376.2 | ATAAGACCTACGACCCCACCTA | GCTCCCTCCACCATCATTCA |
| XIST | NR_001564.2 | TTGGATGGGTTGCCAGCTA | TCTCCACCTAGGGATCGTCAA |
| YPEL3 | NM_031477.4 | GGATGATTGTCACCGGAGGTA | CCCTGACTGCCCTGGAA |
| ZFP36 | NM_003407.3 | CCTCATGGCCAACCGTTACA | AGCCTGGGCTGGACTCA |

**Table Sup 2. Sequences of PCR Primers used for validation step by HT-qPCR using Biomark HD.**

| GENES | **Memory B** | **Naive B** | **Naive CD4** | **Naive CD8** | **MAIT** | **Th17** | **Th1** | **Treg** | Th2 | Tfh | NK | NKT | CD8 CM | CD8 EM | CD8 EMRA+ | gdTcells |
|---|---|---|---|---|---|---|---|---|---|---|---|---|---|---|---|---|
| INPP4B* | -2,944 | | | | | | | | | | | | | | | |
| CAHM* | -2,851 | | | | | | | | | | | | | | | |
| AC012076.1* | -2,692 | | | | | | | | | | | | | | | |
| HTR3A* | -2,637 | | | | | | | | | | | | | | | |
| GPD1L* | -2,525 | | | | | | | | | | | | | | | |
| C20orf194* | -2,508 | | | | | | | | | | | | | | | |
| LINC02018* | -2,468 | 2,286 | | | | | | | | | | | | | | |
| BBS1* | -2,412 | | | | | | | | | | | | | | | |
| SLC38A11* | -2,361 | | | | | | | | | | | | | | | |
| AL020997.2* | -2,235 | -1,726 | | | | | | | | | | | | | | |
| AC104024.1* | -2,181 | | | | | | | | | | | | | | | |
| AL645728.1* | -2,123 | | -1,057 | | | | | | | | | | | | | |
| SENP3* | -2,054 | | | | | | | | | | | | | | | |
| SLC17A5* | -2,016 | | | | | | | | | | | | | | | |
| MDC1* | -2,011 | | | | | | | | | | | | | | | |
| **FOS*** | -1,981 | -2,328 | -1,684 | | | | | | | | | | | | | |
| GGT7* | -1,909 | | | | | | | | | | | | | | | |
| AC124016.2* | -1,806 | | | | | | | | | | | | | | | |
| LIN54* | -1,713 | | | | | | | | | | | | | | | |
| AL450998.2* | -1,696 | | | | | | | | | | | | | | | |
| C5orf22* | -1,662 | | | | | | | | | | | | | | | |
| GLE1* | -1,644 | | | | | | | | | | | | | | | |
| ZNF347* | -1,622 | | | | | | | | | | | | | | | |
| CXorf40B* | -1,610 | | | | | | | | | | | | | | | |
| URB2* | -1,602 | | | | | | | | | | | | | | | |
| MIS18A* | -1,580 | | | | | | | | | | | | | | | |
| DPYD* | -1,569 | | | | | | | | | | | | | | | |
| VPS9D1* | -1,563 | | | | | | | | | | | | | | | |
| AL353622.1* | -1,554 | | | | | | | | | | | | | | | |
| SPATA20* | -1,491 | | | | | | | | | | | | | | | |
| TRIM65* | -1,453 | | | | | | | | | | | | | | | |
| NIT1* | -1,413 | | | | | | | | | | | | | | | |
| ATP8B2* | -1,408 | | | | | | | | | | | | | | | |
| IMMP2L* | -1,352 | | | | | | | | | | | | | | | |
| DNASE1L1* | -1,351 | | | | | | | | | | | | | | | |

| GENES | **Memory B** | **Naive B** | **Naive CD4** | **Naive CD8** | **MAIT** | **Th17** | **Th1** | **Treg** | Th2 | Tfh | NK | NKT | CD8 CM | CD8 EM | CD8 EMRA+ | gdTcells |
|---|---|---|---|---|---|---|---|---|---|---|---|---|---|---|---|---|
| UTP20* | -1,347 | | | | | | | | | | | | | | | |
| **CD69*** | -1,346 | -1,244 | | | | | | | | | | | | | | |
| TRAF6* | -1,341 | | | | | | | | | | | | | | | |
| MPP5* | -1,320 | | | | | | | | | | | | | | | |
| TNFRSF10B* | -1,319 | | | | | | | | | | | | | | | |
| NPTN* | -1,290 | | | | | | | | | | | | | | | |
| **FOSB*** | -1,281 | | -1,575 | -1,547 | | | -0,842 | | | | | | | | | |
| ADARB1* | -1,236 | | | | | | | | | | | | | | | |
| C2orf69* | -1,151 | | | | | | | | | | | | | | | |
| MYC* | -1,103 | | | | | | | | | | | | | | | |
| ODR4* | -1,068 | | | | | | | | | | | | | | | |
| APPBP2* | -1,066 | | | | | | | | | | | | | | | |
| KNOP1* | -1,048 | | | | | | | | | | | | | | | |
| AC044849.1* | -1,046 | -1,173 | | | | | | | | | | | | | | |
| EHMT2* | -1,022 | | | | | | | | | | | | | | | |
| MCM7* | -0,976 | | | | | | | | | | | | | | | |
| CEP295* | -0,959 | | | | | | | | | | | | | | | |
| ZC3H6* | -0,944 | | | | | | | | | | | | | | | |
| PEA15* | -0,898 | | | | | | -1,778 | | | | | | | | | |
| PPP1R15A* | -0,889 | | -1,376 | -1,428 | | | | | | | | | | | | |
| ZMYM4* | -0,888 | | | | | | | | | | | | | | | |
| JOSD1* | -0,880 | | | | | | | | | | | | | | | |
| PRXL2C* | -0,864 | | | | | | | | | | | | | | | |
| DDX50* | -0,818 | | | | | | | | | | | | | | | |
| FAM172A* | -0,790 | | | | | | | | | | | | | | | |
| JUNB* | -0,782 | | | | | | | | | | | | | | | |
| BLOC1S6* | -0,756 | | | | | | | | | | | | | | | |
| GNL1* | -0,750 | | | | | | | | | | | | | | | |
| NARS* | -0,744 | | | | | | | | | | | | | | | |
| RNF167* | -0,731 | | | | | | | | | | | | | | | |
| OTUD1* | -0,709 | | | | | | | | | | | | | | | |
| BACH2* | -0,682 | | | | | | | | | | | | | | | |
| GNA13* | -0,666 | | | | | | | | | | | | | | | |
| CYSLTR1* | -0,586 | | | | | | | | | | | | | | | |
| PDCD10* | -0,540 | | | | | | | | | | | | | | | |

| GENES | **Memory B** | **Naive B** | **Naive CD4** | **Naive CD8** | **MAIT** | **Th17** | **Th1** | **Treg** | Th2 | Tfh | NK | NKT | CD8 CM | CD8 EM | CD8 EMRA + | gdTcells |
|---|---|---|---|---|---|---|---|---|---|---|---|---|---|---|---|---|
| SERBP1* | 0,335 | | | | | | | | | | | | | | | |
| ISG20* | 0,452 | | | | | | | | | | | | | | | |
| CPNE5* | 0,494 | | | | | | | | | | | | | | | |
| PSMB4* | 0,525 | | | | | | | | | | | | | | | |
| **JUN** | 0,594 | 0,523/-1,091* | 0,403/-1,468* | | -1,646* | | -1,383* | | | | | 0,469 | | | | |
| WAS* | 0,594 | | | | | | | | | | | | | | | |
| MPG* | 0,608 | | | | | | | | | | | | | | | |
| MHENCR* | 0,629 | | | | 1,245 | | | | | | | | | | | |
| RAB14* | 0,629 | | | | | | | | | | | | | | | |
| PSMD7* | 0,641 | | | | | | | | | | | | | | | |
| THUMPD3-AS1* | 0,660 | | | | | | | | | | | | | | | |
| ANXA4* | 0,679 | | | | | | | | | | | | | | | |
| HSH2D* | 0,679 | | | | | | | | | | | | | | | |
| **PDCD6IP** | 0,718 | | | | | | | | | | | | | | | |
| WIPF1 | 0,726 | | | | | | | | | | | | | | | |
| GTF2B* | 0,815 | | | | | | | | | | | | | | | |
| **UXT** | 0,828 | | | | | 1,210 | | | | | | | | | | |
| CTSC* | 0,838 | | | | | | | | | | | | | | | |
| SLC35A3* | 0,845 | | | | | | | | | | | | | | | |
| **HNRNPDL** | 0,865 | | 0,827 | 0,840 | | | | | | | | | | | | |
| HIST1H1D* | 0,911 | 0,866 | | | -0,776 | | | | | | | | | | | |
| TYMP* | 0,936 | 0,958 | | | | | 1,477 | | | | | | | | | |
| TIMM8B* | 0,954 | | | | | | | | | | | | | | | |
| MAU2* | 0,962 | | | | | | | | | | | | | | | |
| POP4* | 0,971 | | | | | | | | | | | | | | | |
| FBXO21* | 1,020 | | | | | | | | | | | | | | | |
| PARP9* | 1,074 | 1,000 | | | | 0,933 | | | | | | | | | | |
| RGP1* | 1,098 | | | | | | | | | | | | | | | |
| ARMH3* | 1,141 | | | | | | | | | | | | | | | |
| OAS1* | 1,142 | | | | | | | | | | | | | | | |
| TP53BP1* | 1,165 | | | | | | | | | | | | | | | |
| IFI35* | 1,168 | | | | | | | | | | | | | | | |
| IGHA1 | 1,168 | | | | | | | | | | | | | | | |
| MLEC* | 1,187 | | | | | | | | | | | | | | | |

| GENES | **Memory B** | **Naive B** | **Naive CD4** | **Naive CD8** | **MAIT** | **Th17** | **Th1** | **Treg** | Th2 | Tfh | NK | NKT | CD8 CM | CD8 EM | CD8 EMRA+ | gdTcells |
|---|---|---|---|---|---|---|---|---|---|---|---|---|---|---|---|---|
| BDP1 | 1,204 | | | | | | | | | | | | | | | |
| CDKN2D | 1,219 | | | | | | | | | | | 1,508 | | | | |
| **HLA.DMB** | 1,250 | | | | | | | | | | | | | | | |
| DPYSL2* | 1,289 | | | | | | | | | | | | | | | |
| KLHL14* | 1,289 | | | | | | | | | | | | | | | |
| AC007384.1* | 1,315 | | | | | | | | | | | | | | | |
| CD68* | 1,332 | | | | | | | | | | | | | | | |
| C9orf78 | 1,337 | | | | 1,531 | | | | | | | 1,489 | | | | |
| **SNRPB** | 1,343 | | | | | | | | | | | | | | | |
| TTBK2* | 1,344 | | | | | | | | | | | | | | | |
| ITFG2-AS1* | 1,357 | | | | | | | | | | | | | | | |
| SOX4* | 1,363 | | | | | | | | | | | | | | | |
| RBBP8* | 1,553 | | | | | | | | | | | | | | | |
| APOL6* | 1,631 | | | | | | | | | | | | | | | |
| CASP6* | 1,650 | | | | | | | | | | | | | | | |
| AC024060.2* | 1,654 | | | | | | | | | | | | | | | |
| METTL2A* | 1,698 | | | | | | | | | | | | | | | |
| ZNF484* | 1,949 | | | | | | | | | | | | | | | |
| C5* | 2,173 | | | | | | | | | | | | | | | |
| EPB41L4A* | 2,186 | | | | | | | | | | | | | | | |
| ARMCX5* | 2,289 | | | | | | | | | | | | | | | |
| AL079338.1* | 2,644 | | | | | | | | | | | | | | | |
| SELENOW/* | 0,514* | | | 0,832 | | | | | | | | | | | | |
| PSME2/* | 0,533* | 0,594* | | | | | | 1,330 | | | | | | | | |
| **ZFP36/*** | 0,696-0,540* | | | | | | | | | | | | | | | |
| **IRF8/*** | 0,758/-0,462* | | | | | | | | | | | | | | | |
| DGKZ/* | -0,98* | | | | | | | | 1,491 | | | | | | | |
| **PSMB3/*** | 1,428/0,431* | | | | | 1,224 | | 1,503 | | | | | | | | |
| **S100A6/*** | 1,522/0,518* | | | | | | | | | | | | | | | |
| **S100A4/*** | 1,655/0,734* | | | | | | | | | | | | | | | |

| GENES | **Memory B** | **Naive B** | **Naive CD4** | **Naive CD8** | **MAIT** | **Th17** | **Th1** | **Treg** | Th2 | Tfh | NK | NKT | CD8 CM | CD8 EM | CD8 EMRA + | gdTcells |
|---|---|---|---|---|---|---|---|---|---|---|---|---|---|---|---|---|
| IGLC7* | | -2,710 | | | | | | | | | | | | | | |
| AC127035.1* | | -2,706 | | | | | | | | | | | | | | |
| DZANK1* | | -2,532 | | | | | | | | | | | | | | |
| TNFRSF25* | | -2,348 | | | | | | | | | | | | | | |
| FBXO5* | | -2,190 | | | | | | | | | | | | | | |
| DNAJB4* | | -2,018 | | | | | | | | | | | | | | |
| MMUT* | | -2,007 | | | | | | | | | | | | | | |
| FSD1L* | | -1,970 | | | | | | | | | | | | | | |
| AC005838.2* | | -1,880 | | | | | | | | | | | | | | |
| ORC6* | | -1,843 | | | | | | | | | | | | | | |
| IRF5* | | -1,821 | | | | | | | | | | | | | | |
| CRY1* | | -1,763 | | | | | | | | | | | | | | |
| MAD2L1* | | -1,750 | | | | | | | | | | | | | | |
| TNPO2* | | -1,718 | | | | | | | | | | | | | | |
| AC073896.2* | | -1,710 | | | | | | | | | | | | | | |
| LINC01534* | | -1,651 | | | | | | | | | | | | | | |
| ALKBH1* | | -1,635 | | | | | | | | | | | | | | |
| BUD13* | | -1,610 | | | | | | | | | | | | | | |
| FASTKD5* | | -1,605 | | | | | | | | | | | | | | |
| AL121603.2* | | -1,604 | | | | | | | | | | | | | | |
| ACAD11* | | -1,549 | | | | | | | | | | | | | | |
| TRMT12* | | -1,499 | | | | | | | | | | | | | | |
| EHD4* | | -1,469 | | | | -0,973 | | | | | | | | | | |
| TRIM41* | | -1,430 | | | | | | | | | | | | | | |
| ATG4A* | | -1,430 | | | | | | | | | | | | | | |
| FTO* | | -1,426 | | | | | | | | | | | | | | |
| ABCB1* | | -1,376 | | | | | | | | | | | | | | |
| SOCS5* | | -1,361 | | | | | | | | | | | | | | |
| REXO1* | | -1,292 | | | | | | | | | | | | | | |
| EXD3* | | -1,260 | | | | | | | | | | | | | | |
| FAN1* | | -1,259 | | | | | | | | | | | | | | |
| ARHGAP32* | | -1,234 | | | | | | | | | | | | | | |
| SREBF2-AS1* | | -1,234 | -1,243 | | | | | | | | | | | | | |
| LNPK* | | -1,215 | | | | | | | | | | | | | | |
| NUP155* | | -1,199 | | | | | | | | | | | | | | |

| GENES | **Memory B** | **Naive B** | **Naive CD4** | **Naive CD8** | **MAIT** | **Th17** | **Th1** | **Treg** | Th2 | Tfh | NK | NKT | CD8 CM | CD8 EM | CD8 EMRA+ | gdTcells |
|---|---|---|---|---|---|---|---|---|---|---|---|---|---|---|---|---|
| R3HCC1L* | | -1,134 | | | | | | | | | | | | | | |
| ORC3* | | -1,120 | | | | | | | | | | | | | | |
| PPP2CB* | | -1,118 | | | | | | | | | | | | | | |
| CSTF2T* | | -1,097 | | | | | | | | | | | | | | |
| DEDD2* | | -1,070 | | | | | | | | | | | | | | |
| SAMD4B* | | -1,024 | | | | | | | | | | | | | | |
| TUBB4B* | | -1,009 | | | | | | | | | | | | | | |
| LIMS1* | | -0,963 | | | | | | | | | | | | | | |
| TMEM260* | | -0,962 | | | | | | | | | | | | | | |
| RAB11FIP1* | | -0,912 | | | | | | | | | | | | | | |
| MPHOSPH9* | | -0,910 | | | | | | | | | | | | | | |
| GTF2H5* | | -0,782 | | | | | | | | | | | | | | |
| NBPF19* | | -0,767 | | | | | | | | | | | | | | |
| HDAC8* | | -0,753 | | | | | | | | | | | | | | |
| SIN3A* | | -0,699 | | | | | | | | | | | | | | |
| TMEM179B* | | -0,695 | | | | | | | | | | | | | | |
| DENND4B* | | -0,687 | | | | | | | | | | | | | | |
| SUN2* | | -0,676 | | | | | | | | | | | | | | |
| LIMD1* | | -0,655 | | | | | | | | | | | | | | |
| SERPINB9P1* | | -0,653 | | | | | | | | | | | | | | |
| NLRP1* | | -0,649 | | | | | | | | | | | | | | |
| ITPR2* | | -0,599 | | | | | | | | | | | | | | |
| SAT2* | | -0,586 | | | | | | | | | | | | | | |
| AHNAK/* | | -0,550 | | | | | | | | 0,717 | | | | | | |
| VDAC2* | | -0,457 | | | | | | | | | | | | | | |
| PNRC1 | | -0,314 | | | -0,550 | | | | | | | | | | | 0,615 |
| HLA-C* | | 0,332 | | | | | | | | | | | | | | |
| DNAJC10* | | 0,446 | | | | | | | | | | | | | | |
| HHEX* | | 0,478 | | | | | | | | | | | | | | |
| TRIM56* | | 0,498 | | | | 0,649 | | | | | | | | | | |
| PARP14* | | 0,528 | | | | | | | | | | | | | | |
| RESF1/* | | 0,530 | 0,380 | | | 1,244 | | | | | | | | | | |
| BST2* | | 0,548 | | | | | | | | | | | | | | |
| SNRPC* | | 0,594 | | | | | | | | | | | | | | |
| C17orf49* | | 0,605 | | | | | | | | | | | | | | |

| GENES | **Memory B** | **Naive B** | **Naive CD4** | **Naive CD8** | **MAIT** | **Th17** | **Th1** | **Treg** | Th2 | Tfh | NK | NKT | CD8 CM | CD8 EM | CD8 EMRA+ | gdTcells |
|---|---|---|---|---|---|---|---|---|---|---|---|---|---|---|---|---|
| AC004687.1/* | | 0,629 | | | | | | | | | | | 1,810 | | 2,150 | |
| STMN3* | | 0,631 | | | | | | | | | | | | | | |
| DENND11* | | 0,634 | | | | | | | | | | | | | | |
| RAB18* | | 0,653 | | | | | | | | | | | | | | |
| LPP* | | 0,668 | | | | | | | | | | | | | | |
| CXCR5* | | 0,682 | | | | | | | | | | | | | | |
| SAMD9* | | 0,686 | | | | | | | | | | | | | | |
| PSMD2* | | 0,696 | | | | | | | | | | | | | | |
| IRF9* | | 0,737 | | | | 0,908 | | | | | | | | | | |
| ZBED1* | | 0,738 | | | | | | | | | | | | | | |
| ZNF292 | | 0,755 | | | | | | | | | | | | | | 1,630 |
| NUAK2* | | 0,759 | | | | | | | | | | | | | | |
| PWWP3A* | | 0,759 | | | | | | | | | | | | | | |
| **GCC2** | | 0,772 | 0,891 | | | | | | | | | | | | | |
| UROS* | | 0,776 | | | | | | | | | | | | | | |
| TBCC* | | 0,792 | | | | | | | | | | | | | | |
| USP14* | | 0,797 | | | | | | | | | | | | | | |
| **LYST** | | 0,805 | | | | | | | | | | | | | | |
| **DGKD** | | 0,806 | | | | | | | | | | | | | | |
| ADD1 | | 0,825 | | | | | | | | | | 1,548 | | | | |
| CTSZ | | 0,864 | | | | | | | | | | | | | | |
| BOD1L1 | | 0,874 | | | 1,420 | | | | | | | | | | | |
| **CHD2** | | 0,899 | | | | | | | | | | | | | | |
| PIK3CA* | | 0,905 | | | | | | | | | | | | | | |
| PLSCR1* | | 0,913 | | | | | | | | | | | | | | |
| ARRDC3* | | 0,953 | | 0,873 | | | | | | | | | | | | |
| GNLY* | | 1,060 | | | | -1,955 | | | | | | | | | | |
| UNC45A* | | 1,067 | | | | | | | | | | | | | | |
| MBD4 | | 1,122 | | | | | | | | | | | | | | |
| STAT1* | | 1,126 | 0,691 | 1,062 | | 0,768 | | | | | | | | | | |
| N4BP2L2 | | 1,155 | | | | | | | | | | | | | | |
| KDM3A* | | 1,315 | | | | | | | | | | | | | | |
| AC010618.3* | | 1,380 | | | | | | | | | | | | | | |
| WARS* | | 1,411 | | | | | | | | | | | | | | |
| MAGEH1* | | 1,444 | | | | | | | | | | | | | | |

| GENES | **Memory B** | **Naive B** | **Naive CD4** | **Naive CD8** | **MAIT** | **Th17** | **Th1** | **Treg** | Th2 | Tfh | NK | NKT | CD8 CM | CD8 EM | CD8 EMRA+ | gdTcells |
|---|---|---|---|---|---|---|---|---|---|---|---|---|---|---|---|---|
| GK5* | | 1,520 | | | | | | | | | | | | | | |
| C6orf120* | | 1,524 | | | | | | | | | | | | | | |
| RNF14* | | 1,579 | | | | | | | | | | | | | | |
| NACC1* | | 1,627 | | | | | | | | | | | | | | |
| ZNF805* | | 1,719 | | | | | | | | | | | | | | |
| CD3G/* | | 1,941 | | | | | | | 1,245 | | | | | | | |
| AL360270.1* | | 2,025 | | | | | | | | | | | | | | |
| ZNF257* | | 2,144 | | | | | | | | | | | | | | |
| IGIP* | | 2,311 | | | | | | | | | | | | | | |
| CD226* | | 2,422 | | | | | | | | | | | | | | |
| IFIT3* | | 2,480 | | | | | | | | | | | | | | |
| HAUS7* | | 2,525 | | | | 2,148 | | | | | | | | | | |
| TNKS2-AS1* | | 2,571 | | | | | | | | | | | | | | |
| AC002470.1* | | 2,680 | | | | | | | | | | | | | | |
| AC104971.1* | | 2,689 | | | | | | | | | | | | | | |
| AL136419.3* | | 2,734 | | | | | | | | | | | | | | |
| AC018809.1* | | 2,819 | | | | | | | | | | | | | | |
| **CLEC2D/*** | | 0,343* | | 1,276/0,300* | | | | | | | | | | | | |
| **DUSP1/*** | | 0,446/-1,174* | | | | | | | | | | | | | | |
| **TUBA1A/*** | | 0,638/-0,773* | -0,525 | 0,571/-0,876* | | -0,465* | | | | | | | | | | |
| B2M/* | | 1,182 / 0,267 | | | | | | 1,165 | | | | | | | | |
| **SET/*** | | 1,215/0,265* | | | | | | | | | | | | | | |
| KLF2/* | | 1,226/0,304* | | | | | | 0,477* | | | | | | | | |
| **PSME1/*** | | 1,242/0,351* | | | | | | | | | | | | | | |
| **CAST/*** | | 1,333/0,433 | | | | | | | | | | | | | | |
| IGLC2/* | | 1,379/0,482* | | | | | | | | | | | | | | |

| GENES | **Memory B** | **Naive B** | **Naive CD4** | **Naive CD8** | **MAIT** | **Th17** | **Th1** | **Treg** | Th2 | Tfh | NK | NKT | CD8 CM | CD8 EM | CD8 EMRA+ | gdTcells |
|---|---|---|---|---|---|---|---|---|---|---|---|---|---|---|---|---|
| ANKRD37* | | | -3,206 | | | | | | | | | | | | | |
| C4orf46* | | | -2,830 | | | | | | | | | | | | | |
| NCAPG2* | | | -2,809 | | | | | | | | | | | | | |
| DEPDC7* | | | -2,728 | | | | | | | | | | | | | |
| AC010896.1* | | | -2,507 | | | | | | | | | | | | | |
| FLVCR1-DT* | | | -2,499 | | | | | | | | | | | | | |
| AL353743.2* | | | -2,469 | | | | | | | | | | | | | |
| MKLN1-AS* | | | -2,362 | | | | | | | | | | | | | |
| AC104389.5* | | | -2,309 | | | | | | | | | | | | | |
| SLC25A20* | | | -2,274 | | | | | | | | | | | | | |
| **MAPK7*** | | | -2,136 | | | | | | | | | | | | | |
| AL139330.1* | | | -2,107 | | | | | | | | | | | | | |
| RPS6KL1* | | | -2,076 | | | | | | | | | | | | | |
| ASPHD2* | | | -2,008 | | | | | | | | | | | | | |
| AL355581.1* | | | -1,830 | | | | | | | | | | | | | |
| AC018638.7* | | | -1,803 | | | | | | | | | | | | | |
| AC020978.7* | | | -1,653 | | | | | | | | | | | | | |
| TMEM147-AS1* | | | -1,651 | | | | | | | | | | | | | |
| SPRYD4* | | | -1,645 | | | | | | | | | | | | | |
| SLC22A18* | | | -1,622 | | | | | | | | | | | | | |
| **CD302*** | | | -1,607 | | | | | | | | | | | | | |
| ADD2* | | | -1,605 | | | | | | | | | | | | | |
| IER5* | | | -1,592 | | -1,685 | | -1,051 | | | | | | | | | |
| ACTR5* | | | -1,560 | | | | | | | | | | | | | |
| IFT22* | | | -1,495 | | | | | | | | | | | | | |
| ADPRHL2* | | | -1,408 | | | | | | | | | | | | | |
| PEX5* | | | -1,397 | | | | | | | | | | | | | |
| DENND1A* | | | -1,298 | | | | | | | | | | | | | |
| ISOC1* | | | -1,288 | | | | | | | | | | | | | |
| KLHL22* | | | -1,282 | | | | | | | | | | | | | |
| MXD1* | | | -1,269 | | | | | | | | | | | | | |
| CBR3-AS1* | | | -1,263 | | | | | | | | | | | | | |
| MOSPD1* | | | -1,193 | | | | | | | | | | | | | |
| RABIF* | | | -1,137 | | | | | | | | | | | | | |
| SEPTIN11* | | | -1,094 | | | | | | | | | | | | | |

| GENES | **Memory B** | **Naive B** | **Naive CD4** | **Naive CD8** | **MAIT** | **Th17** | **Th1** | **Treg** | Th2 | Tfh | NK | NKT | CD8 CM | CD8 EM | CD8 EMRA+ | gdTcells |
|---|---|---|---|---|---|---|---|---|---|---|---|---|---|---|---|---|
| NXT2* | | | -1,071 | | | | | | | | | | | | | |
| PI4K2B* | | | -1,053 | | | | | | | | | | | | | |
| NRBF2* | | | -1,037 | | | | | | | | | | | | | |
| DDX41* | | | -1,028 | -1,270 | | | | | | | | | | | | |
| CBWD2* | | | -1,026 | | | | | | | | | | | | | |
| HAUS2* | | | -1,020 | | | | | | | | | | | | | |
| FBLN5* | | | -1,012 | | | | | | | | | | | | | |
| SOCS2* | | | -0,956 | | | | | | | | | | | | | |
| AC009061.2* | | | -0,956 | | | | | | | | | | | | | |
| PIK3CG* | | | -0,946 | | | | | | | | | | | | | |
| SEH1L* | | | -0,942 | | | | | | | | | | | | | |
| TXNDC17* | | | -0,942 | | | | | | | | | | | | | |
| SPIN2B* | | | -0,920 | | | | | | | | | | | | | |
| ASPSCR1* | | | -0,920 | | | | 1,245 | | | | | | | | | |
| POLR3F* | | | -0,903 | | | | | | | | | | | | | |
| ETFDH* | | | -0,900 | | -2,172 | | | | | | | | | | | |
| HSPBAP1* | | | -0,894 | | | | | | | | | | | | | |
| ODC1* | | | -0,880 | -0,774 | | | | | | | | | | | | |
| PPP1R21* | | | -0,823 | | | | | | | | | | | | | |
| RCC1L* | | | -0,821 | | | | | | | | | | | | | |
| A1BG* | | | -0,815 | | | | | | | | | | | | | |
| GTF3C5* | | | -0,811 | | | | | | | | | | | | | |
| SMAD3* | | | -0,796 | | | | | | | | | | | | | |
| AC253572.2* | | | -0,794 | | | | | | | | | | | | | |
| NDEL1* | | | -0,788 | | | | | | | | | | | | | |
| ZNF737* | | | -0,776 | | | | | | | | | | | | | |
| TUBD1* | | | -0,773 | | | | | | | | | | | | | |
| OSER1-DT* | | | -0,769 | | | | | | | | | | | | | |
| PIGT* | | | -0,697 | | | | | | | | | | | | | |
| DERL2* | | | -0,696 | | | | | | | | | | | | | |
| SPAG9* | | | -0,687 | | | | | | | | | | | | | |
| AC007952.4* | | | -0,681 | | | | | | | | | | | | | |
| DNAJB2* | | | -0,656 | | | | | | | | | | | | | |
| BCL7B* | | | -0,651 | | | | | | | | | | | | | |
| KPNA3* | | | -0,624 | | | | | | | | | | | | | |

| GENES | **Memory B** | **Naive B** | **Naive CD4** | **Naive CD8** | **MAIT** | **Th17** | **Th1** | **Treg** | Th2 | Tfh | NK | NKT | CD8 CM | CD8 EM | CD8 EMRA + | gdTcells |
|---|---|---|---|---|---|---|---|---|---|---|---|---|---|---|---|---|
| NAA35* | | | -0,621 | | | | | | | | | | | | | |
| PPP1R18* | | | -0,614 | | | | | | | | | | | | | |
| GTF2F2* | | | -0,601 | | | | | | | | | | | | | |
| FAM76B* | | | -0,552 | | | | | | | | | | | | | |
| SETD5* | | | -0,520 | | | | | | | | | | | | | |
| LZIC* | | | -0,516 | | | | | | | | | | | | | |
| CLDND1* | | | -0,507 | | | | | | | | | | | | | |
| C1orf162* | | | -0,494 | | | | | | | | | | | | | |
| SF3A3* | | | -0,486 | | | | | | | | | | | | | |
| DNM1L* | | | -0,476 | | | | | | | | | | | | | |
| SMARCB1* | | | -0,475 | | | | | | | | | | | | | |
| ACAA1* | | | -0,471 | | | | | | | | | | | | | |
| NSMCE3* | | | -0,468 | | | | | | | | | | | | | |
| THAP12* | | | -0,431 | | | | | | | | | | | | | |
| MRPL34* | | | -0,406 | | | | | | | | | | | | | |
| SF3B5* | | | -0,402 | | | | | | | | | | | | | |
| HSPA5* | | | -0,354 | | | | | | | | | | | | | |
| UFC1* | | | -0,352 | | | | | | | | | | | | | |
| TRAPPC1* | | | -0,327 | | | | | | | | | | | | | |
| OGT* | | | 0,330 | | | | | | | | | | | | | |
| ITK* | | | 0,363 | | | | | | | | | | | | | |
| DOCK10* | | | 0,365 | | | | | | | | | | | | | |
| FRG1* | | | 0,384 | | | | | | | | | | | | | |
| ESYT2* | | | 0,396 | | | | | | | | | | | | | |
| TMEM161B-AS1* | | 0,429 | | | | | | | | | | | | | | |
| DDHD1* | | | 0,438 | | | | | | | | | | | | | |
| TOP2B* | | | 0,441 | | | | | | | | | | | | | |
| CREBZF* | | | 0,444 | | | | | | | | | | | | | |
| NMT2* | | | 0,447 | | | | | | | | | | | | | |
| CITED2* | | | 0,462 | | | | | | | | | | | | | |
| U2AF1L4* | | | 0,480 | | | | | | | | | | | | | |
| AP2B1* | | | 0,483 | | | | | | | | | | | | | |
| PPP6R3* | | | 0,484 | | | | | | | | | | | | | |
| SETD2* | | | 0,516 | | | | | | | | | | | | | |
| LPXN* | | | 0,566 | | | | | | | | | | | | | |

| GENES | **Memory B** | **Naive B** | **Naive CD4** | **Naive CD8** | **MAIT** | **Th17** | **Th1** | **Treg** | Th2 | Tfh | NK | NKT | CD8 CM | CD8 EM | CD8 EMRA+ | gdTcells |
|---|---|---|---|---|---|---|---|---|---|---|---|---|---|---|---|---|
| CERT1* | | | 0,588 | | | | | | | | | | | | | |
| SNHG25* | | | 0,608 | 0,489 | | | | | | | | | | | | |
| CEP68* | | | 0,636 | | | | | | | | | | | | | |
| MCFD2* | | | 0,651 | | | | | | | | | | | | | |
| GIGYF1* | | | 0,652 | | | | | | | | | | | | | |
| SLC25A42* | | | 0,653 | | | | | | | | | | | | | |
| MATR3.1* | | | 0,656 | | | | | | | | | | | | | |
| EDEM3* | | | 0,656 | | | -0,821 | | | | | | | | | | |
| GPR183* | | | 0,660 | | | | | | | | | | | | | |
| IFRD2* | | | 0,660 | | | | | | | | | | | | | |
| TMX3* | | | 0,669 | | | | | | | | | | | | | |
| HNRNPA1P48* | | | 0,710 | | | | | | | | | | | | | |
| AC090948.1* | | | 0,776 | | | | | | | | | | | | | |
| TRA2B/* | | | 0,788 | | | -0,508* | | | | | | | | | | |
| TAGLN2 | | | 0,803 | | | | | | | 0,826 | | | | 0,753 | | |
| FAM226B* | | | 0,815 | | | | | | | | | | | | | |
| WDR20* | | | 0,815 | | | | | | | | | | | | | |
| C12orf57 | | | 0,815 | | | | | | | | | | | | | |
| MKNK1* | | | 0,832 | | | | | | | | | | | | | |
| CERS6* | | | 0,835 | | | | | | | | | | | | | |
| TCIRG1* | | | 0,837 | | | | | | | | | | | | | |
| DNAJC5* | | | 0,840 | | | | | | | | | | | | | |
| NME6* | | | 0,841 | | | | | | | | | | | | | |
| REX1BD | | | 0,846 | | | | | | | | | | | | | |
| KMT2A | | | 0,851 | | | | | | | | | | | | | |
| **C1QBP** | | | 0,853 | | | | | | | | | | | | | |
| ARHGEF7* | | | 0,855 | | | | | | | | | | | | | |
| DDX5 | | | 0,856 | | | | | | | | | | | | | |
| TPR | | | 0,858 | | | | | | | | | | | | | |
| DGKA | | | 0,865 | | | | | | | | | | | | | |
| KAT2B* | | | 0,870 | | | | | | | | | | | | | |
| KHDRBS1 | | | 0,875 | | | | | | | | | | | | | |
| **DUSP2/*** | | | 0,880 | | | | 1,924/0,858* | | | | | | | | | |

| GENES | **Memory B** | **Naive B** | **Naive CD4** | **Naive CD8** | **MAIT** | **Th17** | **Th1** | **Treg** | Th2 | Tfh | NK | NKT | CD8 CM | CD8 EM | CD8 EMRA+ | gdTcells |
|---|---|---|---|---|---|---|---|---|---|---|---|---|---|---|---|---|
| LAMTOR4 | | | 0,884 | 1,230 | | | | | | | | 1,410 | | | | |
| LMBR1* | | | 0,884 | | | | | | | | | | | | | |
| HMGN1 | | | 0,893 | | | | | | | | | | | | | |
| **NDUFA1/*** | | | 0,897 | | 1,776/0,531* | | | | | | | | | | | |
| ARPC3 | | | 0,900 | | | | | | | | | | | | | |
| COMMD6 | | | 0,916 | | | | | | 1,263 | | | | | | | |
| RPLP1 | | | 0,917 | | | | | | | | | | | | | |
| SLC25A23* | | | 0,999 | | | | | | | | | | | | | |
| AC003102.1* | | | 1,015 | | | | | | | | | | | | | |
| TGFBR1* | | | 1,030 | | | | | | | | | | | | | |
| WASL* | | | 1,038 | | | | | | | | | | | | | |
| TRIM11* | | | 1,045 | | | | | | | | | | | | | |
| ZNF66* | | | 1,054 | | | | | | | | | | | | | |
| NEDD9* | | | 1,062 | | | | | | | | | | | | | |
| **IKZF1** | | | 1,115 | 1,243 | 0,824 | | | | | | | | | | | |
| PALM2-AKAP2* | | 1,123 | | | | | | | | | | | | | | |
| LRRC7* | | | 1,157 | | | | | | | | | | | | | |
| AL161457.2* | | | 1,170 | | | | | | | | | | | | | |
| AC068888.2* | | | 1,294 | | | | | | | | | | | | | |
| GTDC1* | | | 1,296 | | | | | | | | | | | | | |
| MSANTD3* | | | 1,393 | | | | | | | | | | | | | |
| ZHX3* | | | 1,430 | | | | | | | | | | | | | |
| CST7* | | | 1,464 | | | | | | | | | | | | | |
| HEXD-IT1* | | | 1,482 | | | | | | | | | | | | | |
| AC093227.1* | | | 1,487 | | | | | | | | | | | | | |
| **PIK3IP1-AS1*** | | | 1,505 | | | | | | | | | | | | | |
| AMT* | | | 1,566 | | | | | | | | | | | | | |
| ABCC4* | | | 1,641 | | | | | | | | | | | | | |
| NRROS* | | | 1,654 | | | | | | | | | | | | | |
| FAM174A* | | | 1,671 | | | | | | | | | | | | | |
| ADCY3* | | | 1,848 | | | | | | | | | | | | | |
| AC105429.1* | | | 1,853 | | | | | | | | | | | | | |
| TRIM46* | | | 1,921 | | | | | | | | | | | | | |
| AL162390.1* | | | 2,134 | | | | | | | | | | | | | |

| GENES | **Memory B** | **Naive B** | **Naive CD4** | **Naive CD8** | **MAIT** | **Th17** | **Th1** | **Treg** | Th2 | Tfh | NK | NKT | CD8 CM | CD8 EM | CD8 EMRA + | gdTcells |
|---|---|---|---|---|---|---|---|---|---|---|---|---|---|---|---|---|
| **IFI44L*** | | | 2,273 | 2,044 | | | | | | | | | | | | |
| KLHL17* | | | 2,278 | | | | | | | | | | | | | |
| AL358781.1* | | | 2,417 | | | | | | | | | | | | | |
| AL049780.1* | | | 2,430 | | | | | | | | | | | | | |
| AL356488.2* | | | 2,492 | | | | | | | | | | | | | |
| HOXB3* | | | 2,496 | | | | | | | | | | | | | |
| RTP4* | | | 2,514 | | | | | | | | | | | | | |
| KIZ-AS1* | | | 2,850 | | | | | | | | | | | | | |
| U73169.1* | | | 2,865 | | | | | | | | | | | | | |
| DISP2* | | | 3,375 | | | | | | | | | | | | | |
| MSH5* | | | 3,437 | | | | | | | | | | | | | |
| INCENP* | | | 3,575 | | | | | | | | | | | | | |
| RCAN3/* | | | 0,230* | | | 1,224 | | | | | | | | | | |
| SEMA4D/* | | | 0,378* | | | | | | 1,614 | | | | | | | |
| **CYTIP/*** | | | 0,407* | | | 1,527/0,520* | | | | | | | | | | |
| **KLF6/*** | | | -0,743* | | | | 0,652/-0,631* | | | | | | | | | |
| GPRIN3/* | | | 0,779* | | | | | | 1,588 | | | | | | | |
| **CDC42SE2/*** | | | 1,168/0,255 | | | | | | | | | | | | | |
| **UCKL1-AS1*** | | | | -3,137 | | | | | | | | | | | | |
| FANCA* | | | | -2,892 | | | | | | | | | | | | |
| MED18* | | | | -2,841 | | | | | | | | | | | | |
| C9orf40* | | | | -2,779 | | | | | | | | | | | | |
| ZNF619* | | | | -2,740 | | | | | | | | | | | | |
| SERPINI1* | | | | -2,477 | | | | | | | | | | | | |
| SLC35C1* | | | | -2,309 | | | | | | | | | | | | |
| LMTK3* | | | | -2,046 | | | | | | | | | | | | |
| DOCK7* | | | | -1,950 | | | | | | | | | | | | |
| STARD3NL* | | | | -1,728 | | | | | | | | | | | | |
| TRIM39* | | | | -1,710 | | | | | | | | | | | | |
| UBE2S* | | | | -1,607 | | | | | | | | | | | | |

| GENES | **Memory B** | **Naive B** | **Naive CD4** | **Naive CD8** | **MAIT** | **Th17** | **Th1** | **Treg** | Th2 | Tfh | NK | NKT | CD8 CM | CD8 EM | CD8 EMRA+ | gdTcells |
|---|---|---|---|---|---|---|---|---|---|---|---|---|---|---|---|---|
| CASTOR1* | | | | -1,478 | | | | | | | | | | | | |
| AP3S2* | | | | -1,467 | | | | | | | | | | | | |
| AC124016.1* | | | | -1,456 | | | | | | | | | | | | |
| ZNF561* | | | | -1,443 | -2,337 | | | | | | | | | | | |
| GALNT4* | | | | -1,400 | | | | | | | | | | | | |
| SNAPC3* | | | | -1,388 | | | | | | | | | | | | |
| TAB1* | | | | -1,380 | | | | | | | | | | | | |
| TTC16* | | | | -1,324 | -1,702 | | | | | | | | | | | |
| AC005224.2* | | | | -1,173 | | | | | | | | | | | | |
| C15orf61* | | | | -1,064 | | | | | | | | | | | | |
| STRN* | | | | -1,000 | | | | | | | | | | | | |
| ELMSAN1* | | | | -0,946 | | | | | | | | | | | | |
| RNF5* | | | | -0,934 | | | | | | | | | | | | |
| PLPBP* | | | | -0,896 | | | | | | | | | | | | |
| OXR1* | | | | -0,861 | | | | | | | | | | | | |
| C4orf48* | | | | -0,776 | | | | | | | | | | | | |
| KMT2B* | | | | -0,773 | | | | | | | | | | | | |
| TSPYL2* | | | | -0,743 | | | | | | | | | | | | |
| HIKESHI* | | | | -0,740 | | | | | | | | | | | | |
| CNST* | | | | -0,729 | | | | | | | | | | | | |
| RNF187* | | | | -0,719 | | | | | | | | | | | | |
| NAE1* | | | | -0,608 | | | | | | | | | | | | |
| DHPS* | | | | -0,561 | | | | | | | | | | | | |
| SARS* | | | | -0,558 | | | | | | | | | | | | |
| FCMR* | | | | 0,390 | | | | | | | | | | | | |
| PAN3* | | | | 0,562 | | | | | | | | | | | | |
| TTC39C* | | | | 0,574 | | | | | | | | | | | | |
| CDKN2AIP* | | | | 0,631 | | | | | | | | | | | | |
| TBRG1* | | | | 0,707 | | | | | | | | | | | | |
| MIS18BP1* | | | | 0,711 | | | | | | | | | | | | |
| FAM117B* | | | | 0,733 | | | | | | | | | | | | |
| KPNA6* | | | | 0,755 | | | | | | | | | | | | |
| SLC5A3* | | | | 0,797 | | | | | | | | | | | | |
| FBXW4* | | | | 0,806 | | | | | | | | | | | | |
| RALGAPB* | | | | 0,868 | | | | | | | | | | | | |

| GENES | **Memory B** | **Naive B** | **Naive CD4** | **Naive CD8** | **MAIT** | **Th17** | **Th1** | **Treg** | Th2 | Tfh | NK | NKT | CD8 CM | CD8 EM | CD8 EMRA+ | gdTcells |
|---|---|---|---|---|---|---|---|---|---|---|---|---|---|---|---|---|
| LIPA* | | | | 0,871 | | | | | | | | | | | | |
| IFI6* | | | | 0,889 | | | | | | | | | | | | |
| CAMK1D* | | | | 0,917 | | | | | | | | | | | | |
| PDE4D* | | | | 0,925 | | | | | | | | | | | | |
| CCL5* | | | | 1,051 | | | | | | | | | | | | |
| LRIF1* | | | | 1,066 | | | | | | | | | | | | |
| GPBP1 | | | | 1,100 | | | | | | | 1,157 | | | | | |
| AC104532.2* | | | | 1,128 | | | | | | | | | | | | |
| FBL | | | | 1,155 | | | | | | | | | | | | |
| **YPEL3** | | | | 1,173 | | | | 1,471 | | 0,842 | | | | | | |
| **RICTOR** | | | | 1,208 | | | 0,744 | | | | | | | | | |
| MIGA2* | | | | 1,214 | | | | | | | | | | | | |
| ICAM3 | | | | 1,224 | | | | | | | | | | | | |
| DPP4* | | | | 1,241 | | | | | | | | | | | | |
| CMPK1 | | | | 1,250 | | | | | 1,340 | | | | | | | |
| RELL1* | | | | 1,258 | | | | | | | | | | | | |
| EID1 | | | | 1,272 | | 1,246 | | | | | | | | | | |
| PPP2R5C | | | | 1,281 | | | | | | | | | | | | |
| **CD53/*** | | | | 1,298 | | | | 0,896 | | | | | | | | |
| MZT2A | | | | 1,323 | | | | | | | | | | | | |
| ROCK2* | | | | 1,334 | | | | | | | | | | | | |
| TRAPPC9* | | | | 1,382 | | | | | | | | | | | | |
| **TLN1** | | | | 1,412 | | | | | | | | | | | | |
| METTL6* | | | | 1,474 | | | | | | | | | | | | |
| TMEM68* | | | | 1,603 | -1,910 | | | | | | | | | | | |
| DSE* | | | | 1,809 | | | | | | | | | | | | |
| TRMT5* | | | | 2,367 | | | | | | | | | | | | |
| CDKN2C* | | | | 2,477 | | | | | | | | | | | | |
| SLC39A11* | | | | 2,494 | | | | | | | | | | | | |
| RAPGEF2* | | | | 2,512 | | | | | | | | | | | | |
| LINC02019* | | | | 2,875 | | | | | | | | | | | | |
| **SP100/*** | | | | 1,299/0,418* | | | | | | | | | | | | |
| **RNF213/*** | | | | 1,307/0,377* | | | 0,478* | | | | | | | 1,420 | | |

| GENES | Memory B | Naive B | Naive CD4 | Naive CD8 | MAIT | Th17 | Th1 | Treg | Th2 | Tfh | NK | NKT | CD8 CM | CD8 EM | CD8 EMRA+ | gdTcells |
|---|---|---|---|---|---|---|---|---|---|---|---|---|---|---|---|---|
| **PABPC4/*** | | | | 1,316/0,450* | | | | | | | | | | | | |
| TTC27* | | | | | -3,764 | | | | | | | | | | | |
| CARF* | | | | | -3,469 | | | | | | | | | | | |
| SUMF1* | | | | | -3,220 | | | | | | | | | | | |
| DIABLO* | | | | | -2,867 | | | | | | | | | | | |
| LMF1* | | | | | -2,862 | | | | | | | | | | | |
| ATP6V1A* | | | | | -2,823 | | | | | | | | | | | |
| RGS3* | | | | | -2,776 | | | | | | | | | | | |
| NOD1* | | | | | -2,770 | | | | | | | | | | | |
| ZADH2* | | | | | -2,743 | | | | | | | | | | | |
| GMDS-DT* | | | | | -2,462 | | | | | | | | | | | |
| ICA1L* | | | | | -2,338 | | | | | | | | | | | |
| ITGA5* | | | | | -2,325 | | | | | | | | | | | |
| PUS10* | | | | | -2,301 | | | | | | | | | | | |
| LINC00641* | | | | | -2,299 | | | | | | | | | | | |
| DUSP14* | | | | | -2,289 | | | | | | | | | | | |
| ZNF566* | | | | | -2,186 | | | | | | | | | | | |
| ATP7A* | | | | | -2,152 | | | | | | | | | | | |
| MBOAT7* | | | | | -2,099 | | | | | | | | | | | |
| EID3* | | | | | -2,096 | | | | | | | | | | | |
| MIR22HG* | | | | | -1,977 | | | | | | | | | | | |
| NHSL2* | | | | | -1,950 | | | | | | | | | | | |
| C11orf54* | | | | | -1,937 | | | | | | | | | | | |
| SDHAF1* | | | | | -1,935 | | | | | | | | | | | |
| LRMP* | | | | | -1,837 | | | | | | | | | | | |
| PEX6* | | | | | -1,797 | | | | | | | | | | | |
| MAPK8IP3* | | | | | -1,794 | | | | | | | | | | | |
| RPUSD2* | | | | | -1,790 | | | | | | | | | | | |
| HDAC5* | | | | | -1,714 | | | | | | | | | | | |
| LINC00685* | | | | | -1,597 | | | | | | | | | | | |
| WDR37* | | | | | -1,587 | | | | | | | | | | | |
| AC048341.1* | | | | | -1,557 | | | | | | | | | | | |
| KAT8* | | | | | -1,525 | | | | | | | | | | | |
| SPRY1* | | | | | -1,494 | | | | | | | | | | | |

| GENES | Memory B | Naive B | Naive CD4 | Naive CD8 | MAIT | Th17 | Th1 | Treg | Th2 | Tfh | NK | NKT | CD8 CM | CD8 EM | CD8 EMRA + | gdTcells |
|---|---|---|---|---|---|---|---|---|---|---|---|---|---|---|---|---|
| ATE1* | | | | | -1,441 | | | | | | | | | | | |
| TCF12* | | | | | -1,440 | | | | | | | | | | | |
| KLHL28* | | | | | -1,417 | | | | | | | | | | | |
| RING1* | | | | | -1,373 | | | | | | | | | | | |
| SLC33A1* | | | | | -1,347 | | | | | | | | | | | |
| TARSL2* | | | | | -1,325 | | | | | | | | | | | |
| KIF21A* | | | | | -1,324 | | | | | | | | | | | |
| MRPL18* | | | | | -1,245 | | | | | | | | | | | |
| ZFYVE28* | | | | | -1,217 | | | | | | | | | | | |
| TM2D1* | | | | | -1,211 | | | | | | | | | | | |
| GOPC* | | | | | -1,203 | | | | | | | | | | | |
| SNX14* | | | | | -1,169 | | | | | | | | | | | |
| CEP290* | | | | | -1,153 | | | | | | | | | | | |
| PIEZO1* | | | | | -1,133 | | | | | | | | | | | |
| VCPIP1* | | | | | -1,100 | | | | | | | | | | | |
| BAG5* | | | | | -1,086 | | | | | | | | | | | |
| GAK* | | | | | -1,059 | | | | | | | | | | | |
| TMEM147* | | | | | -1,045 | | | | | | | | | | | |
| SSBP3* | | | | | -1,042 | | | | | | | | | | | |
| POU2F2* | | | | | -1,041 | | | | | | | | | | | |
| SLC38A2* | | | | | -0,986 | | | | | | | | | | | |
| MIDN* | | | | | -0,836 | | | | | | | | | | | |
| EIF2S2* | | | | | -0,801 | | | | | | | | | | | |
| CD27* | | | | | -0,796 | | | | | | | | | | | |
| EIF4A1* | | | | | 0,426 | | | | | | | | | | | |
| LCP1* | | | | | 0,435 | | | | | | | | | | | |
| LDHA* | | | | | 0,680 | | | | | | | | | | | |
| ITGB7* | | | | | 0,791 | | | | | | | | | | | |
| SRP19* | | | | | 0,810 | | | | | | | | | | | |
| DNAJC1* | | | | | 0,814 | | | | | | | | | | | |
| TXN* | | | | | 0,815 | | | | | | | | | | | |
| BUD31* | | | | | 0,842 | | | | | | | | | | | |
| RNF216* | | | | | 0,935 | | | | | | | | | | | |
| SECISBP2* | | | | | 1,074 | | | | | | | | | | | |
| TFAM* | | | | | 1,106 | | | | | | | | | | | |

| GENES | **Memory B** | **Naive B** | **Naive CD4** | **Naive CD8** | **MAIT** | **Th17** | **Th1** | **Treg** | Th2 | Tfh | NK | NKT | CD8 CM | CD8 EM | CD8 EMRA + | gdTcells |
|---|---|---|---|---|---|---|---|---|---|---|---|---|---|---|---|---|
| AP3B1* | | | | | 1,140 | | | | | | | | | | | |
| CAPRIN1* | | | | | 1,142 | | | | | | | | | | | |
| ZNF331* | | | | | 1,154 | | | | | | | | | | | |
| TMEM258 | | | | | 1,172 | | | | | | | | | | | |
| GNAS | | | | | 1,183 | | | | | | | | | | | |
| CD52 | | | | | 1,210 | | | | | | | | | | | 1,418 |
| **KIF2A/*** | | | | | 1,221 | | 0,672/-0,487* | | | | | | | | | |
| PPA2* | | | | | 1,228 | | | | | | | | | | | |
| PLEKHB2* | | | | | 1,243 | | | | | | | | | | | |
| RPS17 | | | | | 1,246 | | | | | 0,834 | | | | | | |
| ATP8A1* | | | | | 1,246 | | | | | | | | | | | |
| PHACTR2 | | | | | 1,249 | | | | | | | | | | | |
| UPF3B* | | | | | 1,263 | | | | | | | | | | | |
| POLR2L | | | | | 1,265 | | | | | | | | | | | |
| SUPT16H* | | | | | 1,280 | | | | | | | | | | | |
| GMFG | | | | | 1,282 | | | | | | | | | | | 0,777 |
| VIM | | | | | 1,289 | | | | | | | | | | | |
| ANKRD13A* | | | | | 1,294 | | | | | | | | | | | |
| HLA.E | | | | | 1,311 | 1,232 | | | | | | | | | | |
| GPSM3 | | | | | 1,316 | | | | | | | | | | | |
| RNF220* | | | | | 1,320 | | | | | | | | | | | |
| ANKRD28* | | | | | 1,343 | | | | | | | | | | | |
| GZMM | | | | | 1,362 | | | | | | | | | | | |
| ATRX | | | | | 1,364 | | | | | | | | | | | |
| ARPC5L | | | | | 1,367 | | | | | | | | | | | |
| **MTDH** | | | | | 1,375 | 1,311 | | | | | | | | | | |
| CAPZB | | | | | 1,378 | | | | | | | 1,803 | | | | |
| ITFG2* | | | | | 1,380 | | | 1,579 | | | | | | | | |
| SLC38A1 | | | | | 1,395 | | | | | | | | | | | |
| COX8A | | | | | 1,396 | | | | | | | | | | | |
| **TGOLN2** | | | | | 1,398 | | | | | | | | | | | |
| TMED2 | | | | | 1,412 | | | | | | | | | | | |
| TRAM1 | | | | | 1,447 | | | | | | | | 1,458 | | | |

| GENES | **Memory B** | **Naive B** | **Naive CD4** | **Naive CD8** | **MAIT** | **Th17** | **Th1** | **Treg** | Th2 | Tfh | NK | NKT | CD8 CM | CD8 EM | CD8 EMRA + | gdTcells |
|---|---|---|---|---|---|---|---|---|---|---|---|---|---|---|---|---|
| STK17B | | | | | 1,447 | | | | | | | | | | | |
| MGAT4A | | | | | 1,449 | | | | | | | | | | | |
| COPS4* | | | | | 1,466 | | | | | | | | | | | |
| SUMO1 | | | | | 1,475 | | | | | | | | | | | |
| DDX39B | | | | | 1,479 | | | | | | | | | | | |
| CCNL2* | | | | | 1,480 | | | | | | | | | | | |
| DAZAP2 | | | | | 1,486 | | | | | | | | | 0,867 | | |
| **SEC11A** | | | | | 1,490 | | | | | | | | | | | |
| PDIA3 | | | | | 1,506 | 1,204 | | | | | | | | | | |
| **PAIP2** | | | | | 1,527 | | | | | | | 1,461 | | | 1,520 | |
| C4orf3 | | | | | 1,546 | | | | | | | | | | | |
| RBMX | | | | | 1,558 | | | | | | | | | | | |
| **SRI** | | | | | 1,620 | | | | | | | | | | | |
| **ISG15/*** | | | | | 1,627 | | | 1,017 | | | | | | | | |
| H2AFZ | | | | | 1,649 | | | | | | | | | | | |
| **SNRPN** | | | | | 1,681 | | | 0,746 | | | | | | | | |
| **STK17A** | | | | | 1,686 | | | | | | | | | | | |
| STX10* | | | | | 1,695 | | | | | | | | | | | |
| ULK3* | | | | | 1,718 | | | | | | | | | | | |
| HPF1* | | | | | 1,720 | | | | | | | | | | | |
| **MAF** | | | | | 1,726 | | | | | | | | | | | |
| ENO1 | | | | | 1,727 | 1,205 | | | | | | | | | | |
| HADH* | | | | | 1,796 | | | | | | | | | | | |
| **DIAPH1** | | | | | 1,797 | | | | | | | | | | | |
| COA4* | | | | | 1,803 | | | | | | | | | | | |
| P2RX5* | | | | | 1,821 | | | | | | | | | | | |
| HMG20B* | | | | | 1,897 | | | | | | | | | | | |
| ACAT2* | | | | | 1,901 | | | | | | | | | | | |
| NME4* | | | | | 1,912 | | | | | | | | | | | |
| **TAOK3** | | | | | 1,916 | | | | 1,692 | | | | | | | |
| TCF20* | | | | | 1,929 | | | | | | | | | | | |
| WDR18* | | | | | 2,035 | | | | | | | | | | | |
| SFXN3* | | | | | 2,039 | | | | | | | | | | | |
| **PRPF40A** | | | | | 2,046 | | | | | | | | | | | |
| TRIQK* | | | | | 2,098 | | | | | | | | | | | |

| GENES | **Memory B** | **Naive B** | **Naive CD4** | **Naive CD8** | **MAIT** | **Th17** | **Th1** | **Treg** | Th2 | Tfh | NK | NKT | CD8 CM | CD8 EM | CD8 EMRA + | gdTcells |
|---|---|---|---|---|---|---|---|---|---|---|---|---|---|---|---|---|
| PET117* | | | | | 2,149 | | | | | | | | | | | |
| GUF1* | | | | | 2,180 | | | | | | | | | | | |
| NOP14-AS1* | | | | | 2,363 | | | | | | | | | | | |
| PRPS2* | | | | | 2,374 | | | | | | | | | | | |
| GALNT2* | | | | | 2,444 | | | | | | | | | | | |
| PRCP* | | | | | 2,448 | | | | | | | | | | | |
| HIST1H2AC* | | | | | 2,486 | | | | | | | | | | | |
| TMEM185B* | | | | | 2,556 | | | | | | | | | | | |
| NUP85* | | | | | 2,613 | | | | | | | | | | | |
| ARHGAP21* | | | | | 2,620 | | | | | | | | | | | |
| UQCC1* | | | | | 2,707 | | | | | | | | | | | |
| TMEM159* | | | | | 2,870 | | | | | | | | | | | |
| ATP6V1E2* | | | | | 2,892 | | | | | | | | | | | |
| NKIRAS2* | | | | | 2,978 | | | | | | | | | | | |
| UBE2D4* | | | | | 2,986 | | | | | | | | | | | |
| URGCP* | | | | | 3,200 | | | | | | | | | | | |
| INTS13* | | | | | 3,230 | | | | | | | | | | | |
| SNN* | | | | | 3,357 | | | | | | | | | | | |
| USP19* | | | | | 3,382 | | | | | | | | | | | |
| BRI3BP* | | | | | 3,430 | | | | | | | | | | | |
| HIST2H2BF* | | | | | 3,623 | | | | | | | | | | | |
| FBXL6* | | | | | 3,888 | | | | | | | | | | | |
| **UBXN1/*** | | | | | 0,699/-0,601* | | | | | | | | | | | |
| **NDUFA11/*** | | | | | 0,708/-0,593* | | | | 1,334 | | | | | | | |
| SAT1/* | | | | | 1,495/0,534* | | | | | | | | | | | |
| PABPC1/* | | | | | 1,588/0,539* | | 1,217 | | | | | 1,630 | | | | |
| **SUB1/*** | | | | | 1,663/0,426* | | | | | | | | | | | |
| **CADM1*** | | | | | | -3,261 | | | | | | | | | | |
| **E2F2*** | | | | | | -3,153 | | | | | | | | | | |
| DHX34* | | | | | | -2,781 | | | | | | | | | | |

| GENES | **Memory B** | **Naive B** | **Naive CD4** | **Naive CD8** | **MAIT** | **Th17** | **Th1** | **Treg** | Th2 | Tfh | NK | NKT | CD8 CM | CD8 EM | CD8 EMRA + | gdTcells |
|---|---|---|---|---|---|---|---|---|---|---|---|---|---|---|---|---|
| SLC25A14* | | | | | | -2,651 | | | | | | | | | | |
| TRDC* | | | | | | -2,506 | -3,112 | | | | | | | | | |
| ZSCAN16* | | | | | | -2,330 | | | | | | | | | | |
| **NEDD4L*** | | | | | | -2,315 | | | | | | | | | | |
| ZSWIM9* | | | | | | -2,232 | | | | | | | | | | |
| RNF121* | | | | | | -2,170 | | | | | | | | | | |
| TCEAL1* | | | | | | -1,762 | | | | | | | | | | |
| KIAA0930* | | | | | | -1,726 | | | | | | | | | | |
| MED31* | | | | | | -1,481 | | | | | | | | | | |
| INTS3* | | | | | | -1,284 | | | | | | | | | | |
| WDR36* | | | | | | -1,260 | | | | | | | | | | |
| ITGAE* | | | | | | -1,160 | | | | | | | | | | |
| GTF2H3* | | | | | | -1,065 | | | | | | | | | | |
| ACLY* | | | | | | -1,033 | | | | | | | | | | |
| EXOC5* | | | | | | -1,015 | | | | | | | | | | |
| SUDS3* | | | | | | -1,007 | | | | | | | | | | |
| ATF1* | | | | | | -0,931 | | | | | | | | | | |
| PTRHD1* | | | | | | -0,875 | | | | | | | | | | |
| KIAA0586* | | | | | | -0,859 | | | | | | | | | | |
| PTTG1IP* | | | | | | -0,822 | | | | | | | | | | |
| WDFY2* | | | | | | -0,820 | | | | | | | | | | |
| MYADM* | | | | | | -0,748 | -1,136 | | | | | | | | | |
| HPS1* | | | | | | -0,743 | | -1,560 | | | | | | | | |
| ICE2* | | | | | | -0,743 | | | | | | | | | | |
| OGDH* | | | | | | -0,732 | | | | | | | | | | |
| WDR43* | | | | | | -0,676 | | | | | | | | | | |
| UCP2* | | | | | | -0,621 | | | | | | | | | | |
| TMEM256* | | | | | | -0,607 | | | | | | | | | | |
| CCNH* | | | | | | -0,580 | | | | | | | | | | |
| MYO1G* | | | | | | -0,494 | | | | | | | | | | |
| NOSIP* | | | | | | -0,392 | | | | | | | | | | |
| ITGB1* | | | | | | -0,363 | | | | | | | | | | |
| MT-ND1* | | | | | | 0,205 | | | | | | | | | | |
| MT-ATP6* | | | | | | 0,222 | | | | | | | | | | |
| MT-ND2* | | | | | | 0,229 | | | | | | | | | | |

| GENES | **Memory B** | **Naive B** | **Naive CD4** | **Naive CD8** | **MAIT** | **Th17** | **Th1** | **Treg** | Th2 | Tfh | NK | NKT | CD8 CM | CD8 EM | CD8 EMRA + | gdTcells |
|---|---|---|---|---|---|---|---|---|---|---|---|---|---|---|---|---|
| RPL38* | | | | | | 0,260 | | | | | | | | | | |
| HLA-E* | | | | | | 0,262 | | | | | | | | | | |
| RBMS1* | | | | | | 0,450 | | | | | | | | | | |
| TNFRSF14* | | | | | | 0,487 | | | | | | | | | | |
| SNRNP70* | | | | | | 0,534 | | | | | | | | | | |
| N4BP2L1* | | | | | | 0,656 | | | | | | | | | | |
| GATAD2A* | | | | | | 0,662 | | | | | | | | | | |
| **MALT1** | | | | | | 0,729 | | | | | | | | | | |
| PPM1G | | | | | | 0,752 | | | | | | | | | | |
| NELL2* | | | | | | 0,756 | | | | | | | | | | |
| **PRDX5** | | | | | | 0,757 | | | | | | | | | | |
| TAP2* | | | | | | 0,766 | | | | | | | | | | |
| APOL3* | | | | | | 0,766 | | | | | | | | | | |
| FNIP1* | | | | | | 0,767 | | | | | | | | | | |
| SYF2 | | | | | | 0,801 | | | | | | | | | | |
| TRPM7* | | | | | | 0,811 | | | | | | | | | | |
| CRIP1 | | | | | | 0,852 | | | 1,457 | | | 0,792 | | | | |
| GBP5* | | | | | | 0,884 | | | | | | | | | | |
| PRR12* | | | | | | 0,931 | | | | | | | | | | |
| MAPKAPK3* | | | | | | 0,987 | | | | | | | | | | |
| AP2A1* | | | | | | 1,005 | | | | | | | | | | |
| AASDH* | | | | | | 1,035 | | | | | | | | | | |
| FAU | | | | | | 1,041 | | | | | 1,105 | | | | | |
| RPS13 | | | | | | 1,048 | | | | | | | | | | |
| RPL32 | | | | | | 1,064 | | | | | | | | | | |
| RPS19 | | | | | | 1,068 | | | | | | | | | | |
| RPS9 | | | | | | 1,068 | | | | | | | | | | |
| CSDE1 | | | | | | 1,069 | | | | | | 1,441 | | | | |
| RPS23 | | | | | | 1,070 | | | | | | | | | | |
| RPL12 | | | | | | 1,071 | | | | | | | | | | |
| SH2D3A* | | | | | | 1,071 | | | | | | | | | | |
| PHYH* | | | | | | 1,072 | | | | | | | | | | |
| RPL30 | | | | | | 1,072 | | | | | | | | | | |
| EEF1A1/* | | | | | | 1,075 | | -0,184/* | | | | | | | | |
| NACA | | | | | | 1,077 | | | | | | | | | | |

| GENES | **Memory B** | **Naive B** | **Naive CD4** | **Naive CD8** | **MAIT** | **Th17** | **Th1** | **Treg** | Th2 | Tfh | NK | NKT | CD8 CM | CD8 EM | CD8 EMRA + | gdTcells |
|---|---|---|---|---|---|---|---|---|---|---|---|---|---|---|---|---|
| RPS12 | | | | | | 1,087 | | | | | | | | | | |
| RPL9 | | | | | | 1,088 | | | | | 1,103 | | | | | |
| RPL5 | | | | | | 1,089 | | | | | | | | | | |
| **EEF1B2/*** | | | | | | 1,091 | | 0,727/-0,401* | | | | | | | | |
| RPL17 | | | | | | 1,092 | | | | | | | | | | |
| RPS5 | | | | | | 1,093 | | | | | | | | | | |
| UBA52 | | | | | | 1,093 | | | | | | | | | | |
| TPT1 | | | | | | 1,093 | | | | | | | | | | |
| RPS6 | | | | | | 1,095 | | | | | | | | | | |
| RPL41 | | | | | | 1,097 | | | | | | | | | | |
| RPS15 | | | | | | 1,100 | | | | | | | | | | |
| RPL13A | | | | | | 1,100 | | | | | | | | | | |
| RPL19 | | | | | | 1,101 | | | | | | | | | | |
| RPL29 | | | | | | 1,101 | | | | | | | | | | |
| RPS25 | | | | | | 1,103 | | | | | | | | | | |
| RPL11 | | | | | | 1,103 | | | | | | | | | | |
| RPL13 | | | | | | 1,103 | | | | | | | | | | |
| TOMM7 | | | | | | 1,104 | | | | | | | | | | |
| RPLP2 | | | | | | 1,105 | | | | | | | | | | |
| RPS2 | | | | | | 1,106 | | | | | | | | | | |
| RPL35 | | | | | | 1,106 | | | | | | | | | | |
| RPS18 | | | | | | 1,108 | | | | | | | | | | |
| EEF1G | | | | | | 1,110 | | | | | | | | | | |
| RPL18A | | | | | | 1,114 | | | | | | 1,215 | | | | |
| EEF1D | | | | | | 1,116 | | | | | | | | | | |
| RPL24 | | | | | | 1,123 | | | | | | | | | | |
| XAF1* | | | | | | 1,125 | | | | | | | | | | |
| TRIR | | | | | | 1,135 | | | | | | | | | | |
| PTMA | | | | | | 1,137 | | | | | | | | | | |
| RPL27A | | | | | | 1,141 | | | | | | | | | | |
| RPS20 | | | | | | 1,145 | | | | | | | | | | |
| RPL31 | | | | | | 1,159 | | | | | | | | | | |
| YWHAB | | | | | | 1,160 | | | | | | | 1,615 | | | |
| GNAI2 | | | | | | 1,167 | | | | | | | | | | |

| GENES | **Memory B** | **Naive B** | **Naive CD4** | **Naive CD8** | **MAIT** | **Th17** | **Th1** | **Treg** | Th2 | Tfh | NK | NKT | CD8 CM | CD8 EM | CD8 EMRA+ | gdTcells |
|---|---|---|---|---|---|---|---|---|---|---|---|---|---|---|---|---|
| KTI12* | | | | | | 1,168 | | | | | | | | | | |
| SNX10* | | | | | | 1,184 | | | | | | | | | | |
| MT.ND1 | | | | | | 1,185 | | | | | | | | 1,118 | | |
| MT.ND2 | | | | | | 1,187 | | | | | | | | | | |
| MT.CYB | | | | | | 1,191 | | | | | | | | | | |
| CD48 | | | | | | 1,199 | | | | | | | | | | |
| SREBF2* | | | | | | 1,199 | | | | | | | | | | |
| MT.ATP6 | | | | | | 1,205 | | | | | | | | | | |
| ACTR3 | | | | | | 1,243 | | | | | | | | | | |
| PPIA | | | | | | 1,244 | | | | | | | | | | |
| FKBP11 | | | | | | 1,279 | | | | | | | | | | |
| GPS2 | | | | | | 1,282 | | | | | | | | | | |
| **PSMB9** | | | | | | 1,299 | | | | | | | | | | |
| **HNRNPM** | | | | | | 1,314 | | | | | | | | | | |
| **PRKDC** | | | | | | 1,337 | | | | | | | | | | |
| **RAP1A** | | | | | | 1,349 | | | | | | | | | | |
| LSM8 | | | | | | 1,409 | | | | | | | | | | |
| **SNHG14/*** | | | | | | 1,418 | 0,592/-0,676* | | | 0,611 | | | | | | |
| TMEM14B | | | | | | 1,477 | | | | | | | | | | |
| FMR1-IT1* | | | | | | 1,598 | | | | | | | | | | |
| SLC38A5* | | | | | | 1,648 | | | | | | | | | | |
| TRIM25* | | | | | | 1,651 | | | | | | | | | | |
| CIAO3* | | | | | | 1,695 | | | | | | | | | | |
| ZNF677* | | | | | | 1,743 | | | | | | | | | | |
| SNRNP25* | | | | | | 1,783 | | | | | | | | | | |
| **GZMK*** | | | | | | 1,879 | | | | | | | | | | |
| BRPF3* | | | | | | 1,941 | | | | | | | | | | |
| CABP4* | | | | | | 1,953 | | | | | | | | | | |
| ZNF337-AS1* | | | | | | 1,984 | | | | | | | | | | |
| MCOLN3* | | | | | | 2,110 | | | | | | | | | | |
| FAM229A* | | | | | | 2,138 | | | | | | | | | | |
| AC108134.2* | | | | | | 2,237 | | | | | | | | | | |
| DNAL1* | | | | | | 2,306 | | | | | | | | | | |

| GENES | **Memory B** | **Naive B** | **Naive CD4** | **Naive CD8** | **MAIT** | **Th17** | **Th1** | **Treg** | Th2 | Tfh | NK | NKT | CD8 CM | CD8 EM | CD8 EMRA+ | gdTcells |
|---|---|---|---|---|---|---|---|---|---|---|---|---|---|---|---|---|
| **MIER2*** | | | | | | 2,619 | | | | | | | | | | |
| **IFITM2/*** | | | | | | 1,317/0, 336* | | | | | | | | | | |
| **NUCKS1/*** | | | | | | 1,403/0, 394* | | | | | | | | | | |
| **NFE2L3*** | | | | | | | -2,870 | | | | | | | | | |
| NOTCH2NLA* | | | | | | | -2,800 | | | | | | | | | |
| C15orf41* | | | | | | | -2,682 | | | | | | | | | |
| ACVR1B* | | | | | | | -2,597 | | | | | | | | | |
| BLM* | | | | | | | -2,548 | | | | | | | | | |
| KCTD15* | | | | | | | -2,443 | | | | | | | | | |
| RFC3* | | | | | | | -2,432 | | | | | | | | | |
| **LAPTM4B*** | | | | | | | -2,425 | | | | | | | | | |
| HIST1H2BF* | | | | | | | -2,415 | | | | | | | | | |
| THAP3* | | | | | | | -2,033 | | | | | | | | | |
| SPATS2L* | | | | | | | -2,006 | | | | | | | | | |
| PAXIP1* | | | | | | | -1,852 | | | | | | | | | |
| ZNF699* | | | | | | | -1,823 | | | | | | | | | |
| INTS8* | | | | | | | -1,814 | | | | | | | | | |
| CD151* | | | | | | | -1,790 | | | | | | | | | |
| ZNF587B* | | | | | | | -1,638 | | | | | | | | | |
| ZNF346* | | | | | | | -1,609 | | | | | | | | | |
| UBE2D1* | | | | | | | -1,589 | | | | | | | | | |
| SPICE1* | | | | | | | -1,584 | | | | | | | | | |
| SNAPC2* | | | | | | | -1,540 | | | | | | | | | |
| SNX27* | | | | | | | -1,442 | | | | | | | | | |
| NTPCR* | | | | | | | -1,431 | | | | | | | | | |
| DLGAP1-AS1* | | | | | | | -1,377 | | | | | | | | | |
| CCDC65* | | | | | | | -1,181 | | | | | | | | | |
| LINC00824* | | | | | | | -1,180 | | | | | | | | | |
| UBN2* | | | | | | | -1,178 | | | | | | | | | |
| MBD1* | | | | | | | -1,166 | -1,905 | | | | | | | | |
| **THEMIS*** | | | | | | | -1,081 | -3,341 | | | | | | | | |

| GENES | **Memor y B** | **Naive B** | **Naive CD4** | **Naive CD8** | **MAIT** | **Th17** | **Th1** | **Treg** | Th2 | Tfh | NK | NKT | CD8 CM | CD8 EM | CD8 EMRA + | gdTcell s |
|---|---|---|---|---|---|---|---|---|---|---|---|---|---|---|---|---|
| AC103591.3* | | | | | | | -1,077 | | | | | | | | | |
| CNIH4* | | | | | | | -1,049 | | | | | | | | | |
| TRNAU1AP* | | | | | | | -1,033 | | | | | | | | | |
| RBM14* | | | | | | | -0,976 | | | | | | | | | |
| HMGXB4* | | | | | | | -0,936 | | | | | | | | | |
| AL021155.5* | | | | | | | -0,880 | | | | | | | | | |
| NDUFV3* | | | | | | | -0,868 | | | | | | | | | |
| AATF* | | | | | | | -0,810 | | | | | | | | | |
| ZNF33A* | | | | | | | -0,731 | | | | | | | | | |
| AL499604.1* | | | | | | | -0,721 | | | | | | | | | |
| SCML4* | | | | | | | -0,676 | | | | | | | | | |
| PUF60* | | | | | | | -0,661 | | | | | | | | | |
| NCOR1* | | | | | | | -0,629 | | | | | | | | | |
| LCP2* | | | | | | | -0,594 | | | | | | | | | |
| SMARCA5* | | | | | | | -0,562 | -0,852 | | | | | | | | |
| PSMA6* | | | | | | | 0,557 | | | | | | | | | |
| ATF6B* | | | | | | | 0,610 | | | | | | | | | |
| SUMO3* | | | | | | | 0,750 | | | | | | | | | |
| SF3B1 | | | | | | | 0,767 | | | | | | | | | |
| DNAJB9* | | | | | | | 0,777 | | | | | | | | | |
| SDHAF2* | | | | | | | 0,780 | | | | | | | | | |
| MED13* | | | | | | | 0,784 | | | | | | | | | |
| DR1* | | | | | | | 0,808 | | | | | | | | | |
| GABARAP | | | | | | | 0,810 | | | | | | | | | |
| PSME4* | | | | | | | 0,836 | | | | | | | | | |
| ZNHIT3* | | | | | | | 0,845 | | | | | | | | | |
| EBPL* | | | | | | | 0,873 | | | | | | | | | |
| TTC39B* | | | | | | | 0,935 | | | | | | | | | |
| TRIAP1* | | | | | | | 0,946 | | | | | | | | | |
| MFHAS1* | | | | | | | 0,948 | | | | | | | | | |
| TNFRSF4* | | | | | | | 1,042 | | | | | | | | | |
| CSKMT* | | | | | | | 1,051 | | | | | | | | | |
| TPST2* | | | | | | | 1,125 | | | | | | | | | |
| KLHL20* | | | | | | | 1,142 | | | | | | | | | |
| GABARAPL1* | | | | | | | 1,209 | | | | | | | | | |

| GENES | **Memory B** | **Naive B** | **Naive CD4** | **Naive CD8** | **MAIT** | **Th17** | **Th1** | **Treg** | Th2 | Tfh | NK | NKT | CD8 CM | CD8 EM | CD8 EMRA+ | gdTcells |
|---|---|---|---|---|---|---|---|---|---|---|---|---|---|---|---|---|
| CFH* | | | | | | | 1,219 | | | | | | | | | |
| IFI44* | | | | | | | 1,222 | | | | | | | | | |
| C19orf53 | | | | | | | 1,230 | | | | | | | | | |
| GUSB* | | | | | | | 1,255 | | | | | | | | | |
| RAN | | | | | | | 1,323 | | | | | | | | | 1,700 |
| RBM38* | | | | | | | 1,369 | | | | | | | | | |
| MGST3 | | | | | | | 1,384 | | 1,410 | | | | | | | |
| SSU72 | | | | | | | 1,443 | | | | | | | | | |
| URI1 | | | | | | | 1,467 | | | | | | | | | |
| NR2C2AP* | | | | | | | 1,493 | | | | | | | | | |
| HYPK* | | | | | | | 1,506 | | | | | | | | | |
| MAP3K14* | | | | | | | 1,536 | | | | | | | | | |
| MPP6* | | | | | | | 1,609 | | | | | | | | | |
| APOOL* | | | | | | | 1,777 | | | | | | | | | |
| HEATR3* | | | | | | | 1,810 | | | | | | | | | |
| MARK4* | | | | | | | 1,961 | | | | | | | | | |
| TRIM21* | | | | | | | 1,965 | | | | | | | | | |
| PKIA* | | | | | | | 2,050 | | | | | | | | | |
| HSCB* | | | | | | | 2,057 | | | | | | | | | |
| OPA3* | | | | | | | 2,068 | | | | | | | | | |
| AC006480.2* | | | | | | | 2,083 | | | | | | | | | |
| ZNF552* | | | | | | | 2,125 | | | | | | | | | |
| CPT2* | | | | | | | 2,212 | | | | | | | | | |
| **NOTCH2NLC*** | | | | | | | 2,236 | | | | | | | | | |
| **ARHGAP18*** | | | | | | | 2,577 | | | | | | | | | |
| **ISM1*** | | | | | | | 2,763 | | | | | | | | | |
| **GPR15*** | | | | | | | 2,934 | | | | | | | | | |
| **LYZ*** | | | | | | | 3,138 | | | | | | | | | |
| **RTKN2/*** | | | | | | | -1,560* | 1,953/0,763* | | | | | | | | |
| **TRUB1*** | | | | | | | | -3,682 | | | | | | | | |
| PBX3* | | | | | | | | -3,140 | | | | | | | | |
| ACVR2A* | | | | | | | | -2,968 | | | | | | | | |
| POLRMT* | | | | | | | | -2,645 | | | | | | | | |
| **DDIT3*** | | | | | | | | -2,505 | | | | | | | | |

| GENES | Memory B | Naive B | Naive CD4 | Naive CD8 | MAIT | Th17 | Th1 | Treg | Th2 | Tfh | NK | NKT | CD8 CM | CD8 EM | CD8 EMRA+ | gdTcells |
|---|---|---|---|---|---|---|---|---|---|---|---|---|---|---|---|---|
| COX10* | | | | | | | | -2,465 | | | | | | | | |
| SFR1* | | | | | | | | -2,387 | | | | | | | | |
| WDR75* | | | | | | | | -2,236 | | | | | | | | |
| FLOT1* | | | | | | | | -2,173 | | | | | | | | |
| MRPS28* | | | | | | | | -2,121 | | | | | | | | |
| ARF3* | | | | | | | | -2,078 | | | | | | | | |
| HEIH* | | | | | | | | -2,078 | | | | | | | | |
| HIPK2* | | | | | | | | -2,075 | | | | | | | | |
| VPS39* | | | | | | | | -2,007 | | | | | | | | |
| DDX10* | | | | | | | | -1,986 | | | | | | | | |
| AKAP7* | | | | | | | | -1,906 | | | | | | | | |
| OSBP* | | | | | | | | -1,854 | | | | | | | | |
| SIRT1* | | | | | | | | -1,774 | | | | | | | | |
| CNOT6* | | | | | | | | -1,756 | | | | | | | | |
| SLC9A9* | | | | | | | | -1,751 | | | | | | | | |
| SH3GLB2* | | | | | | | | -1,723 | | | | | | | | |
| VEZF1* | | | | | | | | -1,719 | | | | | | | | |
| MTMR6* | | | | | | | | -1,706 | | | | | | | | |
| TMEM131L* | | | | | | | | -1,664 | | | | | | | | |
| RARS* | | | | | | | | -1,648 | | | | | | | | |
| ADSL* | | | | | | | | -1,620 | | | | | | | | |
| C1orf52* | | | | | | | | -1,543 | | | | | | | | |
| MMP24OS* | | | | | | | | -1,503 | | | | | | | | |
| BLCAP* | | | | | | | | -1,474 | | | | | | | | |
| LSM10* | | | | | | | | -1,308 | | | | | | | | |
| MOAP1* | | | | | | | | -1,259 | | | | | | | | |
| LPIN2* | | | | | | | | -1,245 | | | | | | | | |
| CCR7* | | | | | | | | -1,234 | | | | | | | | |
| MTR* | | | | | | | | -1,217 | | | | | | | | |
| TRABD2A* | | | | | | | | -1,180 | | | | | | | | |
| PIM1* | | | | | | | | -1,035 | | | | | | | | |
| MSN* | | | | | | | | -0,699 | | | | | | | | |
| ZFAS1* | | | | | | | | -0,482 | | | | | | | | |
| ACTG1* | | | | | | | | -0,418 | | | | | | | | |
| HLA-A* | | | | | | | | 0,304 | | | | | | | | |

| GENES | **Memory B** | **Naive B** | **Naive CD4** | **Naive CD8** | **MAIT** | **Th17** | **Th1** | **Treg** | Th2 | Tfh | NK | NKT | CD8 CM | CD8 EM | CD8 EMRA+ | gdTcells |
|---|---|---|---|---|---|---|---|---|---|---|---|---|---|---|---|---|
| SYNE2* | | | | | | | | 0,516 | | | | | | | | |
| **RSF1** | | | | | | | | 0,555 | | | | | | | | |
| MYH9* | | | | | | | | 0,578 | | | | | | | | |
| CD2* | | | | | | | | 0,583 | | | | | | | | |
| UBB* | | | | | | | | 0,598 | | | | | | | | |
| **PAK2** | | | | | | | | 0,659 | | | | | | | | |
| **FKBP1A** | | | | | | | | 0,668 | | | | | | | | |
| TRA2A | | | | | | | | 0,693 | | | | | | | | |
| BPTF | | | | | | | | 0,697 | | | | 1,680 | | | | |
| **HNRNPA2B1** | | | | | | | | 0,746 | | | | | | | 1,307 | |
| TPM3 | | | | | | | | 0,751 | | | | | | | | |
| FUS | | | | | | | | 0,763 | | | | 0,802 | | | | |
| NSA2 | | | | | | | | 0,791 | | | | | | | | |
| RPL4 | | | | | | | | 0,824 | | | | 1,236 | | | | |
| CD7* | | | | | | | | 0,866 | | | | | | | | |
| HERPUD1* | | | | | | | | 0,902 | | | | | | | | |
| FOXP3* | | | | | | | | 0,910 | | | | | | | | |
| SLU7* | | | | | | | | 1,034 | | | | | | | | |
| UBE2K* | | | | | | | | 1,128 | | | | | | | | |
| APH1A* | | | | | | | | 1,135 | | | | | | | | |
| BAX* | | | | | | | | 1,142 | | | | | | | | |
| INPP5D* | | | | | | | | 1,184 | | | | | | | | |
| LONP2* | | | | | | | | 1,244 | | | | | | | | |
| HLA.A | | | | | | | | 1,255 | | | | | | | | |
| KDM5B* | | | | | | | | 1,327 | | | | | | | | |
| **JAK1** | | | | | | | | 1,364 | | | | | | | | |
| FAM120A* | | | | | | | | 1,426 | | | | | | | | |
| ARHGEF1 | | | | | | | | 1,458 | | | | | | | | |
| SLC25A39* | | | | | | | | 1,508 | | | | | | | | |
| NDUFB8 | | | | | | | | 1,574 | | | | | | | | |
| AKR1B1* | | | | | | | | 1,612 | | | | | | | | |
| GLRX | | | | | | | | 1,627 | | | | | | | | |
| FBH1* | | | | | | | | 1,639 | | | | | | | | |
| CNOT6L | | | | | | | | 1,648 | | | | | | | | |
| RPS6KB2* | | | | | | | | 1,720 | | | | | | | | |

| GENES | **Memory B** | **Naive B** | **Naive CD4** | **Naive CD8** | **MAIT** | **Th17** | **Th1** | **Treg** | Th2 | Tfh | NK | NKT | CD8 CM | CD8 EM | CD8 EMRA + | gdTcells |
|---|---|---|---|---|---|---|---|---|---|---|---|---|---|---|---|---|
| TRIM69* | | | | | | | | 1,724 | | | | | | | | |
| BCCIP* | | | | | | | | 1,727 | | | | | | | | |
| MYCBP* | | | | | | | | 1,818 | | | | | | | | |
| RANBP6* | | | | | | | | 1,867 | | | | | | | | |
| XRRA1* | | | | | | | | 1,924 | | | | | | | | |
| RNF170* | | | | | | | | 2,082 | | | | | | | | |
| GPALPP1* | | | | | | | | 2,198 | | | | | | | | |
| **ARRDC1*** | | | | | | | | 2,220 | | | | | | | | |
| MDP1* | | | | | | | | 2,258 | | | | | | | | |
| ZDHHC17* | | | | | | | | 2,275 | | | | | | | | |
| TFPT* | | | | | | | | 2,332 | | | | | | | | |
| SH3GL1* | | | | | | | | 2,551 | | | | | | | | |
| TM2D2* | | | | | | | | 2,592 | | | | | | | | |
| TMCO4* | | | | | | | | 2,790 | | | | | | | | |
| FANCM* | | | | | | | | 2,824 | | | | | | | | |
| **IKZF4*** | | | | | | | | 2,950 | | | | | | | | |
| ABHD17B* | | | | | | | | 2,956 | | | | | | | | |
| AP5B1* | | | | | | | | 2,984 | | | | | | | | |
| BACE1-AS* | | | | | | | | 3,049 | | | | | | | | |
| PUS1* | | | | | | | | 3,073 | | | | | | | | |
| TRBV28* | | | | | | | | 4,282 | | | | | | | | |
| **CLK1/*** | | | | | | | | 0,420/-0,953 | | | | | | | | |
| **OPTN/*** | | | | | | | | 1,803/0,717* | | | | | 1,575 | | | |
| **CDK13/*** | | | | | | | | 1,828/0,831 | | | | | | | | |
| **TRIM22/*** | | | | | | | | 2,158/1,053* | | | | | | | | |
| IRF3 | | | | | | | | | 0,711 | | | | | | | |
| UBE2D2 | | | | | | | | | 0,743 | | | | | | | |
| SPCS1 | | | | | | | | | 1,245 | | | | | | | |
| EIF3F | | | | | | | | | 1,268 | | | | | | | |
| YTHDC1 | | | | | | | | | 1,286 | | | | | | | |
| YWHAZ | | | | | | | | | 1,288 | | | | | | | |

| GENES | **Memory B** | **Naive B** | **Naive CD4** | **Naive CD8** | **MAIT** | **Th17** | **Th1** | **Treg** | Th2 | Tfh | NK | NKT | CD8 CM | CD8 EM | CD8 EMRA + | gdTcells |
|---|---|---|---|---|---|---|---|---|---|---|---|---|---|---|---|---|
| PDIA6 | | | | | | | | | 1,327 | | | | | | | |
| RNASET2 | | | | | | | | | 1,332 | | | | | | | |
| PSMC5 | | | | | | | | | 1,347 | | | | | | | |
| UBE2V1 | | | | | | | | | 1,349 | | | | | | | |
| NDUFB11 | | | | | | | | | 1,389 | | | | | | | |
| JTB | | | | | | | | | 1,454 | | | | | | | |
| KAT6A | | | | | | | | | 1,468 | | | | | | | |
| AKAP9 | | | | | | | | | 1,484 | | | | | | | |
| GYPC | | | | | | | | | 1,490 | | | | | | 1,759 | |
| HP1BP3 | | | | | | | | | 1,538 | | | | 1,366 | | | |
| EVI2B | | | | | | | | | 1,552 | | | | | | | |
| SPTBN1 | | | | | | | | | 1,552 | | | | | | | |
| DOK2 | | | | | | | | | 1,563 | | | | | | | |
| DUT | | | | | | | | | 1,577 | | | | | | | |
| ZMYM2 | | | | | | | | | 1,594 | | | | | | | |
| SFXN1 | | | | | | | | | 1,630 | | | | | | | |
| XRN2 | | | | | | | | | 1,632 | 0,734 | | | | | | |
| PIK3R1 | | | | | | | | | 1,674 | | | | | | | |
| TAF15 | | | | | | | | | 1,827 | | | | | | | |
| DYNLT1 | | | | | | | | | 2,268 | | | | 1,914 | | | |
| MYCBP2 | | | | | | | | | 1,464* | | | | 1,420 | | | |
| EIF5B | | | | | | | | | | 0,607 | | | | | | |
| UBALD2 | | | | | | | | | | 0,707 | | | | | | |
| MDM4 | | | | | | | | | | 0,709 | | | | | | |
| PSMA7 | | | | | | | | | | 0,737 | | | | | | |
| NDUFS5 | | | | | | | | | | 0,742 | | | | | | |
| JPX | | | | | | | | | | 0,752 | | | | | | |
| HCLS1 | | | | | | | | | | 0,753 | | | | | | |
| FKBP8 | | | | | | | | | | 0,753 | | | | | | |
| RPS27L | | | | | | | | | | 0,776 | | | | | | |
| RAD21 | | | | | | | | | | 0,776 | | | | | | |
| NSD3 | | | | | | | | | | 0,783 | | | | | | |
| EIF4A2 | | | | | | | | | | 0,829 | | | | | | |
| GSTK1 | | | | | | | | | | 1,241 | | | | | 1,715 | |
| SMDT1 | | | | | | | | | | 1,311 | | | | | | |

| GENES | **Memory B** | **Naive B** | **Naive CD4** | **Naive CD8** | **MAIT** | **Th17** | **Th1** | **Treg** | Th2 | Tfh | NK | NKT | CD8 CM | CD8 EM | CD8 EMRA+ | gdTcells |
|---|---|---|---|---|---|---|---|---|---|---|---|---|---|---|---|---|
| HLA.F | | | | | | | | | | 1,441 | | 1,461 | 1,372 | | | |
| SF3B2 | | | | | | | | | | 1,553 | | | | | | |
| CD47 | | | | | | | | | | 1,593 | | | | | | |
| PDE3B | | | | | | | | | | 1,896 | | | | | | |
| COX7A2L | | | | | | | | | | | 0,684 | | | | | |
| ROCK1 | | | | | | | | | | | 0,690 | | | | | |
| PSAP | | | | | | | | | | | 0,721 | | | | | |
| ERBIN | | | | | | | | | | | 0,739 | | | | | |
| CFLAR | | | | | | | | | | | 0,777 | | | | | |
| DNAJC8 | | | | | | | | | | | 0,810 | | | | | |
| EIF3K | | | | | | | | | | | 0,836 | | | | | |
| KRTCAP2 | | | | | | | | | | | 0,844 | | | | | |
| MAP2K2 | | | | | | | | | | | 1,180 | | | | | |
| KLRB1 | | | | | | | | | | | | 0,485 | | | | |
| RAB5IF | | | | | | | | | | | | 0,605 | | | | |
| CAPNS1 | | | | | | | | | | | | 0,610 | | | | |
| RHOA | | | | | | | | | | | | 1,254 | | | | |
| PNISR | | | | | | | | | | | | 1,293 | | | | |
| MEAF6 | | | | | | | | | | | | 1,461 | | | | |
| RNASEK | | | | | | | | | | | | 1,537 | | | | |
| BCL11B | | | | | | | | | | | | 1,552 | | | | |
| PCED1B.AS1 | | | | | | | | | | | | 1,582 | | | | |
| TAPBP | | | | | | | | | | | | 1,616 | | | | |
| SCP2 | | | | | | | | | | | | 1,638 | | | | |
| UBXN4 | | | | | | | | | | | | 1,706 | | | | |
| TMED4 | | | | | | | | | | | | 1,768 | | | | |
| CSNK1A1 | | | | | | | | | | | | 1,879 | | | | |
| CREBRF | | | | | | | | | | | | 1,937 | | | | |
| PEBP1 | | | | | | | | | | | | 1,957 | | | | |
| GSTP1 | | | | | | | | | | | | 2,084 | | | | |
| SETX | | | | | | | | | | | | 2,230 | | | | |
| PRDX6 | | | | | | | | | | | | 2,240 | | | | |
| COX6C | | | | | | | | | | | | | 1,156 | | | |
| PDCD4 | | | | | | | | | | | | | 1,243 | | | 0,592 |
| GAPDH | | | | | | | | | | | | | 1,291 | | | |

| GENES | Memory B | Naive B | Naive CD4 | Naive CD8 | MAIT | Th17 | Th1 | Treg | Th2 | Tfh | NK | NKT | CD8 CM | CD8 EM | CD8 EMRA+ | gdTcells |
|---|---|---|---|---|---|---|---|---|---|---|---|---|---|---|---|---|
| ATP5F1D | | | | | | | | | | | | | 1,293 | | | |
| SPCS3 | | | | | | | | | | | | | 1,352 | | | |
| HIGD2A | | | | | | | | | | | | | 1,354 | | | |
| UBE2I | | | | | | | | | | | | | 1,364 | | | |
| SOD1 | | | | | | | | | | | | | 1,382 | | | 2,286 |
| LITAF | | | | | | | | | | | | | 1,389 | | | |
| ARPC2 | | | | | | | | | | | | | 1,390 | | | |
| CCND3 | | | | | | | | | | | | | 1,396 | | | |
| CCDC85B | | | | | | | | | | | | | 1,425 | | | |
| GNB1 | | | | | | | | | | | | | 1,455 | | | |
| RCSD1 | | | | | | | | | | | | | 1,465 | | | |
| GLG1 | | | | | | | | | | | | | 1,488 | | | |
| AC092821.3 | | | | | | | | | | | | | 1,494 | | | |
| TSPO | | | | | | | | | | | | | 1,534 | | | |
| P4HB | | | | | | | | | | | | | 1,551 | | | |
| RBL2 | | | | | | | | | | | | | 1,565 | | | |
| ARHGDIA | | | | | | | | | | | | | 1,569 | | | |
| ATP2B1 | | | | | | | | | | | | | 1,576 | | | |
| XBP1 | | | | | | | | | | | | | 1,577 | | | |
| DOCK8 | | | | | | | | | | | | | 1,614 | | | |
| SH3KBP1 | | | | | | | | | | | | | 1,690 | | | |
| STK10 | | | | | | | | | | | | | 1,691 | | | |
| WDR83OS | | | | | | | | | | | | | 1,807 | | | |
| SNHG7 | | | | | | | | | | | | | 1,907 | | | |
| SLC9A3R1 | | | | | | | | | | | | | 2,359 | | | |
| MIF | | | | | | | | | | | | | | 1,138 | | |
| CBX3 | | | | | | | | | | | | | | 1,299 | | |
| NDUFC2 | | | | | | | | | | | | | | 1,390 | | |
| SNRPG | | | | | | | | | | | | | | 1,449 | | |
| CYTH1 | | | | | | | | | | | | | | | 0,509 | |
| ATP5MD | | | | | | | | | | | | | | | 0,649 | |
| ANKRD11 | | | | | | | | | | | | | | | 0,662 | |
| NDUFA6 | | | | | | | | | | | | | | | 0,723 | |
| RPL10A | | | | | | | | | | | | | | | 1,132 | |
| RPL28 | | | | | | | | | | | | | | | 1,172 | |

| GENES | **Memory B** | **Naive B** | **Naive CD4** | **Naive CD8** | **MAIT** | **Th17** | **Th1** | **Treg** | Th2 | Tfh | NK | NKT | CD8 CM | CD8 EM | CD8 EMRA+ | gdTcells |
|---|---|---|---|---|---|---|---|---|---|---|---|---|---|---|---|---|
| ZEB2 | | | | | | | | | | | | | | | 1,370 | |
| EFHD2 | | | | | | | | | | | | | | | 1,507 | |
| KMT2E | | | | | | | | | | | | | | | 1,574 | |
| CCSER2 | | | | | | | | | | | | | | | 1,774 | |
| RSL24D1 | | | | | | | | | | | | | | | 1,827 | |
| CWC15 | | | | | | | | | | | | | | | 2,118 | |
| ARGLU1 | | | | | | | | | | | | | | | | 0,620 |
| MALAT1 | | | | | | | | | | | | | | | | 0,855 |
| PIK3IP1 | | | | | | | | | | | | | | | | 1,468 |
| U2SURP | | | | | | | | | | | | | | | | 1,883 |
| DBI | | | | | | | | | | | | | | | | 2,153 |

**Table Sup 3. Differentially expressed genes (DEGs) according to two methods in each lymphocyte subsets**.

Genes with * were identified in the DESeq2 analysis and are expressed as Log2(fold-change). Other genes were identified by the global ratio analysis and the value of the global ratio ((M3 vitamin D / D0 vitamin D) / (M3 placebo / D0 placebo)) is reported in the table.

Significant genes from both analyses ($p<0.05$) are quoted ‘gene/*’ and values are reported are as such: global ratio / DESeq2*. Genes in bold were selected for the qPCR validation step.

| | Pathway | pval | padj | qvalue | Fold Change |
|---|---|---|---|---|---|
| Bmemory Vitamin D | POSITIVE_EPIGENETIC_REGULATION_OF_RRNA_EXPRESSION | 6,74E-08 | 6,53E-05 | 2,05 | -0,77 |
| | TRAF6_MEDIATED_NF_KB_ACTIVATION | 6,87E-06 | 0,0067 | 1,48 | -0,76 |
| | INTERLEUKIN_1_FAMILY_SIGNALING | 1,16E-05 | 0,0112 | 1,40 | -1,08 |
| Bnaive Vitamin D | TOLL_LIKE_RECEPTOR_9_TLR9_CASCADE | 3,86E-12 | 3,74E-09 | 2,90 | -1,57 |
| | MAPK_FAMILY_SIGNALING_CASCADES | 3,10E-10 | 3,00E-07 | 2,55 | 0,61 |
| | NGF_STIMULATED_TRANSCRIPTION | 7,08E-10 | 6,86E-07 | 2,48 | -1,18 |
| | SIGNALING_BY_NUCLEAR_RECEPTORS | 3,42E-08 | 3,32E-05 | 2,12 | -2,55 |
| | SENESCENCE_ASSOCIATED_SECRETORY_PHENOTYPE_SASP | 7,06E-08 | 6,84E-05 | 2,04 | -1,14 |
| | SIGNALING_BY_RECEPTOR_TYROSINE_KINASES | 7,06E-08 | 6,84E-05 | 2,04 | -0,22 |
| | INTERLEUKIN_1_SIGNALING | 5,62E-07 | 0,0005 | 1,81 | 1,03 |
| | MYD88_INDEPENDENT_TLR4_CASCADE | 5,62E-07 | 0,0005 | 1,81 | -1,38 |
| | FC_EPSILON_RECEPTOR_FCERI_SIGNALING | 7,83E-07 | 0,0008 | 1,77 | 0,11 |
| | ESTROGEN_DEPENDENT_GENE_EXPRESSION | 2,06E-06 | 0,0020 | 1,64 | -2,44 |
| | RAF_INDEPENDENT_MAPK1_3_ACTIVATION | 2,06E-06 | 0,0020 | 1,64 | -0,81 |
| | TOLL_LIKE_RECEPTOR_TLR1_TLR2_CASCADE | 2,06E-06 | 0,0020 | 1,64 | -1,54 |
| | PRE_NOTCH_EXPRESSION_AND_PROCESSING | 2,82E-06 | 0,0027 | 1,60 | -1,03 |
| | DEFECTIVE_INTRINSIC_PATHWAY_FOR_APOPTOSIS | 5,22E-06 | 0,0051 | 1,52 | -0,33 |
| | DISEASES_OF_SIGNAL_TRANSDUCTION_BY_GROWTH_FACTOR_RECEPTORS_AND_SECOND_MESSENGERS | 1,70E-05 | 0,0165 | 1,34 | 1,78 |
| | ESR_MEDIATED_SIGNALING | 1,70E-05 | 0,0165 | 1,34 | -2,35 |
| | MAPK6_MAPK4_SIGNALING | 1,70E-05 | 0,0165 | 1,34 | 0,45 |
| | LEISHMANIA_INFECTION | 2,27E-05 | 0,0220 | 1,29 | 0,79 |
| | MAPK_TARGETS_NUCLEAR_EVENTS_MEDIATED_BY_MAP_KINASES | 2,27E-05 | 0,0220 | 1,29 | -0,97 |
| | NEGATIVE_REGULATION_OF_MAPK_PATHWAY | 2,27E-05 | 0,0220 | 1,29 | -1,27 |
| | NUCLEAR_EVENTS_KINASE_AND_TRANSCRIPTION_FACTOR_ACTIVATION | 2,27E-05 | 0,0220 | 1,29 | -1,16 |
| | AMINO_ACIDS_REGULATE_MTORC1 | 3,96E-05 | 0,0384 | 1,19 | 0,14 |
| CD4naive Placebo | CELLULAR_RESPONSE_TO_STARVATION | 3,29E-07 | 0,0003 | 1,87 | -0,44 |
| | EUKARYOTIC_TRANSLATION_ELONGATION | 8,79E-07 | 0,0009 | 1,75 | -0,87 |
| | SELENOAMINO_ACID_METABOLISM | 1,42E-06 | 0,0014 | 1,69 | -0,78 |
| | RRNA_PROCESSING | 3,59E-06 | 0,0035 | 1,57 | -0,07 |
| | NONSENSE_MEDIATED_DECAY_NMD | 8,76E-06 | 0,0085 | 1,44 | -0,52 |
| | RESPONSE_OF_EIF2AK4_GCN2_TO_AMINO_ACID_DEFICIENCY | 8,76E-06 | 0,0085 | 1,44 | -0,70 |

| | | | | | |
|---|---|---|---|---|---|
| | DNA_REPLICATION | 2,54E-05 | 0,0246 | 1,27 | 0,84 |
| | INFLUENZA_INFECTION | 4,68E-05 | 0,0454 | 1,16 | -0,38 |
| | SRP_DEPENDENT_COTRANSLATIONAL_PROTEIN_TARGETING_TO_MEMBRANE | 4,68E-05 | 0,0454 | 1,16 | -0,32 |
| CD4naive Vitamin D | TOLL_LIKE_RECEPTOR_9_TLR9_CASCADE | 1,81E-07 | 0,0002 | 1,94 | -1,20 |
| | MAPK_TARGETS_NUCLEAR_EVENTS_MEDIATED_BY_MAP_KINASES | 6,88E-06 | 0,0067 | 1,48 | -0,65 |
| | SIGNALING_BY_ROBO_RECEPTORS | 6,88E-06 | 0,0067 | 1,48 | -1,22 |
| | RESPONSE_OF_EIF2AK4_GCN2_TO_AMINO_ACID_DEFICIENCY | 1,20E-05 | 0,0116 | 1,39 | -1,69 |
| CD8naive Placebo | EUKARYOTIC_TRANSLATION_ELONGATION | 2,31E-06 | 0,0022 | 1,63 | 0,05 |
| | CELLULAR_RESPONSE_TO_STARVATION | 1,13E-05 | 0,0109 | 1,40 | 0,37 |
| | SELENOAMINO_ACID_METABOLISM | 4,89E-05 | 0,0474 | 1,15 | 0,25 |
| MAIT Placebo | TRANSCRIPTIONAL_REGULATION_BY_TP53 | 4,77E-05 | 0,0462 | 1,16 | -0,93 |
| MAIT Vitamin D | TOLL_LIKE_RECEPTOR_TLR1_TLR2_CASCADE | 8,00E-06 | 0,0077 | 1,45 | -1,45 |
| Th17 Placebo | RESPONSE_OF_EIF2AK4_GCN2_TO_AMINO_ACID_DEFICIENCY | 1,33E-05 | 0,0128 | 1,38 | -2,20 |
| Treg Vitamin D | MAP2K_AND_MAPK_ACTIVATION | 3,74E-05 | 0,0363 | 1,20 | 0,92 |

**Table Sup 4. Analysis of pathways regulated by vitamin D revealed by SCPA Package.**

Only pathways with a p-value adjusted <0.05 were represented.
